# Metabolomic profiling of glyphosate resistance evolution in green alga *Chlamydomonas reinhardtii*

**DOI:** 10.1101/2025.07.10.664120

**Authors:** Erika M. Hansson, Heather J. Walker, Dylan Z. Childs, Andrew P. Beckerman

## Abstract

The widespread use of glyphosate has led to a rapid increase in glyphosate resistant weeds at a considerable cost to agriculture. Furthermore, agricultural run-off means non-target, natural ecosystems are also affected. Resistance may arise both through mutations to target enzyme EPSPS in the shikimate pathway generating insensitivity to glyphosate and through non-target site mutations regulating the dose of glyphosate reaching the shikimate pathway. Either mechanism may trade-off against normal cell functioning. Here, using experimental evolution of replicated *Chlamydomonas reinhardtii* populations facing glyphosate, we reveal the metabolomic fingerprint of both glyphosate action and emerging resistance by applying untargeted metabolomic screens throughout the course of resistance evolution. Furthermore, this allows us to evaluate potential underlying molecular mechanisms and fitness costs. We find evidence of build-up of shikimate pathway metabolites, a characteristic signal of glyphosate action, that subsides but does not disappear as resistance evolves, suggesting a fitness trade-off. Concurrent with this is evidence of cell wide disruption of the amino acid pool, that stabilises as resistance evolves. We also found evidence of effects of glyphosate on membrane lipids and increased levels of reactive oxygen species persisting after resistance had evolved. This suggests a considerable effect of the recently described secondary effect of glyphosate: oxidative damage. These data highlight several metabolic processes disrupted by and persisting with the evolution of resistance to glyphosate and may provide a template for enhancing predictions of population and ecosystem effects.

## Introduction

Herbicides are used worldwide to control weeds and ensure high levels of crop production. However, this persistent use constitutes a strong selective pressure for the evolution of herbicide resistant weeds — an increasingly costly problem both ecologically and economically[1, 2]. As resistance emergence is an evolutionary process, evolutionary thinking is necessary both to understand and manage it for the benefit of crop production and biodiversity[3, 4]. This requires characterising the mechanisms by which resistance evolves and the associated fitness costs that may constrain its evolution[5, 6].

There are relatively few approaches that allow the simultaneous evaluation of resistance mechanism and the characterisation of fitness and costs. One of these is the combination of untargeted metabolomics with experimental evolution. Most metabolomic studies target only known compounds in pathways expected to be affected based on the herbicide’s mode of action, introducing a bias that may mischaracterise the full effects of the herbicide. Untargeted metabolomic fingerprinting is a general approach that measures a wider spectrum to give an as comprehensive as possible profile of the metabolome at a given time. This makes it a powerful hypothesis generating tool for identifying the overall metabolomic effects of a herbicide as well as resistance mechanisms and their fitness costs[7, 8, 9]. However, most metabolomics studies of herbicide resistance have focused on comparing resistant to susceptible individuals, missing out on characterising the metabolomic signals of resistance evolution in action. This is afforded by using experimental evolution, which allows controlled insight into the several well-known phases of resistance evolution.

Here we combine experimental evolution and untargeted metabolomic fingerprinting to evaluate the mechanism of resistance and associated fitness dynamics in the green alga *Chlamydomonas reinhardtii* undergoing adaptation to a high dose of glyphosate. *C. reinhardtii* has previously been used as a model system to study efficacy of resistance management strategies[10, 11], characterising fitness costs[12], comparisons of evolutionary potential in communities of algae[13] and molecular analysis of herbicide resistance mutations[14, 15, 16, 17, 18, 19, 20, 21]. Furthermore, while not the primary target of commercial herbicide application, algae are still affected through agricultural runoff[**?**, reviewed in]]VanBruggen2018 and as such the molecular and metabolomic dynamics underlying their adaptation to herbicides has relevance to the functioning of freshwater ecosystems.

Glyphosate is currently the world’s most commonly used herbicide, and the overreliance on glyphosate and genetically engineered glyphosate resistant crops has resulted in a rapid increase in both glyphosate resistant non-crop species and unique glyphosate resistance mechanisms[1, 22]. Glyphosate blocks the shikimate pathway, which is responsible for 30% of the carbon flow in plants, algae and bacteria[23, 24]. By competitively binding to enzyme 5-enolpyruvulshikimate-3-phosphate (EPSPS) instead of intended substrate phosphoenolpyruvate (PEP), production of the aromatic amino acids phenylalanine (Phe), tyrosine (Tyr) and tryptophan (Trp) as well as other downstream products of chorismate metabolism is disrupted, resulting in arrested cell growth and eventual death. Secondary effects include disruption of photosynthesis and increased production of reactive oxygen species (ROS)[25, 26, 27, 28, 29, 30].

There are two primary ways resistance arises to glyphosate. Firstly, single point mutations to EPSPS generally confer low resistance and impair normal enzyme function by lowering affinity for PEP[31, 32, 33, 34, 35, 36, 37]. While there are notable exceptions conferring higher resistance in specific species[33, 38], including sequential double[39, 40, 41] and triple[42, 43] mutations that render EPSPS highly insensitive to glyphosate, the fitness consequences for most of these have yet to be tested. EPSPS amplification has been detected both in the form of upregulated expression[44] and gene multiplication[45, 46], with varied underlying genetic mechanisms[47, 48, 49, 50].

Secondly, glyphosate resistance can also be conferred through adaptations to limit the dose that reaches the target through reduced absorption[51, 52, 53] and increased vacuolar sequestration[54, 55, 56, 57, 58]. Furthermore, glyphosate is metabolised by glyphosate oxidoreductase in bacteria and aldo-keto reductase in at least one plant species[59, 60], and the main metabolic product aminomethylphosphonic acid (AMPA) is commonly detected in plants even when no pathway for catabolism has been identified[61, 62].

We focus on three targeted hypotheses relating to specific compounds selected *a priori* that are expected to change in abundance depending on whether resistance has evolved and by what mechanism: shikimate pathway functioning, glyphosate and its breakdown products, and amino acid (AA) pool composition (Figure 1). We review the rationale for these in the following sections. Further exploratory analysis is used to identify where there are the largest differences in the metabolite profiles between glyphosate-treated and control populations to extend the fingerprint beyond the targeted compounds and identify other cellular processes affected by either glyphosate or glyphosate resistance.

### Predicted metabolic fingerprints of glyphosate resistance evolution

Firstly, to conclude glyphosate resistance has evolved we must first see evidence of glyphosate inhibition of the shikimate pathway presenting as a build-up of upstream pathway metabolites followed by a return to pathway functioning with lower levels of upstream metabolites[63, 64, 65, 66, 67, 68, 69, 25, 70, 71, 72, Figure 1A]. This gives us a timeline of resistance evolving.

Secondly, the mechanism fingerprint is primarily dependent on the presence or absence of glyphosate and its catabolites. Increased levels of intracellular glyphosate post-resistance is consistent with it being sequestered to the vacuole or that insensitive EPSPS is no longer binding it[1, 54, 55, 56], whereas decreased levels post-resistance could indicate decreased absorption into the cell[1] or possibly amplified EPSPS if the applied glyphosate dose is very high (Figure 1B). Presence of known primary breakdown products of glyphosate like AMPA, sarcosine and glyoxylate after resistance has evolved is evidence for it being at least partly conferred by glyphosate catabolism[73, 74, 75, 76], whereas their absence is evidence against (Figure 1B). Out of the four known pathways for glyphosate metabolism, those by glyphosate oxireductase and glyphosate oxidase result only in AMPA and glyoxylate by cleavage of the glyphosate C-N bond[77, 78]. Glyphosate degradation by cleavage of the glyphosate C-P bond by a C-P lyase has several intermediate steps with phosphate and sarcosine being the final product[73]. Similarly, the complete break-down of AMPA results in phosphate after a number of intermediate metabolites[73]. Glyphosate degradation by aldo-keto reductase results in AMPA and glyoxylate, but is associated with possible breakdown of glyoxylate to glycine and 2-oxoglutarate as well as involvement of cinnamaldehyde and cinnamyl alcohol[60]. As such, if the primary glyphosate degradation products are detected, levels of secondary breakdown products can distinguish between known pathways.

Thirdly, trade-offs with normal cell function is determined by upstream shikimate pathway metabolites and the AA pool size and composition. Upstream metabolite levels post-resistance indicates both the effectiveness of resistance and potential fitness trade-offs, with continued elevated levels suggesting some remaining inhibiting effect (Figure 1C) or mutated EPSPS with reduced affinity for PEP (Figure 1B). Perturbation of the AA pool is expected during glyphosate inhibition, but should disappear after resistance has evolved if it is not associated with a fitness cost (Figure 1C). Any remaining effects is indicative of a trade-off between the resistance mechanism and normal cell function. While changes to the abundance of the aromatic AAs Phe, Trp and Tyr is another direct test of shikimate pathway functioning, the effects on the total AA pool and its composition is indicative of wider cell functioning as AAs play multiple, integral roles in cell metabolism beyond protein synthesis, including signalling, regulation and stress response[24, 63, 79, 80, 81, 82, 83, 84]. The effect of glyphosate inhibition on the AA pool differs depending on system, with evidence of both decreases in abundance reflective of widespread disruption to carbon metabolism[65, 64], and increases of specific AAs and the overall pool[85, 63, 86, 87, 25, 71] due to increased catabolism as a stress response[63] or by shifts in production between pathways[86].

## Methods

### Experimental design

#### Algal strain and chemostat setup

*Chlamydomonas reinhardtii* strain Sager’s CC-1690 wild-type 21 gr was obtained from the *Chlamydomonas* Resource Centre (University of Minnesota, St Paul, MN, USA) core collection. Prior to the experiment, the algae had been kept in static stock cultures for two years (Ebert algal medium, 25°C, 24h light), with batches being transferred to fresh medium fortnightly, then in continuous flow culture for two months under the experiment control conditions detailed below.

**Figure 1:**
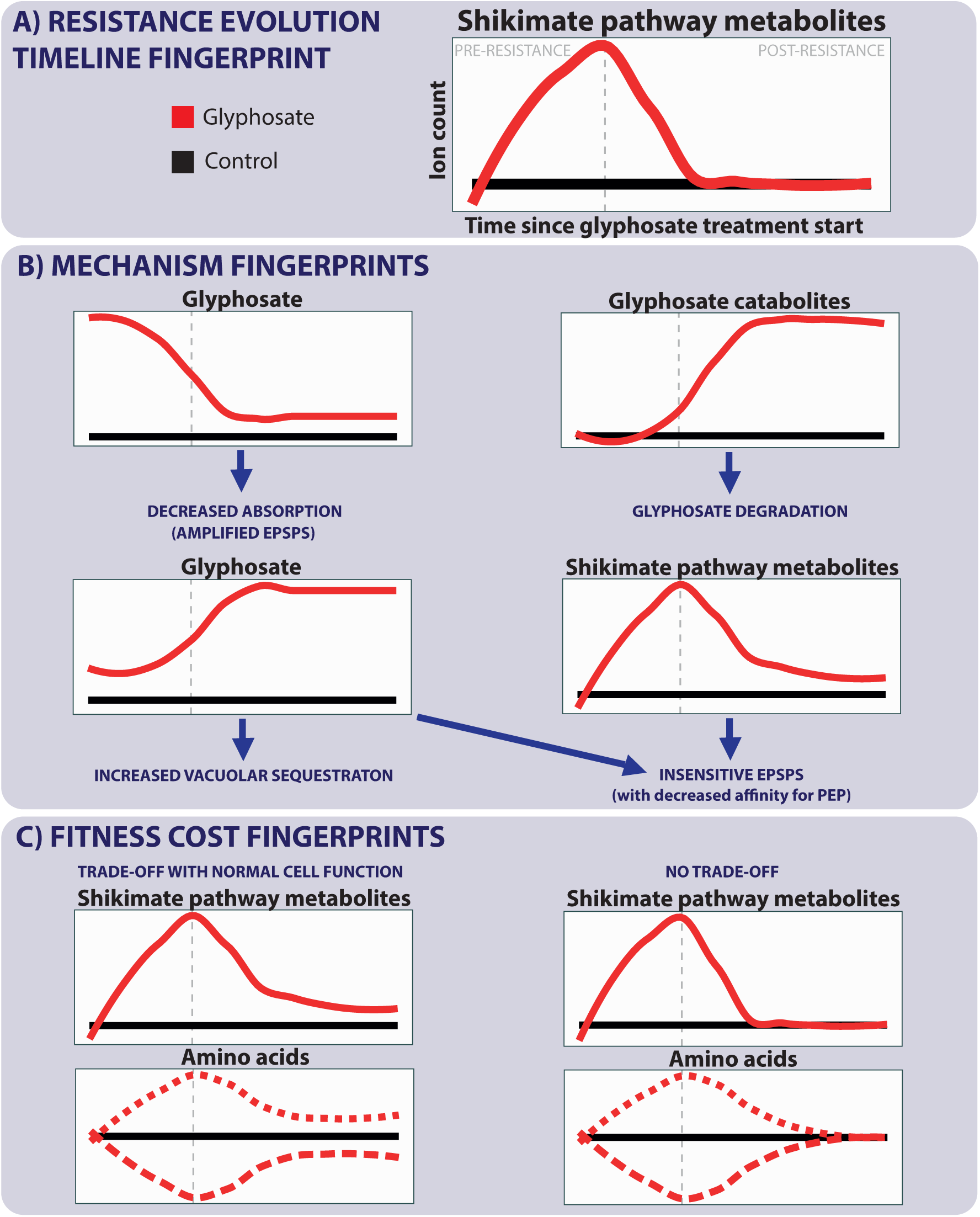
Overview of possible fingerprint outcomes for the three targeted hypotheses. All graphs show ion count vs time since glyphosate treatment start with the glyphosate-treated line in red and the control line in black. The vertical grey dashed line indicates the pre- and post-resistance phase boundary. A) The timeline of glyphosate resistance evolution is based on the shikimate pathway metabolites, the build-up and return to control levels of which define the pre- and post-resistance phases. B) Resistance mechanism fingerprints depend primarily on presence or absence of glyphosate and its catabolites. C) Trade-offs with normal cell function are determined by whether shikimate pathway metabolites and AA levels return to control levels or remain changed after resistance has evolved.

Ten *C. reinhardtii* populations were cultured in a continuous flow through chemostats in sterile Ebert algal medium[88] at a dilution rate of 0.15/24h with a shared multichannel peristaltic pump (Watson-Marlow 205S/CA16) controlling the flow and maintaining a volume of 380ml±5%[89, for detailed protocol]. The cultures were kept on a light box providing white light from below and surrounded on all sides with white light fluorescent bulbs providing a continuous light level of 75 µmol m^-2^ s^-1^. The internal temperature of the cultures was 30°C, heated by the light box and the controlled temperature room they were kept in, and they were continuously mixed by bubbling. The *C. reinhardtii* populations reached steady state in control medium to ensure equal starting population sizes of approximately 250 000 cells/ml before herbicide treatments were introduced 14 days after initial inoculation.

#### Herbicide concentrations and replication

Five culture chambers were exposed to herbicide treatment consisting of glyphosate (PESTANAL®, analytical standard) at 100 mg/L, a concentration completely inhibiting population growth over seven days in batch cultures[10, 11]. Five chambers received control medium. The glyphosate was introduced to each population through direct injection of 38 mg glyphosate in 38 ml of sterile Ebert medium. The control chambers received 38 ml of sterile Ebert medium. Immediately before the treatment injection, 38 ml of the existing culture in the chamber was removed (10% of the total culture volume) to ensure the culture volume remained consistent.

### Cell sample extractions

Algal samples for metabolomic analysis were removed at eight time points throughout the experiment: a sample of the founding population, a sample three days before the glyphosate treatment was applied and 1, 8, 16, 22, 29 and 36 days after glyphosate introduction. The cells were washed with sterile Ebert medium and centrifuged to produce a pellet of cells which was flash frozen with LN2 before storing in -80°C. The frozen algal pellets were added to 2.5 µl 80% ethanol/mg of cells and crushed with a ball bearing in a homogeniser at 6 m/s for 2 minutes. The samples were centrifuged at 14 kRPM for 15 minutes and 7.5 µl of the resulting supernatant was added to 30 µl 80% ethanol.

### Metabolomic profiling using mass spectrometry

Direct injection electrospray ionisation (negative and positive modes) mass spectrometry was applied to the samples and analysed on a Waters Synapt G2-Si ToF mass spectrometer (Waters Corporation, United States). Injections were performed by a Waters Acquity UPLC (Waters Corporation, United States) and the MassLynx data system (Waters Corporation, United States) was used for instrument control and data acquisition. Each biological replicate (n=5 glyphosate, n=5 controls) was evaluated in triplicate (technical replicates) and injected at a flow rate of 5 µl/min for 3 minutes. All spectra were measured from 50 to 1200 Da for both modes with a scan time of 1 scan/second. The conditions for sample introduction are summarised in Table 4.

### Data pre-processing

Following the workflow outlined in Parker (2023)[90], raw data files were converted to open format .mzML using Proteowizard[91]. The MALDIquant[92] and MSnbase[93] packages for R were then used to extract peak tables, remove noise and smooth peaks that were averaged across technical replicates, filtering out any peaks that were not present in at least two technical replicates.

### Targeted analysis: shikimate pathway compounds, glyphosate, glyphosate breakdown products and AAs

All data were analysed in R (version 4.0.5,[94]). The monoisotopic masses for relevant compounds related to the shikimate pathway, known glyphosate breakdown products as well as all AAs were obtained from MetaCyc.org database[95]. Their expected *m/z* peaks (i.e. [M+H]^+^ or [M+H]^-^) putatively matched to the detected peaks in negative and/or positive mode dependent on the compound’s likelihood of ionising under the given conditions. Compounds were accepted as a putative match if the error margin was less than 100 Δppm Da. In order to get a measure of the total AA pool abundance, the percentage ion counts for all peaks matched to an AA were summed for each population by sampling day.

The percentage total ion counts of the matched compound peaks where then compared between the controls and the glyphosate-treated populations throughout the experiment by fitting mixed-effects models with each of the putative peak percentage ion counts as the response (arcsine transformed), treatment (glyphosate or control) and day (as a factor) as well as their interaction as fixed effects and chamber as a random effect to account for repeated measurements using the lme4 package[96]. The emmeans package[97] was used to carry out pairwise comparisons between treatments within each sampling day. The car package function Anova() was used to test the significance of the fixed effects and confirmed through parametric boot-strapping using the pbkrtest package[98]. When the variance of the random effect was zero or near zero for some models, resulting in a singular fit or a convergence failure, these were refit as a linear model without the random effect to validate the robustness of the significance testing.

#### Metabolite identification using tandem mass spectrometry

To validate the presence of the putatively identified compounds, tandem mass spectrometry (MS/MS) using direct injection at 10 µl/min for 1 min on the same system as above was applied targeting the range of each detected mass peak. Fragmentation patterns were resolved against MassBank spectra[99] where available. See Table 5 for instrument settings, MassBank record IDs and matched peaks for each compound.

As the signal for some masses was very low, compounds were accepted as present in the sample as long as the reference spectrum peak with the highest intensity was detected. Compounds where MS/MS fragmentation patterns did not resolve against MassBank spectra or no reference spectra was available were still included as their presence could not be conclusively ruled out. The complete list of compounds used to test each targeted hypothesis is found in Table 1.

### Exploratory analysis to extend the metabolomic profile of evolving resistance

Finally, we located the largest differences in the data set between the glyphosate exposed and control populations to identify other cell machinery changes that may be evidence of costs, trade-offs or other aspects of resistance, such as shifting prioritisation between metabolic pathways or mechanisms specifically targeting the secondary effects of glyphosate. For this extension of the metabolomic fingerprint, the SIMCA (Umetrics, Umeå, Sweden) statistical package was used to carry out Principle Component Analyses (PCA) and supervised multivariate analyses using Orthogonal Partial Least Squares Discriminant Analysis (OPLS-DA). OPLS-DA is used to determine which variables have the largest power to discriminate (loading) between defined groups (explanatory variable).

We applied OPLS-DA models on the whole data set with the explanatory variable as either sampling day or glyphosate treatment, as well as glyphosate treatment as the explanatory variable to the data set subset by sampling day. For each model the 40 most discriminatory *m/z* peaks (i.e. with the largest fold change compared to control) were extracted to create a list of peaks of interest. *m/z* peaks were discarded from the list if: the mean loading was smaller than its standard error, no significant pairwise difference between treatments was found or a significant pair-wise difference between strains was found before the treatment start. Mixed effects models were then fit as above for each remaining peak of interest and the masses putatively matched to compounds using the MetaCyc.org *Chlamydomonas reinhardtii* database for [M-H]^-^, [M-2H]^2-^, [M-3H]^3-^ and [M-H_2_O-H]^-^ ion adducts in negative mode and [M+H]^+^,[M+Na]^+^, [M+K]^+^, [M+2H]^2+^, [M+3H]^3+^, [M+H-H_2_O]^+^ and [M+H-2H_2_O]^+^ ion adducts in positive mode, applying a maximum error margin of 60 Δppm Da.

Lastly, MS/MS analysis was carried out as above for any putatively identified compounds where MassBank reference spectra were available Table 5 and putative compound identities where the fragmentation pattern did not match reference spectra were discarded from further analysis.

**Table 1:** Compounds tested in targeted analysis along with chemical structure, metabolomic role and monoisotopic mass. Note that some compounds have the same monoisotopic mass and thus cannot be distinguished by the metabolite screen carried out here and have been listed together if part of the same hypothesis, and labelled with matching asterisks if part of separate hypotheses. The presence of compounds in *italics* could not be confirmed via MSMS due to absence of reference spectra.

| Compound | Formula | Accurate mass |
| --- | --- | --- |
| ***Hypothesis: Shikimate pathway functioning*** |  |  |
| D-Erythrose-4P (E4P) | C<sub>4</sub>H<sub>7</sub>O<sub>7</sub>P | 200.0086 |
| Phosphoenolpyruvate (PEP) | C<sub>3</sub>H<sub>2</sub>O<sub>6</sub>P | 167.9824 |
| *3-Deoxy-D-Arabino-Heptulosonate 7-Phosphate (DAHP)* | C<sub>7</sub>H<sub>13</sub>O<sub>10</sub>P | 288.0246 |
| *3-Dehydroquininate (3-DHQ)* | C<sub>7</sub>H<sub>9</sub>O<sub>6</sub> | 190.0477 |
| *3-Dehydroshikimate* | C<sub>7</sub>H<sub>7</sub>O<sub>5</sub> | 172.0372 |
| Shikimate | C<sub>7</sub>H<sub>9</sub>O<sub>5</sub> | 174.0528 |
| *Shikimate-3-Phosphate* | C<sub>7</sub>H<sub>8</sub>O<sub>8</sub>P | 254.0192 |
| *5-Enolpyruvoyl-Shikimate 3-Phosphate (EPSP)* | C<sub>10</sub>H<sub>9</sub>O<sub>10</sub>P | 324.0246 |
| *Chorismate/Prephenate* | C<sub>10</sub>H<sub>8</sub>O<sub>6</sub> | 226.0477 |
| ***Hypothesis: Glyphosate and its catabolites*** |  |  |
| Glyphosate | C<sub>3</sub>H<sub>6</sub>NO<sub>5</sub>P | 169.0140 |
| *Aminomethyl-phosphonate (AMPA)* | CH<sub>5</sub>NO<sub>3</sub>P | 111.0085 |
| Sarcosine* | C<sub>3</sub>H<sub>7</sub>NO<sub>2</sub> | 89.0477 |
| Phosphate | HO<sub>4</sub>P | 97.9769 |
| Glyoxylate | C<sub>2</sub>HO<sub>3</sub> | 74.0004 |
| *α-D-ribose-1-[N-(phosphonomethyl)glycine] 5-triphosphate* | C<sub>8</sub>H<sub>13</sub>NO<sub>18</sub>P<sub>4</sub> | 540.9553 |
| *α-D-ribose-1-[N-(phosphonomethyl)glycine] 5-phosphate* | C<sub>8</sub>H<sub>13</sub>NO<sub>12</sub>P<sub>2</sub> | 381.0226 |
| *α-D-ribose 1,5-bisphosphate* | C<sub>5</sub>H<sub>8</sub>O<sub>11</sub>P<sub>2</sub> | 309.9855 |
| *5-phospho-α-D-ribose 1,2-cyclic phosphate* | C<sub>5</sub>H<sub>7</sub>O<sub>10</sub>P<sub>2</sub> | 291.9749 |
| *5-phospho-α-D-ribose 1-diphosphate* | C<sub>5</sub>H<sub>8</sub>O<sub>14</sub>P<sub>3</sub> | 389.9518 |
| *α-D-ribose 1-(acetamidomethylphosphonate) 5-triphosphate* | C<sub>8</sub>H<sub>14</sub>NO<sub>17</sub>P<sub>4</sub> | 524.9603 |
| *α-D-ribose-1-(2-N-acetamidomethylphosphonate) 5-phosphate* | C<sub>8</sub>H<sub>14</sub>NO<sub>11</sub>P<sub>2</sub> | 365.0277 |
| *Cinnamaldehyde* | C<sub>9</sub>H<sub>8</sub>O | 132.0575 |
| *Cinnamyl alcohol* | C<sub>9</sub>H<sub>10</sub>O | 134.0732 |
| *2-oxoglutarate* | C<sub>5</sub>H<sub>4</sub>O<sub>5</sub> | 146.0215 |
| ***Hypothesis: AA pool composition*** |  |  |
| Alanine (Ala)* | C<sub>3</sub>H<sub>7</sub>NO<sub>2</sub> | 89.0477 |
| Arginine (Arg) | C<sub>6</sub>H<sub>14</sub>N<sub>4</sub>O<sub>2</sub> | 174.1117 |
| Asparagine (Asn) | C<sub>4</sub>H<sub>8</sub>N<sub>2</sub>O<sub>3</sub> | 132.0535 |
| Aspartate (Asp) | C<sub>4</sub>H<sub>7</sub>NO<sub>4</sub> | 133.0375 |
| Cysteine (Cys) | C<sub>3</sub>H<sub>7</sub>NO<sub>2</sub>S | 121.0197 |
| Glutamate (Glu) | C<sub>5</sub>H<sub>9</sub>NO<sub>4</sub> | 147.0532 |
| Glutamine (Gln) | C<sub>5</sub>H<sub>10</sub>N<sub>2</sub>O<sub>3</sub> | 146.0691 |
| Glycine (Gly) | C<sub>2</sub>H<sub>5</sub>NO<sub>2</sub> | 75.0320 |
| Histidine (His) | C<sub>6</sub>H<sub>9</sub>N<sub>3</sub>O<sub>2</sub> | 155.0695 |
| Isoleucine (Ile)/Leucine (Leu) | C<sub>6</sub>H<sub>13</sub>NO<sub>2</sub> | 131.0946 |
| Lysine (Lys) | C<sub>6</sub>H<sub>14</sub>N<sub>2</sub>O<sub>2</sub> | 146.1055 |
| Methionine (Met) | C<sub>5</sub>H<sub>11</sub>NO<sub>2</sub>S | 149.0510 |
| Phenylalanine (Phe) | C<sub>9</sub>H<sub>11</sub>NO<sub>2</sub> | 165.0790 |
| Proline (Pro) | C<sub>5</sub>H<sub>9</sub>NO<sub>2</sub> | 115.0633 |
| Serine (Ser) | C<sub>3</sub>H<sub>7</sub>NO<sub>3</sub> | 105.0426 |
| Threonine (Thr) | C<sub>4</sub>H<sub>9</sub>NO<sub>3</sub> | 119.0582 |
| Tryptophan (Trp) | C<sub>11</sub>H<sub>12</sub>N<sub>2</sub>O<sub>2</sub> | 204.0899 |
| Tyrosine (Tyr) | C<sub>9</sub>H<sub>11</sub>NO<sub>3</sub> | 181.0739 |
| Valine (Val) | C<sub>5</sub>H<sub>11</sub>NO<sub>2</sub> | 117.0790 |

Pairwise comparisons between treatments for each sampling day were carried out as above and the compounds were classified according to when significant differences in ion count were observed. Compounds where there was no difference between treatments by the last time point were classified as pre-resistance phase. These are compounds likely involved in initial stress response, build-up or dearth of certain compounds due to the blocked shikimate pathway as well as compounds related to cellular death and breakdown. Sustained differences in compound presence emerging post-resistance is however likely related to the resistance mechanism, including compounds that are up- or down-regulated as part of resource allocation for resistance, or may be due to later effects of chronic glyphosate exposure. Differences throughout the treatment suggests the compound is related to glyphosate action rather than resistance, such as stress responses to secondary effects. As such, compounds where differences were still apparent by the last time point were classified as post-resistance phase if the differences emerged with resistance evolving, whereas those where differences were present from the application of the treatment were classified as persistent differences. The MetaboAnalyst Pathway Analysis tool[100] was used to identify pathways linking the putatively identified compounds within each classification.

## Results

### Shikimate pathway metabolite build-up suggests resistance trait becomes dominant in the population between 16 and 22 days after exposure

We found strong evidence of disrupted shikimate pathway functioning with considerable initial metabolite build-up, as well as clear indication of when resistance evolved as this build-up subsided, with relatively low inter-population variation. Glyphosate treatment and its interaction with day had a strong effect on the ion counts in the *m/z* peaks matched to the three metabolites directly upstream of EPSPS (Figure 2, Table 6). Blockage of the shikimate pathway by glyphosate action is apparent at 8 days after the introduction of the treatment and resulting in considerable build-up of 3-dehydroshikimate (Figure 2c, t-ratio=-21.3, DF=56, p<0.0001), shikimate (Figure 2d, t-ratio=-28.8, DF=56, p<0.0001) and shikimate-3-phosphate (Figure 2e, t-ratio=17.7, DF=51.6, p<0.0001) compared to the control populations, with a 2.4-fold increase in 3-dehydroshikimate, 3.8-fold increase in shikimate and a 4.3-fold increase in shikimate-3-phosphate ion count percentage. While this is the peak for 3-dehydroshikimate and shikimate after which the ion counts stabilise at elevated levels compared to the controls, shikimate-3-phosphate reaches a peak after 16 days with a 4.7-fold increase with subsequent decrease for each week. This suggests the resistance trait becomes dominant in the populations between 16 and 22 days after exposure to glyphosate.

While no *m/z* peaks could be matched to PEP or DHQ, glyphosate treatment had a significant effect (either as a main effect or through its interaction with day, Table 6) on those matched to two targeted metabolites upstream of 3-dehydroshikimate – E4P and DAHP – as well as the two compounds downstream of EPSPS – EPSP and chorismate/prephenate. For DAHP and chorismate/prephenate the pairwise comparisons show relatively small sustained decreases (Figure 2b, DAHP, day 36, t-ratio=-2.4, DF=53.8, p=0.02) and increases (Figure 2g, chorismate/prephenate, day 36, t-ratio=-3.2, DF=51.6, p=0.002) in ion count percentage after the glyphosate treatment was started, but without a clear peak. While there are large fluctuations in the ion count for E4P, this pattern is likely unrelated to the resistance trait as the differences are also present at day -3 (Figure 2a, t-ratio=5.3, DF=31, p<0.0001), before the glyphosate treatment started. EPSP shows minor differences 1 day after treatment start (Figure 2f, t-ratio=2.2, DF=56, p=0.03) and on day 29 (t-ratio=3.3, DF=56, p=0.002), but no difference was found on day 36 (t-ratio=1.4, DF=56, p=0.18).

### No evidence of the free glyphosate in the treated cells

No *m/z* peak could be matched to free glyphosate, indicating that it is either wholly absent from the treated cells or that any glyphosate present is either bound to EPSPS or other enzymes.

### No evidence of glyphosate degradation in resistant populations

Our data suggests no changes in glyphosate catabolite ion count associated with evolved glyphosate resistance. Firstly, no peak could be matched to glyoxylate. Secondly, while a significant effect of either glyphosate treatment as a main effect or its interaction with day was found on the *m/z* peaks putatively matched to several primary and secondary glyphosate degradation products (Table 7), none of the compounds exhibit the expected pattern of an increase to a level that is then sustained after resistance evolves. For the primary catabolites (Figure 3), both phosphate and sarcosine show clear peaks before resistance evolves before stabilising at near-control or control level respectively. Phosphate also shows differences in ion count on day -3 for negative mode, suggesting its fluctuations are likely to be unrelated to glyphosate resistance. While AMPA is slightly higher on day 36 (t-ratio=-2.7, DF=51, p=0.01), this is the only point post-resistance where it is elevated and higher ion counts are otherwise only found pre-resistance (Table 7), with an overall pattern where both controls and treated populations see an increase in ion counts over time. Most secondary compounds show patterns where the effect is in the wrong direction (i.e. lower ion counts than the controls) and/or there is a clear peak or trough in ion count after which the differences between treated populations and controls either are small or disappear entirely (Table 7). Particularly the latter suggests that the changes in the ion counts of these compounds are a signal of cellular breakdown while most of the population is not resistant, and that the cells return to normal or near-normal functioning with regards to these compounds after resistance evolves. Furthermore, this strongly suggests glyphosate degradation does not have a role in the resistance mechanism in these populations.

**Figure 2:**
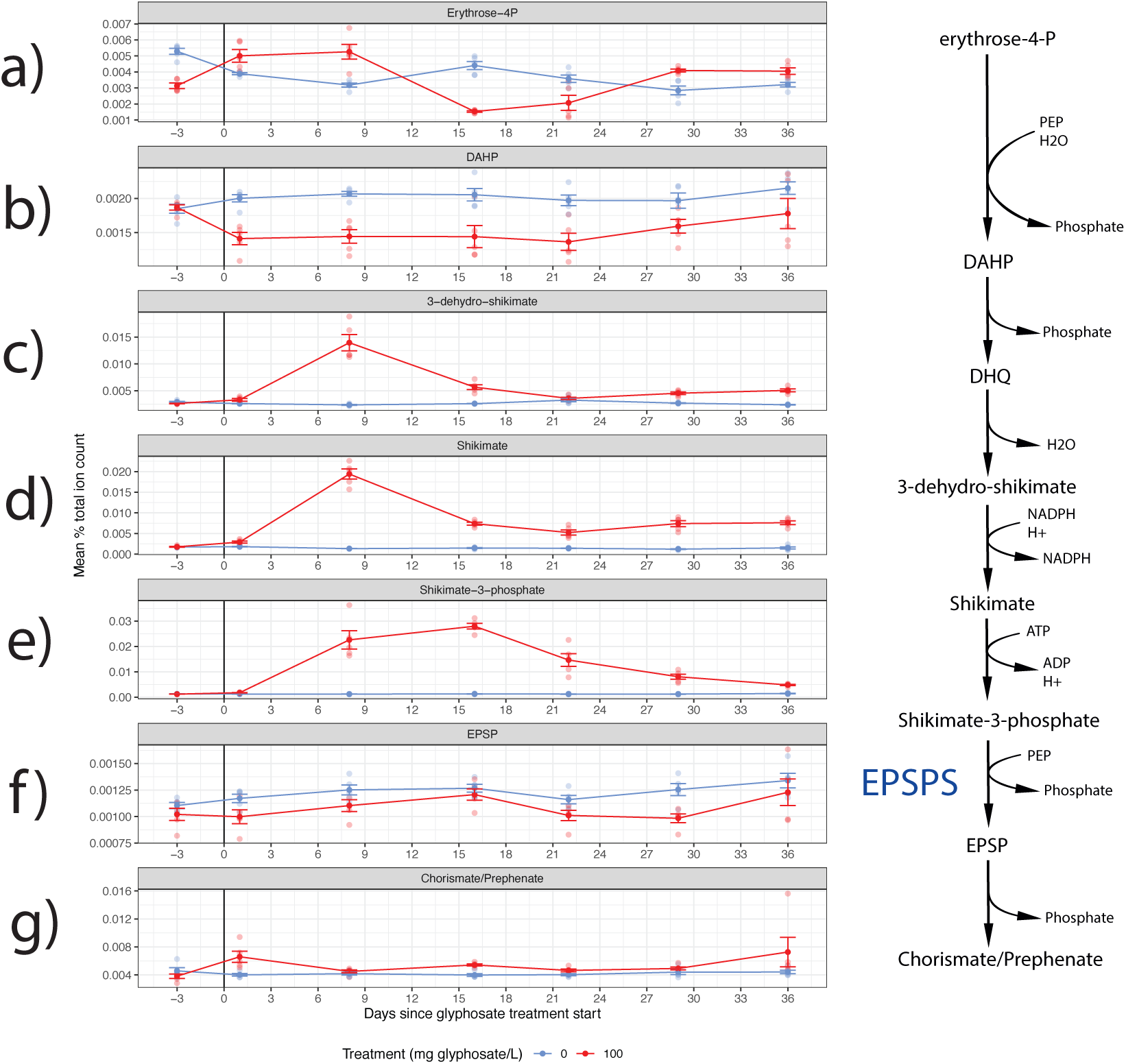
Mean % total ion count with standard error for the putative *m/z* peak matches of compounds in shikimate pathway for control and glyphosate-treated populations throughout the experiment. The compounds are ordered according to their position in the shikimate pathway, but note that PEP appears twice as a substrate. The black vertical line in the graphs indicates start of the glyphosate treatment.

**Figure 3:**
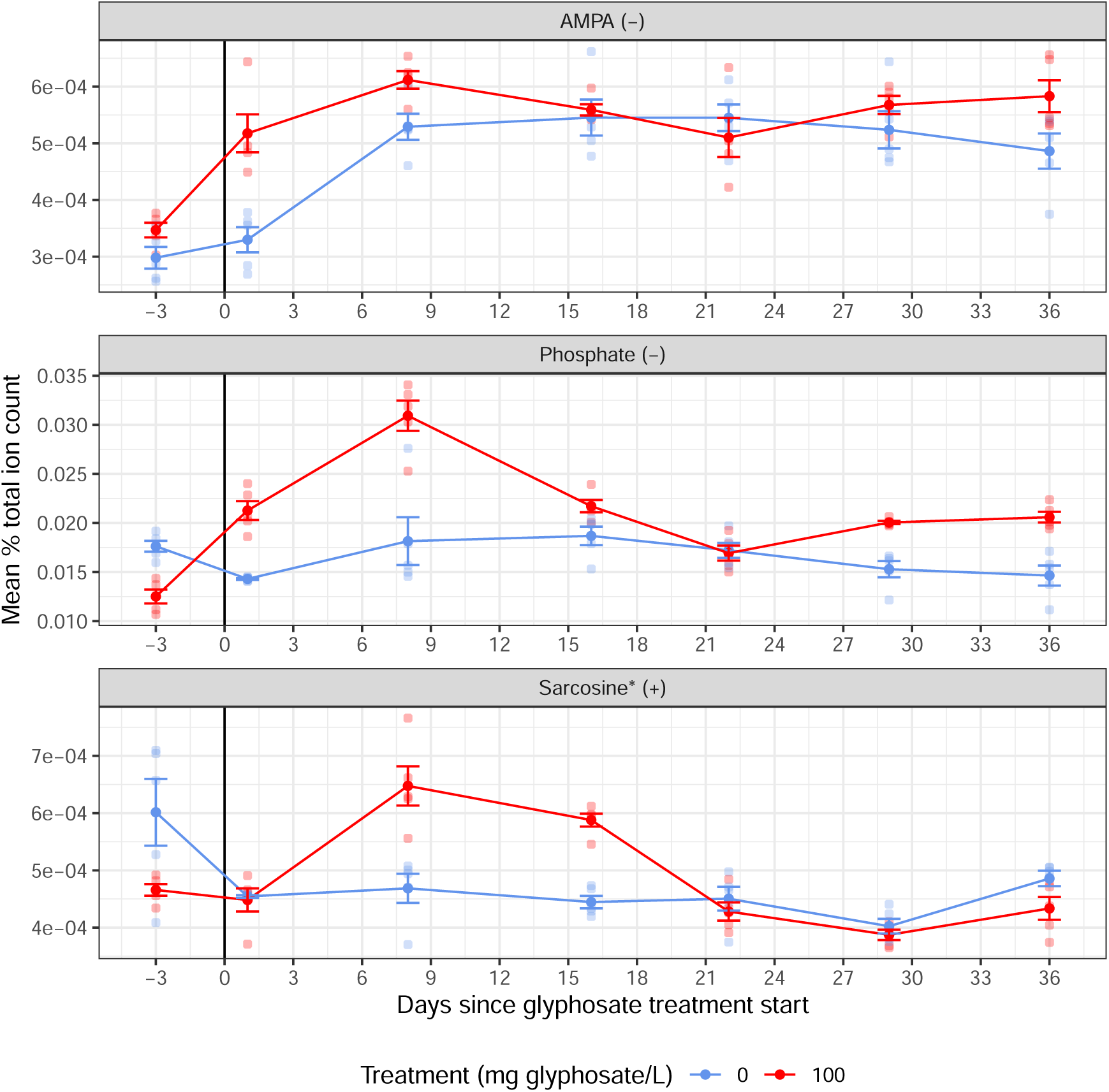
Mean % ion counts with standard error for *m/z* peaks matched to primary glyphosate degradation products in negative and positive mode. The raw data is shown in the background as transparent points and the black vertical line indicates introduction of the glyphosate treatment.

### Shikimate pathway metabolite build-up suggests a possible trade-off between normal EPSPS function and the resistance trait

While the build-up of the three compounds directly upstream of EPSPS subsides with resistance evolving, all three remain at elevated levels compared to the controls equalling a 1.5-(3-dehydroshikimate), 2.3-(shikimate) and 1.8-fold (shikimate-3-phosphate) increases respectively 36 days after the glyphosate treatment was initiated (Figure 2c–f, Table 6). This suggests continued shikimate pathway inhibition and a possible trade-off between normal EPSPS function and the resistance trait. Furthermore, while DAHP and chorismate/prephenate do not show a clear peak, the sustained strain differences after treatment commenced further support the idea that the evolved resistance does not return full shikimate pathway function.

### Changes in AA pool associated with glyphosate treatment, as well as resistance

While the exact percentage ion count pattern through time depends on the AA, a majority showed a distinct “peak” or “trough” pattern compared to the controls that then returns to the same or near the level of the controls, matching the timing of glyphosate resistance evolving as seen in the shikimate pathway metabolites. The effect of glyphosate treatment on the ion counts of the mass peaks matched to AAs depends on compound, the mode and the day (summarised in Table 2, Table 8). However, while a large number of matched *m/z* peaks showed significant pair-wise differences, the largest differences are concentrated around the peak of evolving resistance (days 8–22). Asp, Cys, Glu and Gln showed the largest differences, at some point exceeding a 30% increase or decrease, with the ion count for the *m/z* peak matched to Cys in negative mode increasing more than 3-fold on day 16. The effect of glyphosate treatment during this period is also reflected in the overall ion counts for the total AA pool, but the differences are small in both negative and positive mode, and in positive mode a pairwise difference is also found on day -3. This suggests that mainly composition, more than overall AA pool size, is affected by glyphosate action and resistance evolution.

Asn, Met, Ser, Thr and Lys showed sustained minor pair-wise differences by the end of the experiment, possibly indicating a trade-off between cell functioning and resistance consistent with the pattern seen for the shikimate pathway metabolites. While His, Trp, Val also showed sustained differences, pair-wise differences were also found on day -3, suggesting processes other than resistance evolution may be involved.

### The extended metabolomic fingerprint of glyphosate resistance evolution depends on time after exposure

80 unique compounds were putatively matched to the *m/z* peaks forming the basis of the extended metabolic fingerprint of glyphosate resistance evolution. Out of 200 unique *m/z* peaks of interest (i.e. those with the largest fold differences between treatments), 151 could not be matched to a putative identity (Table 3). These masses likely reflect fragments of larger compounds or adducts not included here. The putative identities of the remaining 49 are listed in Table 9, with several being matched to more than one identity.

Most of the *m/z* peaks of interest were found to have either a pre-resistance or post-resistance pattern, with considerably fewer showing the persistent pattern (Table 3, Table 10, Table 11, Table 12). The putatively identified compounds in both the pre-resistance and post-resistance phase are diverse, including lipids, fatty acids, pigments, hormones, and redox electron carriers. The main pathway identified by MetaboAnalyst in both cases was carotenoid biosynthesis, albeit with different compounds highlighted for the different phases Figure 4. The pre-resistance phase was also linked to linoleic acid metabolism and ubiquinone and other terpenoid-quinoine biosynthesis pathways, while the post-resistance phase was linked to sesquiterpenoid and triterpenoid biosynthesis.

**Table 2:** Summary of significant pairwise differences between glyphosate-treated populations and controls for *m/z* peaks matched to AAs as well as the total AA pool. See **??** for the associated test statistics. Arrows and fold changes (calculated as the average glyphosate-treated ion count divided by the average control ion count) indicate the average % total ion count for the glyphosate-treated populations as compared to the controls. Greyed out cells represent no difference found between treatments. Cells are blacked out where no *m/z* peak could be matched to the target compound.

| AA | Day | Negative mode |  |  |  |  |  |  | Positive mode |  |  |  |  |  |  |
| --- | --- | --- | --- | --- | --- | --- | --- | --- | --- | --- | --- | --- | --- | --- | --- |
|  |  | -3 | 1 | 8 | 16 | 22 | 29 | 36 | -3 | 1 | 8 | 16 | 22 | 29 | 36 |
| Ala* |  |  |  |  |  |  |  |  | ↓0.88 | 0.99 | ↑1.18 | ↑1.15 | 0.98 | 0.98 | 0.94 |
| Arg |  | ↓0.9 | 0.91 | 0.93 | ↓0.73 | ↓0.73 | 0.96 | 0.95 | ↓0.88 | 1.01 | 1.04 | ↓0.82 | ↓0.75 | 0.97 | 1.03 |
| Asn |  | 1.03 | 0.96 | 0.95 | ↓0.77 | ↓0.81 | ↓0.9 | ↓0.88 |  |  |  |  |  |  |  |
| Asp |  | ↓0.86 | 1.01 | ↑1.59 | ↑1.48 | ↑1.26 | ↑1.24 | 1 | ↓0.9 | 1.02 | ↑1.1 | 0.97 | ↓0.9 | 0.94 | 0.99 |
| Cys |  | 1.12 | 1.07 | ↑1.48 | ↑3.04 | ↑2.5 | 1.36 | 1.2 | 0.98 | 0.92 | 1.14 | ↑1.65 | ↑1.4 | 1.06 | 1.06 |
| Glu |  | 1 | 0.99 | ↑1.36 | 0.99 | ↓0.9 | 1.05 | 1.04 | 1 | 0.95 | ↑1.31 | ↑1.15 | 0.93 | 1.04 | 1.05 |
| Gln |  | 0.94 | 1 | ↓0.71 | ↓0.51 | ↓0.56 | ↓0.85 | 0.9 | 1.01 | ↓0.89 | ↓0.83 | 0.99 | ↓0.92 | ↓0.92 | 0.99 |
| Gly |  |  |  |  |  |  |  |  |  |  |  |  |  |  |  |
| His |  | ↓0.9 | 0.98 | ↑1.11 | 0.99 | ↓0.91 | 0.99 | ↓0.87 |  |  |  |  |  |  |  |
| Ile/Leu |  | 0.89 | 1 | ↑1.16 | 1.01 | 0.87 | 0.94 | 0.85 | ↓0.91 | 0.97 | ↑1.15 | ↑1.19 | 1.01 | 0.99 | 0.97 |
| Lys |  | 0.95 | 0.93 | ↑1.19 | 0.97 | 0.95 | 1.03 | 0.94 | 1 | ↓0.93 | ↑1.12 | ↑1.24 | ↑1.1 | ↑1.07 | ↑1.06 |
| Met |  | 0.98 | ↑1.17 | ↑1.21 | 1.08 | 1 | ↑1.15 | ↓0.9 | 1.01 | ↓0.91 | ↓0.87 | ↓0.89 | ↓0.92 | ↓0.93 | ↓0.95 |
| Phe |  | 0.97 | 1.07 | 1.13 | 1.12 | 1.13 | 1.08 | ↓0.86 | ↓0.95 | ↓0.95 | 1 | ↑1.18 | ↑1.13 | 1.01 | 0.99 |
| Pro |  |  |  |  |  |  |  |  | ↓0.87 | 0.96 | 0.97 | ↓0.91 | ↓0.81 | ↓0.94 | 0.99 |
| Ser |  | 0.97 | 1.01 | ↑1.19 | ↓0.9 | ↓0.82 | 0.97 | ↓0.9 |  |  |  |  |  |  |  |
| Thr |  | 0.96 | 1.04 | ↑1.16 | 0.94 | ↓0.83 | 0.93 | ↓0.85 |  |  |  |  |  |  |  |
| Trp |  | ↓0.93 | ↓0.88 | 0.96 | ↓0.87 | ↓0.84 | ↓0.93 | ↓0.89 | 0.96 | 0.97 | ↓0.93 | 0.95 | 1 | 0.96 | 0.99 |
| Tyr |  | 0.95 | 1.08 | 1.09 | 1.04 | 1.01 | 1.06 | 0.9 | ↓0.96 | ↓0.96 | ↓0.96 | ↑1.04 | 1.01 | 1.01 | 0.98 |
| Val |  | 0.95 | ↓0.88 | 0.96 | ↓0.88 | ↓0.85 | ↓0.93 | ↓0.9 |  |  |  |  |  |  |  |
| AApool |  | 0.95 | 0.99 | ↑1.11 | 0.95 | ↓0.89 | 0.99 | 0.93 | ↓0.93 | ↓0.96 | 1.02 | 1.01 | ↓0.92 | 0.98 | 1.01 |

**Table 3:** Distribution of patterns classifications for unidentified and putatively identified peaks of interest.

| Pattern | Without putative ID match | With putative ID match | Total peaks of interest |
| --- | --- | --- | --- |
| Pre-resistance | 58 | 27 | 85 |
| Post-resistance | 65 | 18 | 83 |
| Persistent | 23 | 5 | 28 |
| Unclassifiable | 3 | 1 | 4 |
| Total | 149 | 51 | 200 |

Furthermore, the presence of a persistent pattern also suggests there is a distinctive metabolite profile for *C. reinhardtii* cells experiencing glyphosate action, regardless of whether the population is resistant or susceptible. The few putatively identified compounds here include fatty acids, carboxylates involved in antioxidant biosynthesis, lipids and redox electron carriers. However, the number of putatively identified compounds here was not enough to conclusively link to any specific metabolic pathways.

Four of the peaks of interest also showed a fluctuating pattern that did not fit the pre-defined patterns Table 13, with main differences between treatments found in the pre-resistance phase as well as on the final day, making their role harder to infer. The putatively matched identities for these compounds included fatty acids linked to arachidonic acid metabolism.

**Figure 4:**
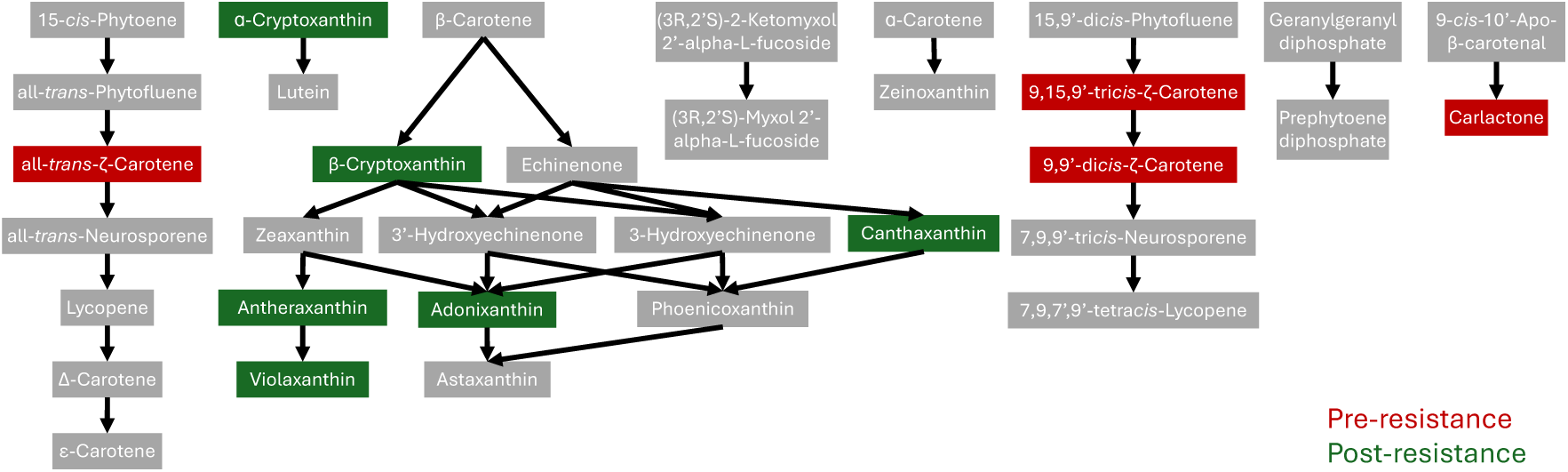
Peaks of interest in the pre-resistance phase are shown in red, peaks of interest in the post-resistance phase are shown in green. Linked compounds shown in grey.

## Discussion

Understanding how the evolutionary dynamics of herbicide resistance connect to the underlying molecular mechanisms is a vital step in developing precise weed management strategies that evade resistance evolution while limiting both overall herbicide use as well as damage to non-target organisms and ecosystems. Here we for the first time characterise the effect of glyphosate resistance evolution in action on the metabolome of model species *C. reinhardtii*, from before treatment introduction to after resistance has evolved, and use this metabolomic fingerprint to make inferences about the underlying resistance mechanism as well as associated fitness costs. We find a clear signal of rapidly evolving resistance alongside evidence of persistent fitness cost associated with widespread disruption of cellular processes, including ROS and lipid peroxidation. Furthermore, we could rule out glyphosate degradation and infer that the resistance mechanism was highly likely a target site mechanism and part of the standing genetic variation in the population.

### Evidence for glyphosate resistance evolution

Like previous work with other organisms, we detected a strong effect of glyphosate action on the metabolites upstream of EPSPS in the shikimate pathway[63, 64, 65, 66, 67, 68, 69, 25, 70, 71, 72], that then returns to a level comparable to the controls. This is not only strong evidence for glyphosate action, it is evidence for evolved resistance returning near normal cell function across the population with a possible trade off preventing a full return, and gives a timeline for resistance evolving in this system.

Similarly, evidence of general disruption is seen through ion count decreases in peaks matched to a number of AAs as well as the overall AA pool coinciding with the blockage of the shikimate pathway. This too returns to near control level by the end of the experiment as resistance has evolved, further suggesting that the resistance mechanism is highly effective at lifting the negative, inhibitory effects of glyphosate but may also trade off against normal cell function. The general effects of glyphosate inhibition on the AA pool are poorly understood, and appears to vary by system[70, 86, 65, 64, 63, 85, 87], timing[85, 87] and stress level[65]. While our results are consistent with previous studies showing the patterns of disruption differ depending on the AA, an the majority of AA *m/z* peaks show an effect of glyphosate treatment at some point, there is no evidence of triggering of parallel synthesis pathways or shifts toward more nitrogen rich AAs to buffer against the effects of glyphosate inhibition[86, 70, 65], especially as some AAs we would then expect an increase in (e.g. Gln or Val) show the opposite pattern. However, as the present study tests whole populations comprising several phenotypes with different resistance strategies emerging at different times, any such effects that are not largely unidirectional may be obscured.

It is likely the decreases in Trp and Thr are a direct effect of the blocked shikimate pathway, as they are downstream products, however Thr does also show a temporary increase on day 8. Phe, however, is found at lower levels than in the controls at the start of the experiment, including before the treatment start, and then increases. This might reflect cellular processes unrelated to glyphosate action, or Phe synthesis is prioritised over Trp and Tyr to maintain downstream biosyntheses[86]. This is also the possible explanation for Phe returning to control levels by the end of the experiment, while Trp and Thr remain reduced. Several studies have also noted decreased Glu coupled to increased Gln, suggesting Gln acting as a nitrogen donor for AA synthesis underlying this shift[70, 65, 86]. However, we found the opposite pattern, suggesting a different mechanism, or a masking effect of Glu by e.g. 2-oxoglutarate.

The *m/z* peaks identified for the extended metabolic fingerprint overwhelmingly showed a pattern where large differences between populations attributable to glyphosate action are lifted as resistance evolves, or emerge as resistance is evolving, either immediately or gradually. The gradual pattern likely reflects that the populations are heterogeneous[101, 102, 103, 104, 105, 106, 107, 108], containing the metabolomic signal of both resistant and dying phenotypes as dominance shifts. Alternatively, it could reflect soft sweeps to fixation as a particular resistant phenotype proportion is slowly growing, or mutation accumulation over time unrelated to the glyphosate treatment.

### Evaluation of possible resistance mechanisms

As the shikimate pathway, AA pool, and several other metabolites only return to near control population levels after resistance has evolved, there may be a trade-off of resistance against normal cell functioning. This suggests that if there is a target mutation to EPSPS, its sensitivity to the intended substrates may be reduced[31, 32, 33, 34, 35, 36, 37]. Similarly, if there is amplification of EPSPS, this is not sufficiently large to fully release the pathway blockage[44, 45, 46].

We would not necessarily expect all populations to exhibit the same dominant resistance mechanism, and there may be more than one mechanism present within each population as well as within each cell, and it is possible some non-resistant cells remain in the population by the end. However, as the level of inter-population variation found here is very low, and there is limited chance for *de novo*-mutation generation in a growth inhibited population, it is likely that the initial steps in resistance evolution are resulting from selection on standing genetic variation and that this is similar in all populations. Furthermore, there may be a limited number of mutations conferring glyphosate resistance that quickly become dominant, even though later mutations may lead to greater divergence between populations as the selective pressures change.

### No evidence for glyphosate degradation or changes in internal glyphosate concentration

There was no indication of glyphosate degradation conferring resistance, with the *m/z* peaks matched to possible catabolites showing patterns in the wrong direction. This is not unexpected as evidence for glyphosate catabolism being the primary resistance mechanism is highly limited[61, 109, 62, 110, 59, 60]. However, presence in a metabolite screen is stronger evidence than absence. A compound may still be present even if not detected in the expected *m/z* peak as it may have formed an unexpected adduct or fractured into smaller molecules[8]. Furthermore, as the metabolite screen is designed to give as broad coverage as possible, the settings used are not optimal for picking up all compounds, and ion suppression — where the finite ionisation energy is utilised to a higher degree by compounds that ionise easily, leaving little energy for other compounds — may reduce the signal of compounds in mixtures compared to separated injections[8, 7]. Targeted confirmatory analysis using e.g. LC-MS to isolate the compounds is thus needed to definitively rule out the presence of glyphosate catabolites.

There was also no evidence for changes in glyphosate levels. While the lack of detectable *m/z* peak may be accounted for by ion suppression, it is more likely due to it primarily being present bound by enzymes. If EPSPS is up-regulated, we would not expect any free glyphosate as it would all be bound by EPSPS[1]. Similarly for increased vacuolar sequestration, glyphosate may be bound by transport enzymes[54, 55, 56]. While mutated EPSPS with lower affinity for glyphosate could theoretically result in free glyphosate, the absence of an increase does not exclude this mechanism as it may work in tandem with other mechanisms. Furthermore, with no detectable intra-cellular glyphosate before resistance evolves, we cannot determine whether resistance involves changed absorption into the cell.

### Evidence for persisting oxidative damage and membrane changes

While evolved resistance lifts the majority glyphosate inhibition effects, a number of *m/z* peaks still show marked differences between treatments by the end of the experiment. Some have persisted from the treatment start, suggesting they are direct effects of glyphosate, whereas some emerge gradually or around the same time as resistance, suggesting they may be related to resistance costs. The putatively identified compounds found here mainly include lipids, fatty acids, electron-transfer quinols and carotenoids, suggesting the differences may involve cell membranes and oxidative damage.

While the carbon flow disruption may be enough to upset the redox balance of the cell[25], and ROS production is commonly found as a secondary effect of different stressors[111, 112, 113, 114, 115, 116], Gomes (2016)[26] found that glyphosate interferes with the mitochondrial electron transport chain to produce ROS. This mechanism may explain the reported glyphosate effects on non-target organisms lacking a shikimate pathway (e.g. animals)[117, 118], and suggests glyphosate could cause continued damage after inhibition of the shikimate pathway is lifted, including lipid peroxidation, the chain reaction of free radical propagation in which electrons are stolen from membrane lipids[115]. Increased antioxidant activity has been found in glyphosate resistant strains in several systems[25, 119, 120], and in some glyphosate continues to have an effect on seed germination through oxidative damage[121].

The electron-transfer quinols/quinones putatively identified are menaquinols and ubiquinols/quinones as well as plastoquinol, compounds involved in electron transport and signalling across membranes, but also antioxidants that reduce damage from ROS and prevent lipid peroxidation[122, 123, 124, 125, 126]. A number of the putatively identified compounds across all three patterns are also involved in the synthesis of electron-transfer quinols, such as demethylphylloquinone[127].

Biosynthesis of secondary carotenoids like the putatively identified adonixanthin is a known general stress response in algae, as well as a ROS production response as they are singlet oxygen scavengers[128, 129]. Immediate abundance changes of these compounds may reflect a plastic response to oxidative damage, whereas differences emerging post-resistance may reflect adaptation. While our data cannot reveal how effective this response is and if the level of ROS present in the cell or the associated damage changes throughout the adaptation process, their continued presence suggests an ongoing stress response separate from shikimate pathway inhibition. The putatively identified lipids are primarily involved in glycerophospholipid metabolism, producing the lipids involved in cellular membranes[130]. Changes here could either be a sign of membrane degradation from lipid peroxidation, or may reflect membrane structure changes to limit either primary or secondary glyphosate damage[131, 111].

Free fatty acids is another possible sign of membrane degradation by lipid peroxidation[111, 113, 116], but the *m/z* peaks putatively matched to fatty acids here, e.g. pristanate and linoleate, show significant differences pre-resistance. This suggests this signal reflects the death and breakdown of glyphosate susceptible cells, and the stabilisation to control levels corresponds to the shift to a population dominated by glyphosate resistant cells not experiencing the same levels of membrane damage regardless of ROS levels. Lastly, some AAs, like Cys, Met and Pro, may also serve a role in cellular redox-regulation[132, 116]. As some of these compounds are only elevated during the pre-resistance phase, they may reflect a first line of defence[112, 133], with stabilisation reflecting a shift to other mechanisms for redox control.

Targeted analysis of implicated pathways, as well as analysis of ROS and lipid peroxidation levels through targeting likely products[116], would provide insight into the severity of the damage and thus selective pressure extent. Furthermore, metabolomic analysis of resistant cells experiencing the ancestral environment would allow untangling adaptive and plastic responses, as well as effects of glyphosate action and the resistance trait.

### Limitations, broader implications and future research

Untargeted metabolomic screening is a powerful hypothesis generating tool, extracting as much information as possible out of a small sample with a high throughput and unbiased broad spectrum analysis of altered biochemical pathways[8]. However, the method comes with a number of caveats, including ion suppression and inability to conclusively identify compounds or definitely rule out their presence. There is also no straightforward connection between the relative ion count and the underlying reason. Abundance may increase due to up-regulation or build-up as downstream pathways are blocked or diverted, and it may decrease by down-regulation, increased demand or catabolism.

While MS/MS analysis strengthened the evidence for the presence of a number of targeted and putatively identified compounds, we were limited by the availability of MassBank reference spectra and could not test all compounds of interest. Targeted analysis using chromatography would allow both confirming the absence of specific compounds (e.g. glyphosate degradation products) and with higher accuracy quantify others as a measure of pathway functioning (e.g. shikimate-3-phosphate). It is also worth noting that the extended metabolomic fingerprint analysis presented here is only the tip of the iceberg, as each analysed sample generates over 10 000 *m/z* peaks. More extensive data analysis would reveal more compounds of interest to generate more specific hypotheses for further targeted analysis. In particular, targeted analysis of metabolites involved in membrane lipid production as well as cellular redox balance would provide more material for evaluating the persistent effects of glyphosate action as these have the potential to affect resistant crops as well as weeds and non-target species after resistance has evolved. A longer running study combined with exploratory analysis targeting *m/z* peaks with high variance in treated populations but not controls would reveal if there are later divergences in strategy as the populations continue to adapt to glyphosate, and could also reveal how persistent the oxidative damage is, and whether more extensive defences against it evolves. Connecting the phenotype observed to a genotype through employing genomics and transcriptomics would also allow greater understanding of the genetic underpinnings, specifically to test for substitutions in the EPSPS gene and whether its transcription has been up-regulated[44, 45, 46, 34, 35, 36, 37].

Metabolomic fingerprinting is also a useful tool for monitoring environmental pollutants in natural ecosystems[134], and our results emphasise that adaptation to pollutants needs to be taken into account and that adaptation stage matters for the associated fingerprint. While the acute metabolomic response to glyphosate is build-up of shikimate pathway products, this is only weakly present in the profile of the resistant population, a shift that happens relatively quickly. However, our results suggest there is indeed a distinctive metabolite profile for *C. reinhardtii* cells experiencing glyphosate action regardless of whether they are resistant or susceptible, but this is primarily related to oxidative damage and membrane lipids rather than the shikimate pathway.

In addition, this has the potential for far reaching consequences in glyphosate contaminated ecosystems, especially if these are permanent changes to the algal cells. Firstly, as primary producers, their biochemical composition determines their quality as food and thus the energy transferred to other trophic levels[111]. With sufficiently large changes, foraging strategies of grazing species may be affected or ultimately, food webs restructured. Secondly, if changed membrane lipid content indeed corresponds to changed membrane structure, this may affect the clumping anti-grazer response or the digestability of the cells[135, 136, 137, 138, 139, 140, 141, 142, 143], which affects energy transfer to grazing species. Thirdly, as oxidative damage is a secondary effect of many stressors[116], this provides a mechanism for interactive effects of stressors. Adaptation to the oxidative damage of glyphosate may provide a generalist defence against damage from other stressors, but a combination of stressors could also have an additive or synergistic effect[144, 145].

Lastly, while the exact molecular mechanisms may be species dependent, our study highlights the necessity of tackling the problem of herbicide resistance and management as a continuous, population level evolutionary process. Particularly pertinent is understanding the full effects of the herbicide on a molecular level at all stages of resistance evolution with temporally-resolved insight to be able to predict its population- and ecosystem-level effects, as secondary effects may continue to affect the organism after resistance to the primary mode of action has evolved.

