## Supplementary materials for "Metabolomic profiling of glyphosate resistance evolution in green alga *Chlamydomonas reinhardtii*"

### Supporting materials

Table 1: Conditions for sample introduction into mass spectrometer.

| Condition | Negative mode | Positive mode |
|---|---|---|
| Capillary voltage | 2.5 kV | 2.8 kV |
| Sample cone | 20 V | 80 V |
| Source offset | 80 V | 20 V |
| Desolvation gas flow | 300 L/h | 500 L/h |
| Desolvation temperature | 280°C | 280°C |
| Source temperature | 100°C | 100°C |

Table 2: Compounds targeted in MS/MS analysis, divided by hypothesis. Peaks from reference spectra have been excluded if below the detection limit of the instrument (<50 Da) or had a relative expected intensity below 30.

| Compound | Accurate mass | Target mass (mode) | Collision energy | MassBank reference spectrum | Major expected peaks | Present? |
|---|---|---|---|---|---|---|
| *Shikimate pathway* | | | | | | |
| D-Erythrose-4P (E4P) | 200.0086 | 199(-) | 10 | PR100545 | 96.9703 ✓<br>78.9602 ✓<br>138.9800 ✓<br>199.0008 ✓<br>154.9088 ✓ | ✓ |
| Shikimate | 174.0528 | 173(-) | 30 | KO001789 | 119.0000 ✓<br>173.3000 ✓<br>137.1000 ✓<br>141.0000 ✓<br>59.2000 ✗<br>172.9000 ✓<br>155.3000 ✓<br>93.3000 ✗<br>111.2000 ✓<br>113.0000 ✓<br>77.0000 ✗ | ✓ |
| *Amino acids* | | | | | | |
| Alanine (Ala)* | 89.0477 | 90(+) | 5 | KO002053 | 90.0000 ✓ | ✓ |
| Arginine (Arg) | 174.1117 | 175(+) | 10 | CE000265 | 158.0923 ✓<br>157.1083 ✓<br>116.0705 ✓<br>130.0975 ✓<br>175.1190 ✓ | ✓ |
| Arginine (Arg) | 174.1117 | 173.2(-) | 20 | PR100578 | 131.0817 ✓<br>173.1039 ✓ | ✓ |
| Asparagine (Asn) | 132.0535 | 131(-) | 20 | KO000025 | 114.0000 ✓<br>113.3000 ✗<br>70.0000 ✗<br>95.3000 ✗<br>131.0000 ✓<br>70.8000 ✗<br>71.9000 ✗<br>58.2000 ✗ | ✓ |
| Aspartate (Asp) | 133.0375 | 132(-) | 10 | CE000453 | 132.0309 ✓<br>88.0410 ✓<br>115.0043 ✓<br>114.0203 ✓ | ✓ |
| Aspartate (Asp) | 133.0375 | 134(+) | 25 | KO002065 | 74.0000 ✗<br>70.2000 ✗<br>88.2000 ✗<br>71.0000 ✗<br>72.1000 ✓ | ✗ |
| Cysteine (Cys) | 121.0197 | 122(+) | 30 | KO000848 | 58.9960 ✗<br>76.0230 ✓<br>86.9900 ✓ | ✗ |
| Glutamate (Glu) | 147.0532 | 146(-) | 10 | KO000848 | 146.1000 ✓<br>127.9000 ✓<br>102.0000 ✓ | ✓ |
| Glutamine (Gln) | 146.0691 | 145(-) | 10 | KO000843 | 145.0000 ✓<br>144.6000 ✓<br>127.0000 ✓ | ✓ |

Continued on next page

Table 2 – continued from previous page

| **Compound** | **Accurate mass** | **Target mass (mode)** | **Collision energy** | **MassBank reference spectrum** | **Major expected peaks** | **Present?** |
|---|---|---|---|---|---|---|
| Histidine (His) | 155.0695 | 154(-) | 20 | BML01100 | 137.0347 ✓<br>110.0720 ✓<br>154.0613 ✓<br>136.0498 ✓<br>118.0387 ✓<br>109.0375 ✓<br>108.0607 ✓<br>137.0719 ✓<br>106.0393 ✓<br>154.0373 ✓ | ✓ |
| Isoleucine (Ile) | 131.0946 | 130(-) | 30 | KO001177 | 130.0000 ✓<br>82.0000 ✗<br>58.7000 ✗ | |
| Isoleucine (Ile) | 131.0946 | 132(+) | 10 | KO003173 | 86.2000 ✓<br>69.2000 ✓<br>115.3000 ✓<br>98.0000 ✓ | ✓ |
| Leucine (Leu) | 131.0946 | 130(-) | 30 | KO001263 | 130.0000 ✓<br>81.9000 ✗<br>84.2000 ✗<br>66.1000 ✗<br>71.9000 ✗ | |
| Leucine (Leu) | 131.0946 | 132(+) | 10 | KO003278 | 86.2000 ✓<br>115.1000 ✓ | ✓ |
| Lysine (Lys) | 146.1055 | 147(-) | 30 | RP001112 | 97.0781 ✓ | ✓ |
| Methionine (Met) | 149.051 | 150(+) | 20 | KO003333 | 104.1000 ✓<br>56.3000 ✗<br>133.1000 ✓<br>101.8000 ✓<br>61.1000 ✗<br>73.0000 ✓<br>74.0000 ✗<br>87.3000 ✗<br>150.3000 ✗<br>114.4000 ✗<br>86.2000 ✓<br>84.2000 ✓<br>85.0000 ✓<br>115.4000 ✗<br>101.0000 ✓<br>105.0000 ✓ | ✓ |
| Phenylalanine (Phe) | 165.079 | 166(+) | 20 | RP000402 | 120.0807 ✓<br>103.0541 ✓<br>93.0694 ✓<br>121.0840 ✓<br>93.0694 ✓<br>79.0538 ✓ | ✓ |
| Phenylalanine (Phe) | 165.079 | 164(-) | 20 | RP000412 | 103.0558 ✓<br>147.0447 ✓<br>164.0727 ✓<br>104.0585 ✓<br>148.0521 ✓ | ✓ |
| Proline (Pro) | 115.0633 | 116(+) | 20 | KO003673 | 70.0000 ✓ | ✓ |
| Serine (Ser) | 105.0426 | 104(-) | 20 | KO001781 | 74.2000 ✗<br>103.9000 ✓ | ✗ |
| Threonine (Thr) | 119.0582 | 118(-) | 20 | KO001878 | 74.1000 ✗<br>118.1000 ✓ | ✗ |
| Tryptophan (Trp) | 204.0899 | 205(+) | 20 | KO004073 | 188.2000 ✓<br>146.1000 ✓<br>144.1000 ✓<br>159.1000 ✓<br>118.1000 ✓<br>170.1000 ✓ | ✓ |
| Tryptophan (Trp) | 204.0899 | 203(-) | 20 | KO001868 | 203.3000 ✓<br>116.0000 ✓<br>73.9000 ✓<br>143.4000 ✗<br>141.9000 ✓<br>159.2000 ✓<br>186.2000 ✓<br>130.0000 ✓<br>129.0000 ✓ | ✓ |
| Tyrosine (Tyr) | 181.0739 | 180(-) | 20 | KO001863 | 163.1000 ✓<br>180.4000 ✗<br>119.3000 ✗<br>92.8000 ✓<br>136.1000 ✓<br>72.2000 ✗ | ✓ |

Continued on next page

Table 2 – continued from previous page

| **Compound** | **Accurate mass** | **Target mass (mode)** | **Collision energy** | **MassBank reference spectrum** | **Major expected peaks** | **Present?** |
|---|---|---|---|---|---|---|
| | | | | | 147.8000 ✗<br>74.2000 ✗<br>106.0000 ✓<br>107.0000 ✓ | |
| Tyrosine (Tyr) | 181.0739 | 182(+) | 10 | KO004067 | 182.1000 ✓<br>165.2000 ✓<br>136.0000 ✓<br>122.2000 ✓<br>121.0000 ✓ | ✓ |
| Valine (Val) | 117.079 | 116(-) | 30 | KO001990 | 116.2000 ✓ | ✓ |
| *Glyphosate catabolites* | | | | | | |
| Sarcosine* | 89.0477 | 88(-) | 10 | KO001799 | 88.2000 ✓ | ✓ |
| Sarcosine* | 89.0477 | 90(+) | 5 | KO003987 | 90.0000 ✓ | ✓ |
| phosphate | 97.9769 | 97(-) | 40 | RP022413 | 78.9591 ✓<br>62.9641 ✗ | ✓ |
| 2-oxoglutarate | 146.0215 | 145(-) | 10 | KO001528 | 145.1000 ✓<br>101.1000 ✓ | ✓ |
| Cinnamyl alcohol | 134.0732 | 135(+) | 10 | PS044801 | 135.0000 ✓<br>134.0000 ✓<br>73.0000 ✓<br>93.0000 ✓ | ✓ |
| *Exploratory analysis compounds* | | | | | | |
| linoleate | 280.2402 | 303(+) | 20 | PS027803 | 281.0000 ✓<br>73.0000 ✓<br>265.0000 ✓<br>264.0000 ✓<br>248.0000 ✓ | |
| thyroxine | 776.6867 | 777(+) | 15 | AU274306 | 731.6904 ✗<br>604.7811 ✗<br>350.9754 ✗<br>379.9523 ✗<br>323.9650 ✗<br>633.7740 ✓<br>732.6928 ✓<br>576.7651 ✓ | ✗ |
| | | 759(+) | | | 731.6904 ✗<br>604.7811 ✗<br>350.9754 ✗<br>379.9523 ✓<br>323.9650 ✗<br>633.7740 ✗<br>732.6928 ✓<br>576.7651 ✗ | ✗ |

Table 3: Analysis of Deviance Table (Type II Wald $\chi^2$-tests) for linear mixed effects model to test effect of glyphosate resistance evolution on shikimate pathway compounds with percentage ion count (arc sine transformed) of the putatively matched target compound as response. Where the lmm yielded a singular fit, an lm has been fit instead. Pairwise contrasts by day, as estimated by the `emmeans` package for R.

| *Shikimate pathway functioning* | | | | | | | | | |
|---|---|---|---|---|---|---|---|---|---|
| | **lmm/lm** | | | | **Pairwise contrasts** | | | | |
| **Compound** | *fixed effect* | $\chi^2/F$ | *df* | *p* | *contrast* | *t-ratio* | *df* | *p* | *fold change* |
| Erythrose-4P (-) (lmm) | day | 126 | 6 | <0.0001 | -3 | 5.3 | 31 | <0.0001 | 0.77 |
| | treatment | 1.4 | 1 | 0.24 | 1 | -2.6 | 31 | 0.014 | 1.13 |
| | day:treatment | 270.5 | 6 | <0.0001 | 8 | -5.1 | 31 | <0.0001 | 1.28 |
| | | | | | 16 | 8.6 | 31 | <0.0001 | 0.59 |
| | | | | | 22 | 4.9 | 31 | <0.0001 | 0.74 |
| | | | | | 29 | -3.5 | 31 | 0.002 | 1.2 |
| | | | | | 36 | -2.2 | 31 | 0.034 | 1.12 |
| DAHP (-) (lmm) | day | 11 | 6 | 0.09 | -3 | -0.1 | 54 | 0.9 | 1.01 |
| | treatment | 40.7 | 1 | <0.0001 | 1 | 3.9 | 54 | <0.0001 | 0.84 |
| | day:treatment | 15.9 | 6 | 0.014 | 8 | 4 | 54 | <0.0001 | 0.83 |
| | | | | | 16 | 4 | 54 | <0.0001 | 0.83 |
| | | | | | 22 | 4.1 | 54 | <0.0001 | 0.83 |
| | | | | | 29 | 2.4 | 54 | 0.021 | 0.9 |
| | | | | | 36 | 2.4 | 54 | 0.02 | 0.9 |
| 3-dehydro-shikimate (-) (lm) | day | 39.4 | 6,56 | <0.0001 | -3 | 0.7 | 56 | 0.48 | 0.96 |
| | treatment | 258 | 1,56 | <0.0001 | 1 | -2 | 56 | 0.054 | 1.12 |
| | day:treatment | 54.3 | 6,56 | <0.0001 | 8 | -21.3 | 56 | <0.0001 | 2.41 |
| | | | | | 16 | -7.5 | 56 | <0.0001 | 1.47 |
| | | | | | 22 | -0.9 | 56 | 0.39 | 1.05 |
| | | | | | 29 | -4.8 | 56 | <0.0001 | 1.3 |
| | | | | | 36 | -6.9 | 56 | <0.0001 | 1.45 |
| Shikimate (-) (lm) | day | 67.5 | 6,56 | <0.0001 | -3 | -0.2 | 56 | 0.87 | 1.01 |
| | treatment | 984.7 | 1,56 | <0.0001 | 1 | -3.1 | 56 | 0.003 | 1.26 |
| | day:treatment | 85.9 | 6,56 | <0.0001 | 8 | -28.8 | 56 | <0.0001 | 3.8 |
| | | | | | 16 | -13.4 | 56 | <0.0001 | 2.25 |
| | | | | | 22 | -9.6 | 56 | <0.0001 | 1.9 |
| | | | | | 29 | -14.3 | 56 | <0.0001 | 2.47 |
| | | | | | 36 | -13.7 | 56 | <0.0001 | 2.26 |
| Shikimate-3-phosphate (-) (lmm) | day | 428 | 6 | <0.0001 | -3 | 0.1 | 52 | 0.9 | 0.98 |
| | treatment | 355.9 | 1 | <0.0001 | 1 | -1.1 | 52 | 0.29 | 1.2 |
| | day:treatment | 436.7 | 6 | <0.0001 | 8 | -17.7 | 52 | <0.0001 | 4.25 |
| | | | | | 16 | -20.4 | 52 | <0.0001 | 4.68 |
| | | | | | 22 | -13.1 | 52 | <0.0001 | 3.39 |
| | | | | | 29 | -8.4 | 52 | <0.0001 | 2.55 |
| | | | | | 36 | -4.9 | 52 | <0.0001 | 1.83 |
| EPSP (-) (lm) | day | 4 | 6,56 | 0.002 | -3 | 1.1 | 56 | 0.29 | 0.96 |
| | treatment | 21.5 | 1,56 | <0.0001 | 1 | 2.2 | 56 | 0.033 | 0.92 |
| | day:treatment | 0.7 | 6,56 | 0.64 | 8 | 1.8 | 56 | 0.081 | 0.94 |
| | | | | | 16 | 0.7 | 56 | 0.48 | 0.98 |
| | | | | | 22 | 1.9 | 56 | 0.067 | 0.93 |
| | | | | | 29 | 3.3 | 56 | 0.002 | 0.89 |
| | | | | | 36 | 1.4 | 56 | 0.18 | 0.95 |
| Chorismate /Prephenate (-) (lmm) | day | 13.9 | 6 | 0.031 | -3 | 1.1 | 52 | 0.26 | 0.91 |
| | treatment | 7.9 | 1 | 0.005 | 1 | -3.4 | 52 | 0.001 | 1.27 |
| | day:treatment | 18 | 6 | 0.006 | 8 | -0.5 | 52 | 0.63 | 1.04 |
| | | | | | 16 | -2.1 | 52 | 0.041 | 1.17 |
| | | | | | 22 | -0.9 | 52 | 0.37 | 1.07 |
| | | | | | 29 | -0.8 | 52 | 0.44 | 1.06 |
| | | | | | 36 | -3.2 | 52 | 0.002 | 1.24 |

Table 4: Analysis of Deviance Table (Type II Wald $\chi^2$-tests) for linear mixed effects model to test presence of secondary glyphosate and AMPA breakdown products with percentage ion count (arc sine transformed) of the putatively matched target compound as response. Where the lmm yielded a singular fit, an lm has been fit instead. Pairwise contrasts by day, as estimated by the `emmeans` package for R.

| *Glyphosate catabolites* | | | | | | | | | |
|---|---|---|---|---|---|---|---|---|---|
| | **lmm/lm** | | | | **Pairwise contrasts** | | | | |
| **Compound** | *fixed effect* | $\chi^2/F$ | *df* | *p* | *contrast* | *t-ratio* | *df* | *p* | *fold change* |
| 2-oxoglutarate (-) (lmm) | day | 41 | 6 | <0.0001 | -3 | 2.8 | 42 | 0.009 | 0.8 |
| | treatment | 21.6 | 1 | <0.0001 | 1 | 1 | 42 | 0.31 | 0.92 |
| | day:treatment | 13.9 | 6 | 0.031 | 8 | 1.3 | 42 | 0.2 | 0.87 |
| | | | | | 16 | 3.4 | 42 | 0.001 | 0.69 |
| | | | | | 22 | 5 | 42 | <0.0001 | 0.6 |

Continued on next page

Table 4 – continued from previous page

| *Glyphosate catabolites* | | | | | | | | | |
|---|---|---|---|---|---|---|---|---|---|
| | **lmm/lm** | | | | **Pairwise contrasts** | | | | |
| **Compound** | *fixed effect* | $\chi^2/F$ | *df* | *p* | *contrast* | *t-ratio* | *df* | *p* | *fold change* |
| | | | | | 29 | 2.5 | 42 | 0.018 | 0.77 |
| | | | | | 36 | 3.1 | 42 | 0.004 | 0.72 |

Continued on next page

Table 4 – continued from previous page

| *Glyphosate catabolites* | | | | | | | | | |
|---|---|---|---|---|---|---|---|---|---|
| | **lmm/lm** | | | | **Pairwise contrasts** | | | | |
| **Compound** | *fixed effect* | $\chi^2/F$ | *df* | *p* | *contrast* | *t-ratio* | *df* | *p* | *fold change* |
| 5-phospho-α-D-ribose 1,2-cyclic phosphate (-) (lm) | day | 1.9 | 6,56 | 0.096 | -3 | 0.4 | 56 | 0.72 | 0.97 |
| | treatment | 31.5 | 1,56 | <0.0001 | 1 | 0.4 | 56 | 0.71 | 0.97 |
| | day:treatment | 2.1 | 6,56 | 0.072 | 8 | 3.3 | 56 | 0.002 | 0.78 |
| | | | | | 16 | 3.8 | 56 | <0.0001 | 0.76 |
| | | | | | 22 | 2.4 | 56 | 0.02 | 0.84 |
| | | | | | 29 | 3.3 | 56 | 0.002 | 0.79 |
| | | | | | 36 | 1.3 | 56 | 0.19 | 0.91 |
| 5-phospho-α-D-ribose 1,2-cyclic phosphate (+) (lmm) | day | 17.4 | 6 | 0.008 | -3 | 1.7 | 44 | 0.091 | 0.9 |
| | treatment | 22.8 | 1 | <0.0001 | 1 | 1.3 | 44 | 0.2 | 0.92 |
| | day:treatment | 52.9 | 6 | <0.0001 | 8 | 0 | 44 | 0.99 | 1 |
| | | | | | 16 | 4.8 | 44 | <0.0001 | 0.73 |
| | | | | | 22 | 7.7 | 44 | <0.0001 | 0.6 |
| | | | | | 29 | 1.9 | 44 | 0.061 | 0.88 |
| | | | | | 36 | 1.6 | 44 | 0.13 | 0.91 |
| 5-phospho-α-D-ribose 1-diphosphate (-) (lmm) | day | 36.7 | 6 | <0.0001 | -3 | 0.6 | 37 | 0.53 | 0.95 |
| | treatment | 26.7 | 1 | <0.0001 | 1 | 0.5 | 37 | 0.64 | 0.97 |
| | day:treatment | 66.7 | 6 | <0.0001 | 8 | 1.5 | 37 | 0.15 | 0.91 |
| | | | | | 16 | 7.1 | 37 | <0.0001 | 0.59 |
| | | | | | 22 | 7 | 37 | <0.0001 | 0.6 |
| | | | | | 29 | 3.8 | 37 | 0.001 | 0.79 |
| | | | | | 36 | 2.4 | 37 | 0.023 | 0.86 |
| AMPA (-) (lmm) | day | 205.9 | 6 | <0.0001 | -3 | -1.7 | 51 | 0.089 | 1.08 |
| | treatment | 13.6 | 1 | <0.0001 | 1 | -5.8 | 51 | <0.0001 | 1.25 |
| | day:treatment | 30.6 | 6 | <0.0001 | 8 | -2.2 | 51 | 0.033 | 1.08 |
| | | | | | 16 | -0.4 | 51 | 0.68 | 1.01 |
| | | | | | 22 | 1 | 51 | 0.33 | 0.97 |
| | | | | | 29 | -1.2 | 51 | 0.23 | 1.04 |
| | | | | | 36 | -2.7 | 51 | 0.01 | 1.1 |
| Cinnamaldehyde (-) (lm) | day | 3 | 6,56 | 0.012 | -3 | -0.6 | 56 | 0.57 | 1.03 |
| | treatment | 40.3 | 1,56 | <0.0001 | 1 | 1 | 56 | 0.33 | 0.96 |
| | day:treatment | 4.4 | 6,56 | 0.001 | 8 | 1.3 | 56 | 0.2 | 0.95 |
| | | | | | 16 | 5.6 | 56 | <0.0001 | 0.77 |
| | | | | | 22 | 4.4 | 56 | <0.0001 | 0.81 |
| | | | | | 29 | 2.5 | 56 | 0.016 | 0.9 |
| | | | | | 36 | 2.6 | 56 | 0.012 | 0.88 |
| Cinnamaldehyde (+) (lm) | day | 5 | 6,56 | <0.0001 | -3 | 1.1 | 56 | 0.28 | 0.95 |
| | treatment | 6.3 | 1,56 | 0.015 | 1 | 1.4 | 56 | 0.16 | 0.93 |
| | day:treatment | 0.4 | 6,56 | 0.89 | 8 | 0.9 | 56 | 0.35 | 0.95 |
| | | | | | 16 | -0.2 | 56 | 0.88 | 1.01 |
| | | | | | 22 | 0.8 | 56 | 0.43 | 0.96 |
| | | | | | 29 | 1.8 | 56 | 0.072 | 0.9 |
| | | | | | 36 | 0.7 | 56 | 0.47 | 0.96 |
| Cinnamyl alcohol (-) (lm) | day | 4.8 | 6,56 | <0.0001 | -3 | -0.3 | 56 | 0.75 | 1.03 |
| | treatment | 16.6 | 1,56 | <0.0001 | 1 | 2.4 | 56 | 0.02 | 0.78 |
| | day:treatment | 1.7 | 6,56 | 0.13 | 8 | 1.6 | 56 | 0.11 | 0.84 |
| | | | | | 16 | 1.1 | 56 | 0.27 | 0.88 |
| | | | | | 22 | 1.6 | 56 | 0.12 | 0.83 |
| | | | | | 29 | 0.6 | 56 | 0.56 | 0.93 |
| | | | | | 36 | 3.8 | 56 | <0.0001 | 0.67 |
| Cinnamyl alcohol (+) (lm) | day | 9.5 | 6,56 | <0.0001 | -3 | 1.1 | 56 | 0.26 | 0.97 |
| | treatment | 3.7 | 1,56 | 0.061 | 1 | 4.6 | 56 | <0.0001 | 0.89 |
| | day:treatment | 6.4 | 6,56 | <0.0001 | 8 | 0.4 | 56 | 0.68 | 0.99 |
| | | | | | 16 | -3.8 | 56 | <0.0001 | 1.1 |
| | | | | | 22 | -0.4 | 56 | 0.7 | 1.01 |
| | | | | | 29 | 1.9 | 56 | 0.059 | 0.95 |
| | | | | | 36 | 1.1 | 56 | 0.26 | 0.97 |
| Phosphate (-) (lmm) | day | 108.3 | 6 | <0.0001 | -3 | 4.1 | 52 | <0.0001 | 0.84 |
| | treatment | 30.8 | 1 | <0.0001 | 1 | -5.1 | 52 | <0.0001 | 1.22 |
| | day:treatment | 105.6 | 6 | <0.0001 | 8 | -8.2 | 52 | <0.0001 | 1.31 |
| | | | | | 16 | -2.1 | 52 | 0.041 | 1.08 |
| | | | | | 22 | 0.2 | 52 | 0.83 | 0.99 |
| | | | | | 29 | -3.5 | 52 | 0.001 | 1.15 |
| | | | | | 36 | -4.4 | 52 | <0.0001 | 1.19 |
| Sarcosine* (+) (lm) | day | 12.9 | 6,56 | <0.0001 | -3 | 3.9 | 56 | <0.0001 | 0.88 |
| | treatment | 0.9 | 1,56 | 0.34 | 1 | 0.2 | 56 | 0.82 | 0.99 |
| | day:treatment | 10.8 | 6,56 | <0.0001 | 8 | -5.3 | 56 | <0.0001 | 1.18 |
| | | | | | 16 | -4.4 | 56 | <0.0001 | 1.15 |
| | | | | | 22 | 0.7 | 56 | 0.47 | 0.98 |
| | | | | | 29 | 0.5 | 56 | 0.61 | 0.98 |
| | | | | | 36 | 1.7 | 56 | 0.093 | 0.94 |
| α-D-ribose 1,5-bisphosphate (-) (lmm) | day | 61.5 | 6 | <0.0001 | -3 | 0 | 56 | 0.99 | 1 |
| | treatment | 106.1 | 1 | <0.0001 | 1 | 1.3 | 56 | 0.18 | 0.94 |
| | day:treatment | 64.3 | 6 | <0.0001 | 8 | 4.7 | 56 | <0.0001 | 0.8 |
| | | | | | 16 | 8.4 | 56 | <0.0001 | 0.66 |
| | | | | | 22 | 8 | 56 | <0.0001 | 0.67 |
| | | | | | 29 | 4.8 | 56 | <0.0001 | 0.81 |
| | | | | | 36 | 2.1 | 56 | 0.044 | 0.92 |

Continued on next page

Table 4 – continued from previous page

| *Glyphosate catabolites* | | | | | | | | | |
|---|---|---|---|---|---|---|---|---|---|
| | **lmm/lm** | | | | **Pairwise contrasts** | | | | |
| **Compound** | *fixed effect* | $\chi^2/F$ | *df* | *p* | *contrast* | *t-ratio* | *df* | *p* | *fold change* |
| α-D-ribose 1,5-bisphosphate (+) (lm) | day | 8.5 | 6,56 | <0.0001 | -3 | 0.9 | 56 | 0.37 | 0.94 |
| | treatment | 2.4 | 1,56 | 0.13 | 1 | 0.5 | 56 | 0.6 | 0.96 |
| | day:treatment | 10 | 6,56 | <0.0001 | 8 | -3.4 | 56 | 0.001 | 1.23 |
| | | | | | 16 | 2.6 | 56 | 0.013 | 0.82 |
| | | | | | 22 | 5.8 | 56 | <0.0001 | 0.66 |
| | | | | | 29 | 0.7 | 56 | 0.5 | 0.95 |
| | | | | | 36 | -3 | 56 | 0.004 | 1.19 |
| α-D-ribose 1-(acetamidomethyl-phosphonate) 5-triphosphate (-) (lmm) | day | 32.3 | 6 | <0.0001 | -3 | 0.5 | 43 | 0.64 | 0.96 |
| | treatment | 32.3 | 1 | <0.0001 | 1 | 3.7 | 43 | 0.001 | 0.73 |
| | day:treatment | 27 | 6 | <0.0001 | 8 | 1.7 | 43 | 0.095 | 0.87 |
| | | | | | 16 | 5.8 | 43 | <0.0001 | 0.56 |
| | | | | | 22 | 5.2 | 43 | <0.0001 | 0.61 |
| | | | | | 29 | 3.3 | 43 | 0.002 | 0.77 |
| | | | | | 36 | 2.9 | 43 | 0.006 | 0.8 |
| α-D-ribose-1-(2-N-acetamidomethyl-phosphonate) 5-phosphate (-) (lmm) | day | 98.1 | 6 | <0.0001 | -3 | 0.4 | 37 | 0.7 | 0.96 |
| | treatment | 8.1 | 1 | 0.004 | 1 | -0.4 | 37 | 0.71 | 1.03 |
| | day:treatment | 31.2 | 6 | <0.0001 | 8 | 0.5 | 37 | 0.64 | 0.97 |
| | | | | | 16 | 3.6 | 37 | 0.001 | 0.75 |
| | | | | | 22 | 4.9 | 37 | <0.0001 | 0.68 |
| | | | | | 29 | 2.4 | 37 | 0.021 | 0.85 |
| | | | | | 36 | 1.1 | 37 | 0.26 | 0.93 |
| α-D-ribose-1-(2-N-acetamidomethyl-phosphonate) 5-phosphate (+) (lmm) | day | 56.2 | 6 | <0.0001 | -3 | 1.2 | 39 | 0.22 | 0.91 |
| | treatment | 19.1 | 1 | <0.0001 | 1 | 0.9 | 39 | 0.38 | 0.93 |
| | day:treatment | 29.3 | 6 | <0.0001 | 8 | 0.8 | 39 | 0.41 | 0.94 |
| | | | | | 16 | 4.1 | 39 | <0.0001 | 0.71 |
| | | | | | 22 | 5.8 | 39 | <0.0001 | 0.62 |
| | | | | | 29 | 3.5 | 39 | 0.001 | 0.76 |
| | | | | | 36 | 2.3 | 39 | 0.029 | 0.87 |
| α-D-ribose-1-[N-(phosphonomethyl) glycine] 5-phosphate (-) (lmm) | day | 51.4 | 6 | <0.0001 | -3 | 1 | 32 | 0.32 | 0.93 |
| | treatment | 22.8 | 1 | <0.0001 | 1 | 0.9 | 32 | 0.36 | 0.94 |
| | day:treatment | 57.9 | 6 | <0.0001 | 8 | 2.1 | 32 | 0.042 | 0.87 |
| | | | | | 16 | 7.1 | 32 | <0.0001 | 0.59 |
| | | | | | 22 | 6.3 | 32 | <0.0001 | 0.63 |
| | | | | | 29 | 2.9 | 32 | 0.007 | 0.83 |
| | | | | | 36 | 1.8 | 32 | 0.078 | 0.9 |
| α-D-ribose-1-[N-(phosphonomethyl) glycine] 5-phosphate (+) (lmm) | day | 59.4 | 6 | <0.0001 | -3 | 1 | 31 | 0.34 | 0.93 |
| | treatment | 17 | 1 | <0.0001 | 1 | 1.4 | 31 | 0.18 | 0.9 |
| | day:treatment | 52.6 | 6 | <0.0001 | 8 | 0.2 | 31 | 0.82 | 0.98 |
| | | | | | 16 | 4.9 | 31 | <0.0001 | 0.67 |
| | | | | | 22 | 6.9 | 31 | <0.0001 | 0.55 |
| | | | | | 29 | 2.8 | 31 | 0.008 | 0.8 |
| | | | | | 36 | 2.2 | 31 | 0.033 | 0.86 |
| α-D-ribose-1-[N-(phosphonomethyl) glycine] 5-triphosphate (-) (lmm) | day | 34.1 | 6 | <0.0001 | -3 | 1 | 42 | 0.31 | 0.93 |
| | treatment | 37.5 | 1 | <0.0001 | 1 | 3.9 | 42 | <0.0001 | 0.73 |
| | day:treatment | 26.2 | 6 | <0.0001 | 8 | 1.5 | 42 | 0.15 | 0.9 |
| | | | | | 16 | 6.1 | 42 | <0.0001 | 0.58 |
| | | | | | 22 | 5.2 | 42 | <0.0001 | 0.65 |
| | | | | | 29 | 4 | 42 | <0.0001 | 0.74 |
| | | | | | 36 | 3.4 | 42 | 0.001 | 0.79 |

Table 5: Analysis of Deviance Table (Type II Wald $\chi^2$-tests) for linear mixed effects model to test presence of secondary glyphosate and AMPA breakdown products with percentage ion count (arc sine transformed) of the putatively matched target compound as response. Where the lmm yielded a singular fit, an lm has been fit instead. Pairwise contrasts by day, as estimated by the `emmeans` package for R.

| *Amino acids* | | | | | | | | | |
|---|---|---|---|---|---|---|---|---|---|
| | **lmm/lm** | | | | **Pairwise contrasts** | | | | |
| **Compound** | *fixed effect* | $\chi^2/F$ | *df* | *p* | *contrast* | *t-ratio* | *df* | *p* | *fold change* |
| AA pool (-) (lm) | day | 7.6 | 6,56 | <0.0001 | -3 | 1.5 | 56 | 0.14 | 0.95 |
| | treatment | 4.2 | 1,56 | 0.046 | 1 | 0.3 | 56 | 0.77 | 0.99 |
| | day:treatment | 3.1 | 6,56 | 0.01 | 8 | -2.7 | 56 | 0.008 | 1.11 |
| | | | | | 16 | 1.3 | 56 | 0.18 | 0.95 |
| | | | | | 22 | 2.8 | 56 | 0.006 | 0.89 |
| | | | | | 29 | 0.3 | 56 | 0.73 | 0.99 |
| | | | | | 36 | 1.8 | 56 | 0.076 | 0.93 |
| AA pool (+) (lm) | day | 18.8 | 6,56 | <0.0001 | -3 | 4.1 | 56 | <0.0001 | 0.93 |
| | treatment | 12.8 | 1,56 | 0.001 | 1 | 2.1 | 56 | 0.04 | 0.96 |
| | day:treatment | 5.1 | 6,56 | <0.0001 | 8 | -1.2 | 56 | 0.23 | 1.02 |
| | | | | | 16 | -0.5 | 56 | 0.6 | 1.01 |
| | | | | | 22 | 4.3 | 56 | <0.0001 | 0.92 |
| | | | | | 29 | 1.1 | 56 | 0.27 | 0.98 |

Continued on next page

Table 5 – continued from previous page

| *Amino acids* | | | | | | | | | |
|---|---|---|---|---|---|---|---|---|---|
| | **lmm/lm** | | | | **Pairwise contrasts** | | | | |
| **Compound** | *fixed effect* | $\chi^2/F$ | *df* | *p* | *contrast* | *t-ratio* | *df* | *p* | *fold change* |
| | | | | | 36 | -0.5 | 56 | 0.62 | 1.01 |
| Alanine* (+) (lm) | day | 12.9 | 6,56 | <0.0001 | -3 | 3.9 | 56 | <0.0001 | 0.88 |
| | treatment | 0.9 | 1,56 | 0.34 | 1 | 0.2 | 56 | 0.82 | 0.99 |
| | day:treatment | 10.8 | 6,56 | <0.0001 | 8 | -5.3 | 56 | <0.0001 | 1.18 |
| | | | | | 16 | -4.4 | 56 | <0.0001 | 1.15 |
| | | | | | 22 | 0.7 | 56 | 0.47 | 0.98 |
| | | | | | 29 | 0.5 | 56 | 0.61 | 0.98 |
| | | | | | 36 | 1.7 | 56 | 0.093 | 0.94 |
| Arginine (-) (lm) | day | 13.5 | 6,56 | <0.0001 | -3 | 2.2 | 56 | 0.031 | 0.9 |
| | treatment | 41.9 | 1,56 | <0.0001 | 1 | 1.5 | 56 | 0.14 | 0.91 |
| | day:treatment | 3.8 | 6,56 | 0.003 | 8 | 1.3 | 56 | 0.19 | 0.93 |
| | | | | | 16 | 5.2 | 56 | <0.0001 | 0.73 |
| | | | | | 22 | 5.2 | 56 | <0.0001 | 0.73 |
| | | | | | 29 | 0.8 | 56 | 0.45 | 0.96 |
| | | | | | 36 | 0.9 | 56 | 0.37 | 0.95 |
| Arginine (+) (lm) | day | 11.1 | 6,56 | <0.0001 | -3 | 3.1 | 56 | 0.003 | 0.88 |
| | treatment | 21 | 1,56 | <0.0001 | 1 | -0.3 | 56 | 0.8 | 1.01 |
| | day:treatment | 7 | 6,56 | <0.0001 | 8 | -0.7 | 56 | 0.46 | 1.04 |
| | | | | | 16 | 4 | 56 | <0.0001 | 0.82 |
| | | | | | 22 | 6.1 | 56 | <0.0001 | 0.75 |
| | | | | | 29 | 0.7 | 56 | 0.47 | 0.97 |
| | | | | | 36 | -0.7 | 56 | 0.51 | 1.03 |
| Asparagine* (-) (lm) | day | 3 | 6,56 | 0.012 | -3 | -0.6 | 56 | 0.57 | 1.03 |
| | treatment | 40.3 | 1,56 | <0.0001 | 1 | 1 | 56 | 0.33 | 0.96 |
| | day:treatment | 4.4 | 6,56 | 0.001 | 8 | 1.3 | 56 | 0.2 | 0.95 |
| | | | | | 16 | 5.6 | 56 | <0.0001 | 0.77 |
| | | | | | 22 | 4.4 | 56 | <0.0001 | 0.81 |
| | | | | | 29 | 2.5 | 56 | 0.016 | 0.9 |
| | | | | | 36 | 2.6 | 56 | 0.012 | 0.88 |
| Aspartate (-) (lmm) | day | 113.2 | 6 | <0.0001 | -3 | 3.4 | 53 | 0.001 | 0.86 |
| | treatment | 71.3 | 1 | <0.0001 | 1 | -0.3 | 53 | 0.79 | 1.01 |
| | day:treatment | 197.7 | 6 | <0.0001 | 8 | -11.8 | 53 | <0.0001 | 1.59 |
| | | | | | 16 | -9.6 | 53 | <0.0001 | 1.48 |
| | | | | | 22 | -5.2 | 53 | <0.0001 | 1.26 |
| | | | | | 29 | -4.5 | 53 | <0.0001 | 1.24 |
| | | | | | 36 | -0.1 | 53 | 0.96 | 1 |
| Aspartate (+) (lmm) | day | 37 | 6 | <0.0001 | -3 | 3.3 | 45 | 0.002 | 0.9 |
| | treatment | 2.7 | 1 | 0.1 | 1 | -0.6 | 45 | 0.54 | 1.02 |
| | day:treatment | 40.7 | 6 | <0.0001 | 8 | -3.1 | 45 | 0.003 | 1.1 |
| | | | | | 16 | 1.2 | 45 | 0.25 | 0.97 |
| | | | | | 22 | 3.5 | 45 | 0.001 | 0.9 |
| | | | | | 29 | 1.9 | 45 | 0.066 | 0.94 |
| | | | | | 36 | 0.3 | 45 | 0.79 | 0.99 |
| Cysteine (-) (lm) | day | 13.6 | 6,56 | <0.0001 | -3 | -0.6 | 56 | 0.56 | 1.12 |
| | treatment | 71.2 | 1,56 | <0.0001 | 1 | -0.3 | 56 | 0.77 | 1.07 |
| | day:treatment | 13.1 | 6,56 | <0.0001 | 8 | -2.2 | 56 | 0.03 | 1.48 |
| | | | | | 16 | -9.5 | 56 | <0.0001 | 3.04 |
| | | | | | 22 | -7.1 | 56 | <0.0001 | 2.5 |
| | | | | | 29 | -1.6 | 56 | 0.1 | 1.36 |
| | | | | | 36 | -1 | 56 | 0.34 | 1.2 |
| Cysteine (+) (lm) | day | 6.2 | 6,56 | <0.0001 | -3 | 0.3 | 56 | 0.78 | 0.98 |
| | treatment | 35 | 1,56 | <0.0001 | 1 | 1.2 | 56 | 0.23 | 0.92 |
| | day:treatment | 11.6 | 6,56 | <0.0001 | 8 | -2 | 56 | 0.052 | 1.14 |
| | | | | | 16 | -8.4 | 56 | <0.0001 | 1.65 |
| | | | | | 22 | -5.2 | 56 | <0.0001 | 1.4 |
| | | | | | 29 | -0.8 | 56 | 0.44 | 1.06 |
| | | | | | 36 | -0.8 | 56 | 0.42 | 1.06 |
| Glutamate (-) (lm) | day | 22.9 | 6,56 | <0.0001 | -3 | 0 | 56 | 0.98 | 1 |
| | treatment | 5.4 | 1,56 | 0.023 | 1 | 0.1 | 56 | 0.88 | 0.99 |
| | day:treatment | 7.6 | 6,56 | <0.0001 | 8 | -6.7 | 56 | <0.0001 | 1.36 |
| | | | | | 16 | 0.2 | 56 | 0.87 | 0.99 |
| | | | | | 22 | 2 | 56 | 0.05 | 0.9 |
| | | | | | 29 | -1 | 56 | 0.32 | 1.05 |
| | | | | | 36 | -0.8 | 56 | 0.45 | 1.04 |
| Glutamate (+) (lm) | day | 31 | 6,56 | <0.0001 | -3 | 0.1 | 56 | 0.92 | 1 |
| | treatment | 20.1 | 1,56 | <0.0001 | 1 | 1.1 | 56 | 0.26 | 0.95 |
| | day:treatment | 13.1 | 6,56 | <0.0001 | 8 | -8.6 | 56 | <0.0001 | 1.31 |
| | | | | | 16 | -4.2 | 56 | <0.0001 | 1.15 |
| | | | | | 22 | 1.9 | 56 | 0.058 | 0.93 |
| | | | | | 29 | -1 | 56 | 0.32 | 1.04 |
| | | | | | 36 | -1.2 | 56 | 0.22 | 1.05 |
| Glutamine* (-) (lm) | day | 5.3 | 6,56 | <0.0001 | -3 | 1.3 | 56 | 0.2 | 0.94 |
| | treatment | 182 | 1,56 | <0.0001 | 1 | 0 | 56 | 1 | 1 |
| | day:treatment | 22 | 6,56 | <0.0001 | 8 | 6.6 | 56 | <0.0001 | 0.71 |
| | | | | | 16 | 11.9 | 56 | <0.0001 | 0.51 |
| | | | | | 22 | 10.6 | 56 | <0.0001 | 0.56 |
| | | | | | 29 | 3.3 | 56 | 0.001 | 0.85 |

Continued on next page

Table 5 – continued from previous page

| *Amino acids* | | | | | | | | | |
|---|---|---|---|---|---|---|---|---|---|
| | **lmm/lm** | | | | **Pairwise contrasts** | | | | |
| **Compound** | *fixed effect* | $\chi^2/F$ | *df* | *p* | *contrast* | *t-ratio* | *df* | *p* | *fold change* |
| | | | | | 36 | 2 | 56 | 0.051 | 0.9 |
| Glutamine* (+) (lm) | day | 14.5 | 6,56 | <0.0001 | -3 | -0.2 | 56 | 0.83 | 1.01 |
| | treatment | 48.2 | 1,56 | <0.0001 | 1 | 4.1 | 56 | <0.0001 | 0.89 |
| | day:treatment | 6.9 | 6,56 | <0.0001 | 8 | 7 | 56 | <0.0001 | 0.83 |
| | | | | | 16 | 0.2 | 56 | 0.81 | 0.99 |
| | | | | | 22 | 3.5 | 56 | 0.001 | 0.92 |
| | | | | | 29 | 3.3 | 56 | 0.002 | 0.92 |
| | | | | | 36 | 0.4 | 56 | 0.7 | 0.99 |
| Histidine (-) (lmm) | day | 73.4 | 6 | <0.0001 | -3 | 2.7 | 54 | 0.008 | 0.9 |
| | treatment | 4.2 | 1 | 0.041 | 1 | 0.5 | 54 | 0.59 | 0.98 |
| | day:treatment | 27.7 | 6 | <0.0001 | 8 | -2.7 | 54 | 0.01 | 1.11 |
| | | | | | 16 | 0.1 | 54 | 0.9 | 0.99 |
| | | | | | 22 | 2 | 54 | 0.046 | 0.91 |
| | | | | | 29 | 0.2 | 54 | 0.84 | 0.99 |
| | | | | | 36 | 3.5 | 54 | 0.001 | 0.87 |
| Isoleucine/Leucine (-) (lmm) | day | 58.4 | 6 | <0.0001 | -3 | 1.8 | 56 | 0.075 | 0.89 |
| | treatment | 2.1 | 1 | 0.15 | 1 | 0 | 56 | 0.98 | 1 |
| | day:treatment | 12.4 | 6 | 0.054 | 8 | -2.1 | 56 | 0.04 | 1.16 |
| | | | | | 16 | -0.1 | 56 | 0.91 | 1.01 |
| | | | | | 22 | 1.5 | 56 | 0.13 | 0.87 |
| | | | | | 29 | 0.7 | 56 | 0.48 | 0.94 |
| | | | | | 36 | 2 | 56 | 0.055 | 0.85 |
| Isoleucine/Leucine (+) (lmm) | day | 177.7 | 6 | <0.0001 | -3 | 3.7 | 54 | 0.001 | 0.91 |
| | treatment | 3 | 1 | 0.084 | 1 | 0.9 | 54 | 0.36 | 0.97 |
| | day:treatment | 78.2 | 6 | <0.0001 | 8 | -4.8 | 54 | <0.0001 | 1.15 |
| | | | | | 16 | -6.2 | 54 | <0.0001 | 1.19 |
| | | | | | 22 | -0.3 | 54 | 0.77 | 1.01 |
| | | | | | 29 | 0.2 | 54 | 0.86 | 0.99 |
| | | | | | 36 | 0.9 | 54 | 0.37 | 0.97 |
| Lysine (-) (lmm) | day | 49 | 6 | <0.0001 | -3 | 1 | 55 | 0.32 | 0.95 |
| | treatment | 0.1 | 1 | 0.8 | 1 | 1.3 | 55 | 0.22 | 0.93 |
| | day:treatment | 21.8 | 6 | 0.001 | 8 | -3.8 | 55 | <0.0001 | 1.19 |
| | | | | | 16 | 0.7 | 55 | 0.48 | 0.97 |
| | | | | | 22 | 1 | 55 | 0.32 | 0.95 |
| | | | | | 29 | -0.5 | 55 | 0.6 | 1.03 |
| | | | | | 36 | 1.2 | 55 | 0.25 | 0.94 |
| Lysine (+) (lmm) | day | 58.3 | 6 | <0.0001 | -3 | 0 | 56 | 0.99 | 1 |
| | treatment | 53.8 | 1 | <0.0001 | 1 | 2.9 | 56 | 0.005 | 0.93 |
| | day:treatment | 100.2 | 6 | <0.0001 | 8 | -4.9 | 56 | <0.0001 | 1.12 |
| | | | | | 16 | -9.8 | 56 | <0.0001 | 1.24 |
| | | | | | 22 | -4.3 | 56 | <0.0001 | 1.1 |
| | | | | | 29 | -2.8 | 56 | 0.008 | 1.07 |
| | | | | | 36 | -2.5 | 56 | 0.016 | 1.06 |
| Methionine (-) (lm) | day | 5.2 | 6,56 | <0.0001 | -3 | 0.4 | 56 | 0.7 | 0.98 |
| | treatment | 8.4 | 1,56 | 0.005 | 1 | -2.8 | 56 | 0.008 | 1.17 |
| | day:treatment | 4.1 | 6,56 | 0.002 | 8 | -3.5 | 56 | 0.001 | 1.21 |
| | | | | | 16 | -1.4 | 56 | 0.17 | 1.08 |
| | | | | | 22 | 0 | 56 | 0.97 | 1 |
| | | | | | 29 | -2.5 | 56 | 0.015 | 1.15 |
| | | | | | 36 | 2.1 | 56 | 0.036 | 0.9 |
| Methionine (+) (lmm) | day | 61 | 6 | <0.0001 | -3 | -0.6 | 52 | 0.56 | 1.01 |
| | treatment | 73.7 | 1 | <0.0001 | 1 | 5.3 | 52 | <0.0001 | 0.91 |
| | day:treatment | 43.1 | 6 | <0.0001 | 8 | 7.2 | 52 | <0.0001 | 0.87 |
| | | | | | 16 | 6 | 52 | <0.0001 | 0.89 |
| | | | | | 22 | 4.5 | 52 | <0.0001 | 0.92 |
| | | | | | 29 | 3.8 | 52 | <0.0001 | 0.93 |
| | | | | | 36 | 3 | 52 | 0.004 | 0.95 |
| Phenylalanine (-) (lmm) | day | 11 | 6 | 0.089 | -3 | 0.5 | 56 | 0.64 | 0.97 |
| | treatment | 2.8 | 1 | 0.097 | 1 | -0.8 | 56 | 0.42 | 1.07 |
| | day:treatment | 12 | 6 | 0.062 | 8 | -1.8 | 56 | 0.085 | 1.13 |
| | | | | | 16 | -1.6 | 56 | 0.11 | 1.12 |
| | | | | | 22 | -1.7 | 56 | 0.093 | 1.13 |
| | | | | | 29 | -1.1 | 56 | 0.29 | 1.08 |
| | | | | | 36 | 2 | 56 | 0.045 | 0.86 |
| Phenylalanine (+) (lmm) | day | 140.1 | 6 | <0.0001 | -3 | 3.5 | 55 | 0.001 | 0.95 |
| | treatment | 16.2 | 1 | <0.0001 | 1 | 3.2 | 55 | 0.002 | 0.95 |
| | day:treatment | 191.2 | 6 | <0.0001 | 8 | -0.1 | 55 | 0.9 | 1 |
| | | | | | 16 | -10.8 | 55 | <0.0001 | 1.18 |
| | | | | | 22 | -7.9 | 55 | <0.0001 | 1.13 |
| | | | | | 29 | -0.8 | 55 | 0.42 | 1.01 |
| | | | | | 36 | 0.4 | 55 | 0.66 | 0.99 |
| Proline (+) (lm) | day | 24.4 | 6,56 | <0.0001 | -3 | 5.6 | 56 | <0.0001 | 0.87 |
| | treatment | 62.1 | 1,56 | <0.0001 | 1 | 1.5 | 56 | 0.14 | 0.96 |
| | day:treatment | 6 | 6,56 | <0.0001 | 8 | 1 | 56 | 0.31 | 0.97 |
| | | | | | 16 | 3.2 | 56 | 0.002 | 0.91 |
| | | | | | 22 | 7 | 56 | <0.0001 | 0.81 |
| | | | | | 29 | 2.1 | 56 | 0.044 | 0.94 |

Continued on next page

Table 5 – continued from previous page

| *Amino acids* | | | | | | | | | |
|---|---|---|---|---|---|---|---|---|---|
| | **lmm/lm** | | | | **Pairwise contrasts** | | | | |
| **Compound** | *fixed effect* | $\chi^2/F$ | *df* | *p* | *contrast* | *t-ratio* | *df* | *p* | *fold change* |
| | | | | | 36 | 0.5 | 56 | 0.61 | 0.99 |
| Serine (-) (lm) | day | 8.7 | 6,56 | <0.0001 | -3 | 0.6 | 56 | 0.54 | 0.97 |
| | treatment | 4.3 | 1,56 | 0.042 | 1 | -0.3 | 56 | 0.76 | 1.01 |
| | day:treatment | 7.5 | 6,56 | <0.0001 | 8 | -4.4 | 56 | <0.0001 | 1.19 |
| | | | | | 16 | 2.4 | 56 | 0.02 | 0.9 |
| | | | | | 22 | 4.1 | 56 | <0.0001 | 0.82 |
| | | | | | 29 | 0.7 | 56 | 0.48 | 0.97 |
| | | | | | 36 | 2.4 | 56 | 0.021 | 0.9 |
| Threonine (-) (lmm) | day | 49.9 | 6 | <0.0001 | -3 | 0.7 | 56 | 0.47 | 0.96 |
| | treatment | 4.8 | 1 | 0.029 | 1 | -0.7 | 56 | 0.48 | 1.04 |
| | day:treatment | 31.1 | 6 | <0.0001 | 8 | -3.1 | 56 | 0.003 | 1.16 |
| | | | | | 16 | 1.3 | 56 | 0.21 | 0.94 |
| | | | | | 22 | 3.4 | 56 | 0.001 | 0.83 |
| | | | | | 29 | 1.3 | 56 | 0.19 | 0.93 |
| | | | | | 36 | 3.2 | 56 | 0.002 | 0.85 |
| Tryptophan (-) (lmm) | day | 216.3 | 6 | <0.0001 | -3 | 2.8 | 53 | 0.006 | 0.93 |
| | treatment | 51.9 | 1 | <0.0001 | 1 | 4.4 | 53 | <0.0001 | 0.88 |
| | day:treatment | 13.6 | 6 | 0.034 | 8 | 1.2 | 53 | 0.23 | 0.96 |
| | | | | | 16 | 4.4 | 53 | <0.0001 | 0.87 |
| | | | | | 22 | 5.3 | 53 | <0.0001 | 0.84 |
| | | | | | 29 | 2.1 | 53 | 0.043 | 0.93 |
| | | | | | 36 | 3.8 | 53 | <0.0001 | 0.89 |
| Tryptophan (+) (lmm) | day | 90.9 | 6 | <0.0001 | -3 | 1.8 | 56 | 0.075 | 0.96 |
| | treatment | 10.2 | 1 | 0.001 | 1 | 1.2 | 56 | 0.22 | 0.97 |
| | day:treatment | 5.9 | 6 | 0.44 | 8 | 2.8 | 56 | 0.007 | 0.93 |
| | | | | | 16 | 1.8 | 56 | 0.084 | 0.95 |
| | | | | | 22 | -0.1 | 56 | 0.95 | 1 |
| | | | | | 29 | 1.4 | 56 | 0.17 | 0.96 |
| | | | | | 36 | 0.2 | 56 | 0.81 | 0.99 |
| Tyrosine (-) (lm) | day | 5.3 | 6,56 | <0.0001 | -3 | 1 | 56 | 0.3 | 0.95 |
| | treatment | 0.5 | 1,56 | 0.47 | 1 | -1.4 | 56 | 0.17 | 1.08 |
| | day:treatment | 1.7 | 6,56 | 0.14 | 8 | -1.7 | 56 | 0.098 | 1.09 |
| | | | | | 16 | -0.6 | 56 | 0.54 | 1.04 |
| | | | | | 22 | -0.2 | 56 | 0.81 | 1.01 |
| | | | | | 29 | -0.9 | 56 | 0.36 | 1.06 |
| | | | | | 36 | 1.9 | 56 | 0.062 | 0.9 |
| Tyrosine (+) (lmm) | day | 93.7 | 6 | <0.0001 | -3 | 2.4 | 54 | 0.019 | 0.96 |
| | treatment | 2.5 | 1 | 0.11 | 1 | 2.3 | 54 | 0.026 | 0.96 |
| | day:treatment | 19.3 | 6 | 0.004 | 8 | 2.1 | 54 | 0.039 | 0.96 |
| | | | | | 16 | -2 | 54 | 0.045 | 1.04 |
| | | | | | 22 | -0.4 | 54 | 0.67 | 1.01 |
| | | | | | 29 | -0.5 | 54 | 0.6 | 1.01 |
| | | | | | 36 | 1.2 | 54 | 0.25 | 0.98 |
| Valine (-) (lmm) | day | 242.3 | 6 | <0.0001 | -3 | 4 | 55 | <0.0001 | 0.92 |
| | treatment | 85.7 | 1 | <0.0001 | 1 | 5.3 | 55 | <0.0001 | 0.88 |
| | day:treatment | 13.9 | 6 | 0.031 | 8 | 1.9 | 55 | 0.069 | 0.96 |
| | | | | | 16 | 4.8 | 55 | <0.0001 | 0.88 |
| | | | | | 22 | 6 | 55 | <0.0001 | 0.85 |
| | | | | | 29 | 2.6 | 55 | 0.014 | 0.93 |
| | | | | | 36 | 4 | 55 | <0.0001 | 0.9 |

Table 6: m/z peaks and their putatively matched compounds, accurate masses, expected masses as given adducts, chemical formulas and expected roll in the cell metabolome. Compounds with the same monoisotopic mass are listed together.

| **m/z peak (mode)** | **Compound** | **Structure** | **Accurate mass** | **Expected mass and ion** | **Class** |
|---|---|---|---|---|---|
| *Pre-resistance* | | | | | |
| 242.073 (-) | mycothiol | $C_{17}H_{30}N_2O_{12}S$ | 486.1519 | 486.1616 $[M-2H]^{2-}$ | thiol |
| | cytidine | $C_9H_{13}N_3O_5$ | 243.0855 | 243.0808 $[M-H]^-$ | pyrimidine ribonucleoside |
| 269.2311 (-) | 9,9'-di-*cis*-ζ-carotene | $C_{40}H_{60}$ | 540.4695 | 540.4778 $[M-2H]^{2-}$ | carotenoid |
| | 9,15,9'-tri-*cis*-ζ-carotene | $C_{40}H_{60}$ | 540.4695 | 540.4778 $[M-2H]^{2-}$ | carotenoid |
| | all-*trans*-ζ-carotene | $C_{40}H_60$ | 540.4695019 | 540.4778 $[M-2H]^{2-}$ | carotenoid |
| 297.263 (-) | pristanate | $C_{19}H_{37}O_2$ | 298.2871805 | 298.2708 $[M-H]^-$ | fatty acid |
| 301.1833 (-) | carlactone | $C_{19}H_{26}O_3$ | 302.1881947 | 302.1911 $[M-H]^-$ | lactone |
| 303.2167 (+) | linoleate | $C_{18}H_{31}O_2$ | 280.2402303 | 280.2269 $[M+Na]^+$ | fatty acid |
| | all-*trans*-3-hydroxyretinol | $C_{20}H_{30}O_2$ | 302.2245802 | 302.2089 $[M+H]^+$ | retinol |
| Continued on next page | | | | | |

Table 6 – continued from previous page

| m/z peak (mode) | Compound | Structure | Accurate mass | Expected mass and ion | Class |
|---|---|---|---|---|---|
| | 11-*cis*-3-hydroxyretinol | $C_{20}H_{30}O_2$ | 302.2245802 | 302.2089 $[M+H]^+$ | retinol |
| 311.1896 (-) | 1-18:3-2-18:3-digalactosyldiacyl-glycerol | $C_{51}H_{84}O_{15}$ | 936.581022 | 936.5922 $[M-3H]^{3-}$ | galactolipid |
| 346.0795 (-) | 3-oxo-2-(*cis*-2'-pentenyl)-cyclopentane-1-octa-2-enoyl-CoA | $C_{39}H_{58}N_7O_{18}P_3S$ | 1041.308488 | 1041.2619 $[M-3H]^{3-}$ | CoA activated ester |
| 431.3313 (-) | calcitetrol | $C_{27}H_{44}O_4$ | 432.3239599 | 432.3391 $[M-H]^-$ | vitamin d3 metabolite |
| | ubiquinol-10 | $C_{59}H_{92}O_4$ | 864.6995614 | 864.6782 $[M-2H]^{2-}$ | electron-transfer quinol |
| 459.3297 (+) | demethylphylloquinone | $C_{30}H_{44}O_2$ | 436.3341307 | 436.3399 $[M+Na]^+$ | electron-transfer quinone |
| | 1-16:0-2-18:2-digalactosyldiacyl-glycerol | $C_{49}H_{88}O_{15}$ | 916.6123222 | 916.6438 $[M+2H]^{2+}$ | galactolipid |
| | 1-18:2-2-16:0-digalactosyldiacyl-glycerol | $C_{49}H_{88}O_{15}$ | 916.6123222 | 916.6438 $[M+2H]^{2+}$ | galactolipid |
| 579.4701 (+) | 3-heptaprenyl-4-hydroxybenzoate | $C_{42}H_{61}O_3$ | 614.4698959 | 614.4833 $[M+H-2H2O]^+$ | isoprenoid lipid |
| 597.4818 (+) | 3-heptaprenyl-4-hydroxybenzoate | $C_{42}H_{61}O_3$ | 614.4698959 | 614.4845 $[M+H-H2O]^+$ | isoprenoid lipid |
| 601.4983 (+) | 2-methoxy-6-(all-*trans*-heptaprenyl)phenol | $C_{42}H_{64}O_2$ | 600.4906313 | 600.4905 $[M+H]^+$ | aromatic ether |
| | demethylmenaquinol-7 | $C_{45}H_{64}O_2$ | 636.4906313 | 636.5115 $[M+H-2H2O]^+$ | electron-transfer quinol |
| 602.5017 (+) | an N-(stearoyl)-sphinga-4,8-dienine | $C_{36}H_{69}NO_3$ | 563.5277451 | 563.538 $[M+K]^+$ | ceramide |
| 694.4306 (+) | 2,3-bis[(3R)-3-hydroxymyristoyl]-$\alpha$-D-glucosaminyl 1-phosphate | $C_{34}H_{64}NO_{12}P$ | 711.4322631 | 711.4333 $[M+H-H2O]^+$ | glycolipid |
| 704.6229 (+) | 1-16:0-2-18:1-diacylglycerol-trimethylhomoserine | $C_{44}H_{83}NO_7$ | 739.6326041 | 739.6361 $[M+H-2H2O]^+$ | lipid |
| 732.6665 (+) | a 4-hydroxysphing-8-enine-26:0 ceramide | $C_{44}H_{87}NO_4$ | 693.6635103 | 693.7028 $[M+K]^+$ | ceramide |
| 733.6681 (+) | 2,3-bis-O-phytanyl-sn-glycerol 1-phosphate | $C_{43}H_{87}O_6P$ | 732.6396771 | 732.6603 $[M+H]^+$ | phsopholipid |
| | a plastoquinol | $C_{53}H_{82}O_2$ | 750.6314819 | 750.6708 $[M+H-H2O]^+$ | electron-transfer quinol |
| | plastoquinol-9 | $C_{53}H_{82}O_2$ | 750.6314819 | 750.6708 $[M+H-H2O]^+$ | electron-transfer quinol |
| 767.5632 (+) | ubiquinol-8 | $C_{49}H_{76}O_4$ | 728.5743609 | 728.5995 $[M+K]^+$ | electron-transfer quinol |
| | 3,4-dihydroxy-5-all-*trans*-nonaprenylbenzoate | $C_{52}H_{77}O_4$ | 766.590011 | 766.5554 $[M+H]^+$ | aromatic compound |
| 768.5688 (+) | 1-18:0-2-18:1-phosphatidyl-ethanolamine | $C_{41}H_{80}NO_8P$ | 745.5621551 | 745.579 $[M+Na]^+$ | phosphatidyl-ethanolamine |
| 777.6983 (+) | thyroxine | $C_{15}H_{11}NO_4I_4$ | 776.6866798 | 776.6905 $[M+H]^+$ | hormone |
| 784.5426 (+) | 1-18:0-2-18:1-phosphatidyl-ethanolamine | $C_{41}H_{80}NO_8P$ | 745.5621551 | 745.5789 $[M+K]^+$ | phosphatidyl-ethanolamine |
| 793.5764 (-) | ubiquinone-9 | $C_{54}H_{82}O_4$ | 794.6213111 | 794.5842 $[M-H]^-$ | electron-transfer quinone |
| 849.5403 (+) | pheophytin b | $C_{55}H_{72}N_4O_6$ | 884.5451861 | 884.5535 $[M+H-2H2O]^+$ | pheophytin |
| 857.5345 (-) | all-*trans*-decaprenyl diphosphate | $C_{50}H_{81}O_7P_2$ | 858.5692281 | 858.5423 $[M-H]^-$ | isoprenoid lipid |
| | mono-*trans*,octa-*cis*-decaprenyl diphosphate | $C_{50}H_{81}O_7P_2$ | 858.5692281 | 858.5423 $[M-H]^-$ | isoprenoid lipid |
| 905.5121 (-) | chlorophyll b | $C_{55}H_{70}N_4O_6Mg$ | 906.5145779 | 906.5199 $[M-H]^-$ | chlorophyll pigment |
| | | *Post-resistance* | | | |
| 89.0332 (-) | coniferyl alcohol | $C_{10}H_{12}O_3$ | 180.0786443 | 180.082 $[M-2H]^{2-}$ | phenylpropanoid |
| 277.236 (+) | $\beta$-cryptoxanthin | $C_{40}H_{56}O$ | 552.4331164 | 552.4564 $[M+2H]^{2+}$ | carotenoid |
| | 3,4-didehydrorhodopin | $C_{40}H_{56}O$ | 552.4331164 | 552.4564 $[M+2H]^{2+}$ | carotenoid |

Continued on next page

Table 6 – continued from previous page

| m/z peak (mode) | Compound | Structure | Accurate mass | Expected mass and ion | Class |
|---|---|---|---|---|---|
| | zeinoxanthin | $C_{40}H_{56}O$ | 552.4331164 | 552.4564 $[M+2H]^{2+}$ | carotenoid |
| | α-cryptoxanthin | $C_{40}H_{56}O$ | 552.4331164 | 552.4564 $[M+2H]^{2+}$ | carotenoid |
| 277.2364 (-) | 6-methoxy-3-methyl-2-all-*trans*-decaprenyl-1,4-benzoquinol | $C_{58}H_{90}O_3$ | 834.6889968 | 834.7326 $[M-3H]^{3-}$ | quinol |
| 279.2516 (+) | 6-methoxy-3-methyl-2-all-*trans*-decaprenyl-1,4-benzoquinol | $C_{58}H_{90}O_3$ | 834.6889968 | 834.7314 $[M+3H]^{3+}$ | quinol |
| 281.1948 (-) | canthaxanthin | $C_{40}H_{52}O_2$ | 564.3967309 | 564.4052 $[M-2H]^{2-}$ | carotenoid |
| 291.2175 (-) | androstan-3α,17β-diol | $C_{19}H_{32}O_2$ | 292.2402303 | 292.2253 $[M-H]^{-}$ | sterol |
| 292.2202 (-) | 3-(all-*trans*-heptaprenyl)benzene-1,2-diol | $C_{41}H_{62}O_2$ | 586.4749812 | 586.456 $[M-2H]^{2-}$ | |
| 348.1044 (+) | 3-oxo-2-(*cis*-2'-pentenyl)-cyclopentane-1-octa-2-enoyl-CoA | $C_{39}H_{58}N_7O_{18}P_3S$ | 1041.308488 | 1041.2898 $[M+3H]^{3+}$ | CoA activated ester |
| 451.1547 (-) | 3'-O-demethyl-staurosporine | $C_{27}H_{25}N_4O_3$ | 452.1848407 | 452.1625 $[M-H]^{-}$ | azole |
| 455.1676 (+) | coenzyme F430 | $C_{42}H_{46}N_6O_{13}Ni$ | 908.3102338 | 908.3196 $[M+2H]^{2+}$ | enzyme cofactor |
| 556.3296 (-) | Und-PP-MurNAc-L-Ala-γ-D-Glu-L-Lys-D-Ala-D-Ala | $C_{86}H_{140}N_7O_{21}P_2$ | 1671.981228 | 1672.0122 $[M-3H]^{3-}$ | glycoconjugate |
| 609.3424 (+) | presqualene diphosphate | $C_{30}H_{49}O_7P_2$ | 586.318827 | 586.3526 $[M+Na]^{+}$ | isoprenoid lipid |
| | all-*trans*-hexaprenyl diphosphate | $C_{30}H_{49}O_7P_2$ | 586.318827 | 586.3526 $[M+Na]^{+}$ | isoprenoid lipid |
| 621.3825 (+) | menaquinol-6 | $C_{41}H_{58}O_2$ | 582.4436811 | 582.4188 $[M+K]^{+}$ | electron-transfer quinol |
| | adonixanthin | $C_{40}H_{54}O_3$ | 582.4072956 | 582.4188 $[M+K]^{+}$ | carotenoid |
| | hydroxyspirilloxanthin | $C_{41}H_{58}O_2$ | 582.4436811 | 582.4188 $[M+K]^{+}$ | carotenoid |
| 623.3743 (+) | caloxanthin | $C_{40}H_{56}O_3$ | 584.4229457 | 584.4106 $[M+K]^{+}$ | carotenoid |
| | antheraxanthin | $C_{40}H_{56}O_3$ | 584.4229457 | 584.4106 $[M+K]^{+}$ | carotenoid |
| | rhodovibrin | $C_{41}H_{60}O_2$ | 584.4593312 | 584.4506 $[M+K]^{+}$ | carotenoid |
| | caloxanthin | $C_{40}H_{56}O_3$ | 584.4229457 | 584.4506 $[M+K]^{+}$ | carotenoid |
| | *trans*-neoxanthin | $C_{40}H_{56}O_4$ | 600.4178603 | 600.3845 $[M+Na]^{+}$ | carotenoid |
| | 9-*cis*-violaxanthin | $C_{40}H_{56}O_4$ | 600.4178603 | 600.3845 $[M+Na]^{+}$ | carotenoid |
| | violaxanthin | $C_{40}H_{56}O_4$ | 600.4178603 | 600.3845 $[M+Na]^{+}$ | carotenoid |
| | 9'-*cis*-neoxanthin | $C_{40}H_{56}O_4$ | 600.4178603 | 600.3845 $[M+Na]^{+}$ | carotenoid |
| | nostoxanthin | $C_{40}H_{56}O_4$ | 600.4178603 | 600.3845 $[M+Na]^{+}$ | carotenoid |
| 625.3405 (+) | coproporphyrinogen III | $C_{36}H_{40}N_4O_8$ | 660.3159144 | 660.3537 $[M+H-2H2O]^{+}$ | porphyrinogen |
| 627.2585 (-) | primary fluorescent chlorophyll catabolite | $C_{35}H_{38}N_4O_7$ | 628.2896997 | 628.2663 $[M-H]^{-}$ | bilin pigment |
| 721.5547 (-) | 3-(all-*trans*-nonaprenyl)benzene-1,2-diol | $C_{51}H_{78}O_2$ | 722.6001817 | 722.5625 $[M-H]^{-}$ | |
| 747.571 (-) | a plastoquinone | $C_{53}H_{80}O_2$ | 748.6158318 | 748.5788 $[M-H]^{-}$ | electron-transfer quinone |
| 759.688 (+) | thyroxine | $C_{15}H_{11}NO_4I_4$ | 776.6866798 | 776.6907 $[M+H-H2O]^{+}$ | hormone |
| | | | *Persistent* | | |
| 157.0178 (-) | (S)-dihydroorotate | $C_5H_5N_2O_4$ | 158.0327567 | 158.0256 $[M-H]^{-}$ | monocarboxylic acid anion |
| | 3-hydroxy-*cis*,*cis*-muconate | $C_6H_4O_5$ | 158.0215233 | 159.0256 $[M-H]^{-}$ | fatty acid |
| | 2-maleylacetate | $C_6H_4O_5$ | 158.0215233 | 160.0256 $[M-H]^{-}$ | beta-keto acid |
| | (2Z,4E)-2-hydroxyhexa-2,4-dienedioate | $C_6H_4O_5$ | 158.0215233 | 161.0256 $[M-H]^{-}$ | |

Continued on next page

Table 6 – continued from previous page

| m/z peak (mode) | Compound | Structure | Accurate mass | Expected mass and ion | Class |
|---|---|---|---|---|---|
| | (3E)-2-oxohex-3-enedioate | $C_6H_4O_5$ | 158.0215233 | 162.0256 [M-H]$^-$ | carboxylate |
| 266.0025 (-) | 2-[(2R,5Z)-2-carboxy-4-methylthiazol-5(2H)-ylidene]ethyl phosphate | $C_7H_7NO_6PS1$ | 266.9966443 | 267.0103 [M-H]$^-$ | azole |
| | 2-(2-carboxy-4-methylthiazol-5-yl)ethyl phosphate | $C_7H_7NO_6PS2$ | 266.9966443 | 267.0103 [M-H]$^-$ | azole |
| 496.4186 (+) | menaquinol-12 | $C_{71}H_{106}O_2$ | 990.8192826 | 990.8216 [M+2H]$^{2+}$ | electron-transfer quinol |
| 615.4999 (+) | 3-heptaprenyl-4-hydroxybenzoate | $C_{42}H_{61}O_3$ | 614.4698959 | 614.4921 [M+H]$^+$ | isoprenoid lipid |
| | menaquinol-7 | $C_{46}H_{66}O_2$ | 650.5062814 | 650.5131 [M+H-2H2O]$^+$ | electron-transfer quinol |
| | | | *Unclassified pattern* | | |
| 317.2156 (+) | 3-oxo-2-(*cis*-2'-pentenyl)-cyclopentane-1-octanoate | $C_{18}H_{29}O_3$ | 294.2194948 | 294.2258 [M+Na]$^+$ | fatty acid |
| | thromboxane A2 | $C_{20}H_{31}O_5$ | 352.2249741 | 352.2288 [M+H-2H2O]$^+$ | fatty acid |
| | prostaglandin D2 | $C_{20}H_{31}O_5$ | 352.2249741 | 352.2288 [M+H-2H2O]$^+$ | fatty acid |
| | (13E)-11-$\alpha$-hydroxy-9,15-dioxoprost-13-enoate | $C_{20}H_{31}O_5$ | 352.2249741 | 352.2288 [M+H-2H2O]$^+$ | fatty acid |
| | prostaglandin-H2 | $C_{20}H_{31}O_5$ | 352.2249741 | 352.2288 [M+H-2H2O]$^+$ | fatty acid |
| | prostaglandin E2 | $C_{20}H_{31}O_5$ | 352.2249741 | 352.2288 [M+H-2H2O]$^+$ | fatty acid |

Table 7: Analysis of Deviance Table (Type II Wald $\chi^2$-tests) for linear mixed effects model for all m/z peaks of interest with pre-resistance pattern from exploratory analysis with percentage ion count (arc sine transformed) of the m/z peak as response. Where the lmm yielded a singular fit, an lm has been fit instead. P-values have been adjusted using the Benjamini-Hochberg procedure. Pairwise contrasts by day, as estimated by the `emmeans` package for R.

| **Pre-resistance pattern** | | | | | | | | | |
|---|---|---|---|---|---|---|---|---|---|
| | **lmm/lm** | | | | **Pairwise contrasts** | | | | |
| **m/z peak** | *fixed effect* | $\chi^2/F$ | *df* | *adj. p* | *contrast* | *t-ratio* | *df* | *p* | *fold change* |
| 146.978 (-) (lmm) | day | 59.83 | 6 | <0.0001 | -3 | -0.3 | 53 | 0.74 | 1.06 |
| | treatment | 8.4 | 1 | 0.005 | 1 | 0.2 | 53 | 0.83 | 0.97 |
| | day:treatment | 11.05 | 6 | 0.094 | 8 | 0.8 | 53 | 0.41 | 0.89 |
| | | | | | 16 | 2.4 | 53 | 0.02 | 0.69 |
| | | | | | 22 | 3.4 | 53 | 0.001 | 0.62 |
| | | | | | 29 | 1.8 | 53 | 0.083 | 0.82 |
| | | | | | 36 | 1.2 | 53 | 0.23 | 0.88 |
| 158.9903 (-) (lmm) | day | 83.14 | 6 | <0.0001 | -3 | -1.6 | 54 | 0.13 | 1.06 |
| | treatment | 63.55 | 1 | <0.0001 | 1 | 3.7 | 54 | 0.001 | 0.87 |
| | day:treatment | 78.23 | 6 | <0.0001 | 8 | 7.3 | 54 | <0.0001 | 0.73 |
| | | | | | 16 | 8.6 | 54 | <0.0001 | 0.67 |
| | | | | | 22 | 4 | 54 | <0.0001 | 0.83 |
| | | | | | 29 | 1.7 | 54 | 0.097 | 0.93 |
| | | | | | 36 | 2.1 | 54 | 0.042 | 0.92 |
| 215.0478 (-) (lmm) | day | 76.5 | 6 | <0.0001 | -3 | 0.7 | 45 | 0.51 | 0.95 |
| | treatment | 1.03 | 1 | 0.32 | 1 | -3.5 | 45 | 0.001 | 1.25 |
| | day:treatment | 96.44 | 6 | <0.0001 | 8 | -3.6 | 45 | 0.001 | 1.23 |
| | | | | | 16 | 5.2 | 45 | <0.0001 | 0.69 |
| | | | | | 22 | 5.2 | 45 | <0.0001 | 0.68 |
| | | | | | 29 | 0.4 | 45 | 0.68 | 0.97 |
| | | | | | 36 | -0.4 | 45 | 0.71 | 1.02 |
| 223.1496 (-) (lm) | day | 12.04 | 6,56 | <0.0001 | -3 | -0.8 | 56 | 0.45 | 1.07 |
| | treatment | 22.89 | 1,56 | <0.0001 | 1 | 0.9 | 56 | 0.39 | 0.93 |
| | day:treatment | 6.83 | 6,56 | <0.0001 | 8 | 1.2 | 56 | 0.24 | 0.85 |
| | | | | | 16 | -5.5 | 56 | <0.0001 | 1.73 |
| | | | | | 22 | -4.9 | 56 | <0.0001 | 1.72 |
| | | | | | 29 | -1.9 | 56 | 0.067 | 1.27 |
| | | | | | 36 | -1.6 | 56 | 0.11 | 1.19 |
| 228.0716 (-) (lmm) | day | 175.59 | 6 | <0.0001 | -3 | 0.9 | 46 | 0.38 | 0.96 |
| | treatment | 0.04 | 1 | 0.84 | 1 | -2.7 | 46 | 0.01 | 1.11 |
| | day:treatment | 34.3 | 6 | <0.0001 | 8 | -1.5 | 46 | 0.15 | 1.05 |
| | | | | | 16 | 1.4 | 46 | 0.17 | 0.95 |
| | | | | | 22 | 3.6 | 46 | 0.001 | 0.88 |

Continued on next page

Table 7 – continued from previous page

| Pre-resistance pattern | | | | | | | | | |
|---|---|---|---|---|---|---|---|---|---|
| | lmm/lm | | | | Pairwise contrasts | | | | |
| **m/z peak** | *fixed effect* | $\chi^2/F$ | *df* | *adj. p* | *contrast* | *t-ratio* | *df* | *p* | *fold change* |
| | | | | | 29 | 0.7 | 46 | 0.48 | 0.98 |
| | | | | | 36 | -1.7 | 46 | 0.1 | 1.06 |
| 229.0732 (-) (lmm) | day | 142.24 | 6 | <0.0001 | -3 | 0.7 | 47 | 0.48 | 0.96 |
| | treatment | 2.32 | 1 | 0.14 | 1 | -1.9 | 47 | 0.065 | 1.09 |
| | day:treatment | 32.27 | 6 | <0.0001 | 8 | -0.6 | 47 | 0.52 | 1.03 |
| | | | | | 16 | 2.1 | 47 | 0.039 | 0.91 |
| | | | | | 22 | 4.4 | 47 | <0.0001 | 0.83 |
| | | | | | 29 | 1.7 | 47 | 0.092 | 0.93 |
| | | | | | 36 | -0.5 | 47 | 0.6 | 1.02 |
| 230.0696 (-) (lmm) | day | 139.46 | 6 | <0.0001 | -3 | 0.8 | 47 | 0.42 | 0.96 |
| | treatment | 2.77 | 1 | 0.1 | 1 | -2 | 47 | 0.049 | 1.1 |
| | day:treatment | 34.44 | 6 | <0.0001 | 8 | -0.4 | 47 | 0.7 | 1.02 |
| | | | | | 16 | 2.3 | 47 | 0.024 | 0.91 |
| | | | | | 22 | 4.5 | 47 | <0.0001 | 0.82 |
| | | | | | 29 | 1.7 | 47 | 0.097 | 0.93 |
| | | | | | 36 | -0.6 | 47 | 0.53 | 1.02 |
| 242.073 (-) (lmm) | day | 93.73 | 6 | <0.0001 | -3 | 0.7 | 42 | 0.52 | 0.94 |
| | treatment | 0.06 | 1 | 0.81 | 1 | -4.2 | 42 | <0.0001 | 1.4 |
| | day:treatment | 85.73 | 6 | <0.0001 | 8 | -4 | 42 | <0.0001 | 1.33 |
| | | | | | 16 | 3.1 | 42 | 0.003 | 0.77 |
| | | | | | 22 | 4.4 | 42 | <0.0001 | 0.68 |
| | | | | | 29 | 0.5 | 42 | 0.64 | 0.97 |
| | | | | | 36 | -1.6 | 42 | 0.12 | 1.11 |
| 261.09 (-) (lmm) | day | 124.86 | 6 | <0.0001 | -3 | 0.7 | 52 | 0.46 | 0.96 |
| | treatment | 27.73 | 1 | <0.0001 | 1 | 7.4 | 52 | <0.0001 | 0.66 |
| | day:treatment | 39.28 | 6 | <0.0001 | 8 | 1.6 | 52 | 0.11 | 0.9 |
| | | | | | 16 | 3.4 | 52 | 0.001 | 0.78 |
| | | | | | 22 | 3.2 | 52 | 0.002 | 0.8 |
| | | | | | 29 | 1.2 | 52 | 0.24 | 0.92 |
| | | | | | 36 | 0.6 | 52 | 0.58 | 0.97 |
| 269.2311 (-) (lmm) | day | 365.62 | 6 | <0.0001 | -3 | 0 | 56 | 0.98 | 1 |
| | treatment | 11.54 | 1 | 0.001 | 1 | 3.2 | 56 | 0.002 | 0.83 |
| | day:treatment | 15.8 | 6 | 0.017 | 8 | 3.4 | 56 | 0.001 | 0.79 |
| | | | | | 16 | 1.5 | 56 | 0.15 | 0.9 |
| | | | | | 22 | 1.8 | 56 | 0.072 | 0.86 |
| | | | | | 29 | -0.3 | 56 | 0.78 | 1.02 |
| | | | | | 36 | -0.5 | 56 | 0.64 | 1.03 |
| 286.1728 (+) (lmm) | day | 156.51 | 6 | <0.0001 | -3 | 0.3 | 51 | 0.74 | 0.98 |
| | treatment | 229.86 | 1 | <0.0001 | 1 | 2.3 | 51 | 0.025 | 0.86 |
| | day:treatment | 292.45 | 6 | <0.0001 | 8 | 11 | 51 | <0.0001 | 0.53 |
| | | | | | 16 | 15.1 | 51 | <0.0001 | 0.42 |
| | | | | | 22 | 15.6 | 51 | <0.0001 | 0.46 |
| | | | | | 29 | 7.6 | 51 | <0.0001 | 0.7 |
| | | | | | 36 | 1 | 51 | 0.31 | 0.94 |
| 297.263 (-) (lmm) | day | 366.94 | 6 | <0.0001 | -3 | -0.7 | 54 | 0.46 | 1.03 |
| | treatment | 7.44 | 1 | 0.007 | 1 | 3.2 | 54 | 0.003 | 0.86 |
| | day:treatment | 19.01 | 6 | 0.005 | 8 | 3.7 | 54 | <0.0001 | 0.81 |
| | | | | | 16 | 1 | 54 | 0.33 | 0.95 |
| | | | | | 22 | 1.7 | 54 | 0.091 | 0.9 |
| | | | | | 29 | 0.2 | 54 | 0.85 | 0.99 |
| | | | | | 36 | -0.3 | 54 | 0.8 | 1.01 |
| 301.1833 (-) (lmm) | day | 149 | 6 | <0.0001 | -3 | 0 | 50 | 0.99 | 1 |
| | treatment | 234.39 | 1 | <0.0001 | 1 | 3.6 | 50 | 0.001 | 0.71 |
| | day:treatment | 283.97 | 6 | <0.0001 | 8 | 11.9 | 50 | <0.0001 | 0.38 |
| | | | | | 16 | 15.1 | 50 | <0.0001 | 0.29 |
| | | | | | 22 | 15.3 | 50 | <0.0001 | 0.35 |
| | | | | | 29 | 7.6 | 50 | <0.0001 | 0.63 |
| | | | | | 36 | 1.5 | 50 | 0.13 | 0.88 |
| 302.1732 (-) (lmm) | day | 72.54 | 6 | <0.0001 | -3 | 1.4 | 50 | 0.17 | 0.92 |
| | treatment | 131.79 | 1 | <0.0001 | 1 | 1.9 | 50 | 0.066 | 0.86 |
| | day:treatment | 177.2 | 6 | <0.0001 | 8 | 5.9 | 50 | <0.0001 | 0.69 |
| | | | | | 16 | 12.6 | 50 | <0.0001 | 0.4 |
| | | | | | 22 | 12.8 | 50 | <0.0001 | 0.43 |
| | | | | | 29 | 5.8 | 50 | <0.0001 | 0.7 |
| | | | | | 36 | 1.1 | 50 | 0.29 | 0.92 |
| 303.2167 (+) (lmm) | day | 131.61 | 6 | <0.0001 | -3 | 0 | 54 | 0.97 | 1 |
| | treatment | 170.75 | 1 | <0.0001 | 1 | 1.1 | 54 | 0.3 | 0.92 |
| | day:treatment | 224.97 | 6 | <0.0001 | 8 | 9.5 | 54 | <0.0001 | 0.52 |
| | | | | | 16 | 12.8 | 54 | <0.0001 | 0.42 |
| | | | | | 22 | 13.2 | 54 | <0.0001 | 0.46 |
| | | | | | 29 | 5.6 | 54 | <0.0001 | 0.75 |
| | | | | | 36 | 0 | 54 | 0.98 | 1 |
| 304.1946 (+) (lmm) | day | 55.3 | 6 | <0.0001 | -3 | 1.4 | 45 | 0.17 | 0.91 |
| | treatment | 71.31 | 1 | <0.0001 | 1 | 0.7 | 45 | 0.49 | 0.94 |
| | day:treatment | 176.08 | 6 | <0.0001 | 8 | 3.5 | 45 | 0.001 | 0.79 |
| | | | | | 16 | 11.5 | 45 | <0.0001 | 0.39 |
| | | | | | 22 | 11.5 | 45 | <0.0001 | 0.44 |

Continued on next page

Table 7 – continued from previous page

| Pre-resistance pattern | | | | | | | | | |
|---|---|---|---|---|---|---|---|---|---|
| | lmm/lm | | | | Pairwise contrasts | | | | |
| **m/z peak** | *fixed effect* | $\chi^2/F$ | *df* | *adj. p* | *contrast* | *t-ratio* | *df* | *p* | *fold change* |
| | | | | | 29 | 4.3 | 45 | <0.0001 | 0.76 |
| | | | | | 36 | 0.1 | 45 | 0.89 | 0.99 |
| 311.1896 (-) (lmm) | day | 411.77 | 6 | <0.0001 | -3 | 0.7 | 47 | 0.49 | 0.97 |
| | treatment | 0.03 | 1 | 0.87 | 1 | -7.8 | 47 | <0.0001 | 1.28 |
| | day:treatment | 107.55 | 6 | <0.0001 | 8 | -0.8 | 47 | 0.41 | 1.02 |
| | | | | | 16 | 3.1 | 47 | 0.004 | 0.92 |
| | | | | | 22 | 3.9 | 47 | <0.0001 | 0.89 |
| | | | | | 29 | 1 | 47 | 0.34 | 0.97 |
| | | | | | 36 | 0.7 | 47 | 0.51 | 0.98 |
| 311.2436 (-) (lm) | day | 30.54 | 6,56 | <0.0001 | -3 | 0.8 | 56 | 0.41 | 0.95 |
| | treatment | 4.84 | 1,56 | 0.036 | 1 | 0.7 | 56 | 0.46 | 0.95 |
| | day:treatment | 3.55 | 6,56 | 0.006 | 8 | 1.7 | 56 | 0.097 | 0.86 |
| | | | | | 16 | -3.2 | 56 | 0.002 | 1.3 |
| | | | | | 22 | -2.4 | 56 | 0.022 | 1.24 |
| | | | | | 29 | -1.7 | 56 | 0.1 | 1.19 |
| | | | | | 36 | -1.8 | 56 | 0.071 | 1.18 |
| 346.0795 (-) (lmm) | day | 93.52 | 6 | <0.0001 | -3 | -1.8 | 21 | 0.085 | 1.22 |
| | treatment | 3.16 | 1 | 0.082 | 1 | -0.8 | 21 | 0.4 | 1.08 |
| | day:treatment | 113.62 | 6 | <0.0001 | 8 | -2.4 | 21 | 0.025 | 1.19 |
| | | | | | 16 | 3.9 | 21 | 0.001 | 0.72 |
| | | | | | 22 | 4.6 | 21 | <0.0001 | 0.67 |
| | | | | | 29 | 4.1 | 21 | <0.0001 | 0.7 |
| | | | | | 36 | 1.9 | 21 | 0.069 | 0.85 |
| 431.3313 (-) (lm) | day | 11.22 | 6,56 | <0.0001 | -3 | -1.7 | 56 | 0.1 | 1.1 |
| | treatment | 17.14 | 1,56 | <0.0001 | 1 | -0.4 | 56 | 0.69 | 1.02 |
| | day:treatment | 15.27 | 6,56 | <0.0001 | 8 | -0.5 | 56 | 0.61 | 1.03 |
| | | | | | 16 | 8.3 | 56 | <0.0001 | 0.52 |
| | | | | | 22 | 5.9 | 56 | <0.0001 | 0.64 |
| | | | | | 29 | -1.3 | 56 | 0.21 | 1.09 |
| | | | | | 36 | 0.6 | 56 | 0.54 | 0.96 |
| 459.3297 (+) (lm) | day | 17.15 | 6,56 | <0.0001 | -3 | -1.9 | 56 | 0.06 | 1.07 |
| | treatment | 318.89 | 1,56 | <0.0001 | 1 | 2.2 | 56 | 0.033 | 0.91 |
| | day:treatment | 41.99 | 6,56 | <0.0001 | 8 | 9.1 | 56 | <0.0001 | 0.69 |
| | | | | | 16 | 14.2 | 56 | <0.0001 | 0.58 |
| | | | | | 22 | 15 | 56 | <0.0001 | 0.59 |
| | | | | | 29 | 7.4 | 56 | <0.0001 | 0.77 |
| | | | | | 36 | 1.5 | 56 | 0.15 | 0.95 |
| 466.3661 (+) (lmm) | day | 147.7 | 6 | <0.0001 | -3 | -0.3 | 25 | 0.78 | 1.06 |
| | treatment | 23.01 | 1 | <0.0001 | 1 | -0.4 | 25 | 0.68 | 1.09 |
| | day:treatment | 127.66 | 6 | <0.0001 | 8 | -2.1 | 25 | 0.042 | 1.48 |
| | | | | | 16 | -8.5 | 25 | <0.0001 | 2.78 |
| | | | | | 22 | -7.9 | 25 | <0.0001 | 2.62 |
| | | | | | 29 | -3.7 | 25 | 0.001 | 1.82 |
| | | | | | 36 | -1.5 | 25 | 0.14 | 1.37 |
| 468.3813 (+) (lmm) | day | 147.74 | 6 | <0.0001 | -3 | -0.3 | 24 | 0.76 | 1.06 |
| | treatment | 21.77 | 1 | <0.0001 | 1 | -0.8 | 24 | 0.41 | 1.15 |
| | day:treatment | 102.18 | 6 | <0.0001 | 8 | -2.8 | 24 | 0.011 | 1.53 |
| | | | | | 16 | -7.9 | 24 | <0.0001 | 2.33 |
| | | | | | 22 | -7.2 | 24 | <0.0001 | 2.21 |
| | | | | | 29 | -3.4 | 24 | 0.002 | 1.64 |
| | | | | | 36 | -1.8 | 24 | 0.086 | 1.38 |
| 470.3982 (+) (lmm) | day | 137.27 | 6 | <0.0001 | -3 | -0.3 | 23 | 0.76 | 1.05 |
| | treatment | 19.89 | 1 | <0.0001 | 1 | -1.2 | 23 | 0.23 | 1.22 |
| | day:treatment | 65.64 | 6 | <0.0001 | 8 | -4.2 | 23 | <0.0001 | 1.73 |
| | | | | | 16 | -6.9 | 23 | <0.0001 | 2.02 |
| | | | | | 22 | -5.5 | 23 | <0.0001 | 1.8 |
| | | | | | 29 | -3.3 | 23 | 0.003 | 1.55 |
| | | | | | 36 | -2.1 | 23 | 0.051 | 1.41 |
| 474.4323 (+) (lm) | day | 6.56 | 6,56 | <0.0001 | -3 | -0.9 | 56 | 0.4 | 1.04 |
| | treatment | 0.48 | 1,56 | 0.5 | 1 | -2.4 | 56 | 0.022 | 1.12 |
| | day:treatment | 12.39 | 6,56 | <0.0001 | 8 | -3.5 | 56 | 0.001 | 1.19 |
| | | | | | 16 | 5.6 | 56 | <0.0001 | 0.72 |
| | | | | | 22 | 3.5 | 56 | 0.001 | 0.82 |
| | | | | | 29 | -3.4 | 56 | 0.001 | 1.19 |
| | | | | | 36 | -0.9 | 56 | 0.38 | 1.05 |
| 475.4348 (+) (lm) | day | 8.2 | 6,56 | <0.0001 | -3 | -1.1 | 56 | 0.26 | 1.07 |
| | treatment | 0.02 | 1,56 | 0.9 | 1 | -1.9 | 56 | 0.057 | 1.12 |
| | day:treatment | 11.82 | 6,56 | <0.0001 | 8 | -2.9 | 56 | 0.006 | 1.18 |
| | | | | | 16 | 5.9 | 56 | <0.0001 | 0.66 |
| | | | | | 22 | 3.8 | 56 | <0.0001 | 0.77 |
| | | | | | 29 | -2.9 | 56 | 0.006 | 1.19 |
| | | | | | 36 | -0.5 | 56 | 0.6 | 1.04 |
| 476.436 (+) (lm) | day | 9.88 | 6,56 | <0.0001 | -3 | -1.4 | 56 | 0.17 | 1.09 |
| | treatment | 1.37 | 1,56 | 0.26 | 1 | -1.1 | 56 | 0.27 | 1.07 |
| | day:treatment | 11.52 | 6,56 | <0.0001 | 8 | -2.3 | 56 | 0.024 | 1.16 |
| | | | | | 16 | 6.3 | 56 | <0.0001 | 0.61 |
| | | | | | 22 | 4 | 56 | <0.0001 | 0.74 |

Continued on next page

Table 7 – continued from previous page

| Pre-resistance pattern | | | | | | | | | |
|---|---|---|---|---|---|---|---|---|---|
| | lmm/lm | | | | Pairwise contrasts | | | | |
| **m/z peak** | *fixed effect* | $\chi^2/F$ | *df* | *adj. p* | *contrast* | *t-ratio* | *df* | *p* | *fold change* |
| | | | | | 29 | -2.4 | 56 | 0.019 | 1.17 |
| | | | | | 36 | 0 | 56 | 1 | 1 |
| 498.4334 (+) (lmm) | day | 12.19 | 6 | 0.063 | -3 | -1.7 | 48 | 0.099 | 1.12 |
| | treatment | 32.85 | 1 | <0.0001 | 1 | -4 | 48 | <0.0001 | 1.3 |
| | day:treatment | 49.96 | 6 | <0.0001 | 8 | -4.9 | 48 | <0.0001 | 1.36 |
| | | | | | 16 | 0.2 | 48 | 0.85 | 0.99 |
| | | | | | 22 | 0.3 | 48 | 0.74 | 0.98 |
| | | | | | 29 | -5.3 | 48 | <0.0001 | 1.39 |
| | | | | | 36 | -6 | 48 | <0.0001 | 1.49 |
| 499.4369 (+) (lmm) | day | 9.36 | 6 | 0.16 | -3 | -1.7 | 48 | 0.099 | 1.14 |
| | treatment | 25.66 | 1 | <0.0001 | 1 | -3.6 | 48 | 0.001 | 1.31 |
| | day:treatment | 47.52 | 6 | <0.0001 | 8 | -4.4 | 48 | <0.0001 | 1.37 |
| | | | | | 16 | 0.6 | 48 | 0.54 | 0.95 |
| | | | | | 22 | 0.7 | 48 | 0.5 | 0.95 |
| | | | | | 29 | -4.9 | 48 | <0.0001 | 1.41 |
| | | | | | 36 | -5.5 | 48 | <0.0001 | 1.52 |
| 500.4505 (+) (lmm) | day | 16.49 | 6 | 0.013 | -3 | -1.6 | 42 | 0.11 | 1.11 |
| | treatment | 13.01 | 1 | <0.0001 | 1 | -3.2 | 42 | 0.003 | 1.23 |
| | day:treatment | 58.05 | 6 | <0.0001 | 8 | -3.4 | 42 | 0.002 | 1.25 |
| | | | | | 16 | 2.7 | 42 | 0.01 | 0.82 |
| | | | | | 22 | 0 | 42 | 0.99 | 1 |
| | | | | | 29 | -4.7 | 42 | <0.0001 | 1.36 |
| | | | | | 36 | -4.7 | 42 | <0.0001 | 1.36 |
| 501.451 (+) (lmm) | day | 16.92 | 6 | 0.011 | -3 | -1.8 | 43 | 0.084 | 1.13 |
| | treatment | 9.36 | 1 | 0.003 | 1 | -2.7 | 43 | 0.009 | 1.21 |
| | day:treatment | 53.21 | 6 | <0.0001 | 8 | -2.7 | 43 | 0.01 | 1.21 |
| | | | | | 16 | 3 | 43 | 0.005 | 0.78 |
| | | | | | 22 | 0.3 | 43 | 0.77 | 0.98 |
| | | | | | 29 | -4.2 | 43 | <0.0001 | 1.35 |
| | | | | | 36 | -4.3 | 43 | <0.0001 | 1.36 |
| 518.4089 (-) (lm) | day | 9.88 | 6,56 | <0.0001 | -3 | -1.6 | 56 | 0.12 | 1.1 |
| | treatment | 9.81 | 1,56 | 0.003 | 1 | -0.9 | 56 | 0.4 | 1.05 |
| | day:treatment | 14.52 | 6,56 | <0.0001 | 8 | -1.2 | 56 | 0.24 | 1.08 |
| | | | | | 16 | 7.7 | 56 | <0.0001 | 0.54 |
| | | | | | 22 | 5.4 | 56 | <0.0001 | 0.66 |
| | | | | | 29 | -1.6 | 56 | 0.12 | 1.11 |
| | | | | | 36 | 0.3 | 56 | 0.76 | 0.98 |
| 519.4099 (-) (lm) | day | 12.78 | 6,56 | <0.0001 | -3 | -1.7 | 56 | 0.1 | 1.1 |
| | treatment | 19.86 | 1,56 | <0.0001 | 1 | -0.3 | 56 | 0.8 | 1.02 |
| | day:treatment | 15.12 | 6,56 | <0.0001 | 8 | -0.4 | 56 | 0.72 | 1.02 |
| | | | | | 16 | 8.4 | 56 | <0.0001 | 0.51 |
| | | | | | 22 | 6 | 56 | <0.0001 | 0.64 |
| | | | | | 29 | -1.1 | 56 | 0.27 | 1.08 |
| | | | | | 36 | 0.8 | 56 | 0.41 | 0.94 |
| 538.432 (+) (lmm) | day | 4.01 | 6 | 0.68 | -3 | -0.6 | 41 | 0.58 | 1.05 |
| | treatment | 17.65 | 1 | <0.0001 | 1 | -3.3 | 41 | 0.002 | 1.33 |
| | day:treatment | 44.68 | 6 | <0.0001 | 8 | -4.4 | 41 | <0.0001 | 1.44 |
| | | | | | 16 | 0 | 41 | 0.98 | 1 |
| | | | | | 22 | 0.3 | 41 | 0.75 | 0.97 |
| | | | | | 29 | -4.4 | 41 | <0.0001 | 1.45 |
| | | | | | 36 | -5.2 | 41 | <0.0001 | 1.58 |
| 540.3954 (-) (lmm) | day | 20.12 | 6 | 0.003 | -3 | -1.6 | 31 | 0.13 | 1.17 |
| | treatment | 15.37 | 1 | <0.0001 | 1 | -2.6 | 31 | 0.015 | 1.27 |
| | day:treatment | 27.59 | 6 | <0.0001 | 8 | -3.8 | 31 | 0.001 | 1.39 |
| | | | | | 16 | -0.9 | 31 | 0.36 | 1.08 |
| | | | | | 22 | -0.6 | 31 | 0.56 | 1.05 |
| | | | | | 29 | -4.4 | 31 | <0.0001 | 1.43 |
| | | | | | 36 | -4.8 | 31 | <0.0001 | 1.59 |
| 553.2922 (-) (lmm) | day | 42.44 | 6 | <0.0001 | -3 | -0.3 | 36 | 0.76 | 1.03 |
| | treatment | 15.32 | 1 | <0.0001 | 1 | -0.2 | 36 | 0.86 | 1.02 |
| | day:treatment | 61.82 | 6 | <0.0001 | 8 | 0 | 36 | 0.98 | 1 |
| | | | | | 16 | -6.8 | 36 | <0.0001 | 1.75 |
| | | | | | 22 | -5.4 | 36 | <0.0001 | 1.57 |
| | | | | | 29 | -2.8 | 36 | 0.009 | 1.32 |
| | | | | | 36 | -2 | 36 | 0.058 | 1.22 |
| 567.374 (+) (lm) | day | 20.2 | 6,56 | <0.0001 | -3 | -1.4 | 56 | 0.17 | 1.07 |
| | treatment | 1.86 | 1,56 | 0.19 | 1 | 3.7 | 56 | <0.0001 | 0.84 |
| | day:treatment | 7.92 | 6,56 | <0.0001 | 8 | 1.6 | 56 | 0.11 | 0.9 |
| | | | | | 16 | -3.9 | 56 | <0.0001 | 1.24 |
| | | | | | 22 | -3.9 | 56 | <0.0001 | 1.27 |
| | | | | | 29 | -0.5 | 56 | 0.64 | 1.03 |
| | | | | | 36 | 0.7 | 56 | 0.46 | 0.96 |
| 579.4701 (+) (lmm) | day | 34.39 | 6 | <0.0001 | -3 | -1.6 | 56 | 0.12 | 1.1 |
| | treatment | 1.03 | 1 | 0.32 | 1 | 3.6 | 56 | 0.001 | 0.8 |
| | day:treatment | 85.33 | 6 | <0.0001 | 8 | 4.3 | 56 | <0.0001 | 0.74 |
| | | | | | 16 | -4.2 | 56 | <0.0001 | 1.28 |
| | | | | | 22 | -5.8 | 56 | <0.0001 | 1.39 |

Continued on next page

Table 7 – continued from previous page

| Pre-resistance pattern | | | | | | | | | |
|---|---|---|---|---|---|---|---|---|---|
| | **lmm/lm** | | | | **Pairwise contrasts** | | | | |
| **m/z peak** | *fixed effect* | $\chi^2/F$ | *df* | *adj. p* | *contrast* | *t-ratio* | *df* | *p* | *fold change* |
| | | | | | 29 | 0 | 56 | 0.97 | 1 |
| | | | | | 36 | 0.8 | 56 | 0.42 | 0.95 |
| 584.4998 (+) (lmm) | day | 83.88 | 6 | <0.0001 | -3 | -1 | 39 | 0.33 | 1.04 |
| | treatment | 4.1 | 1 | 0.047 | 1 | 2.6 | 39 | 0.012 | 0.89 |
| | day:treatment | 25.05 | 6 | <0.0001 | 8 | 4.2 | 39 | <0.0001 | 0.85 |
| | | | | | 16 | 1.1 | 39 | 0.28 | 0.96 |
| | | | | | 22 | -0.2 | 39 | 0.85 | 1.01 |
| | | | | | 29 | 1.2 | 39 | 0.25 | 0.96 |
| | | | | | 36 | 0.6 | 39 | 0.52 | 0.97 |
| 597.4818 (+) (lm) | day | 6.91 | 6,56 | <0.0001 | -3 | -1.6 | 56 | 0.11 | 1.08 |
| | treatment | 0.75 | 1,56 | 0.4 | 1 | 4 | 56 | <0.0001 | 0.83 |
| | day:treatment | 12.95 | 6,56 | <0.0001 | 8 | 5.4 | 56 | <0.0001 | 0.75 |
| | | | | | 16 | -2.5 | 56 | 0.017 | 1.13 |
| | | | | | 22 | -4.7 | 56 | <0.0001 | 1.25 |
| | | | | | 29 | 0.6 | 56 | 0.56 | 0.97 |
| | | | | | 36 | 1.1 | 56 | 0.28 | 0.95 |
| 600.4962 (+) (lmm) | day | 74.72 | 6 | <0.0001 | -3 | -0.2 | 52 | 0.86 | 1.01 |
| | treatment | 40.28 | 1 | <0.0001 | 1 | 1.6 | 52 | 0.12 | 0.95 |
| | day:treatment | 99.49 | 6 | <0.0001 | 8 | 7.7 | 52 | <0.0001 | 0.78 |
| | | | | | 16 | 8.5 | 52 | <0.0001 | 0.75 |
| | | | | | 22 | 4.1 | 52 | <0.0001 | 0.88 |
| | | | | | 29 | 1.4 | 52 | 0.16 | 0.96 |
| | | | | | 36 | -1.5 | 52 | 0.13 | 1.04 |
| 601.4983 (+) (lmm) | day | 95.55 | 6 | <0.0001 | -3 | 0 | 52 | 0.98 | 1 |
| | treatment | 78.03 | 1 | <0.0001 | 1 | 1.7 | 52 | 0.1 | 0.93 |
| | day:treatment | 98.38 | 6 | <0.0001 | 8 | 7.5 | 52 | <0.0001 | 0.72 |
| | | | | | 16 | 10.5 | 52 | <0.0001 | 0.61 |
| | | | | | 22 | 5.8 | 52 | <0.0001 | 0.77 |
| | | | | | 29 | 3.9 | 52 | <0.0001 | 0.86 |
| | | | | | 36 | 1 | 52 | 0.3 | 0.96 |
| 602.5017 (+) (lmm) | day | 106.54 | 6 | <0.0001 | -3 | 0 | 52 | 0.97 | 1 |
| | treatment | 104.12 | 1 | <0.0001 | 1 | 2.7 | 52 | 0.008 | 0.9 |
| | day:treatment | 100.85 | 6 | <0.0001 | 8 | 8.4 | 52 | <0.0001 | 0.71 |
| | | | | | 16 | 10.9 | 52 | <0.0001 | 0.62 |
| | | | | | 22 | 6.4 | 52 | <0.0001 | 0.76 |
| | | | | | 29 | 4.7 | 52 | <0.0001 | 0.83 |
| | | | | | 36 | 1.8 | 52 | 0.082 | 0.94 |
| 691.4552 (-) (lm) | day | 4.81 | 6,56 | 0.001 | -3 | 0 | 56 | 1 | 1 |
| | treatment | 1.62 | 1,56 | 0.22 | 1 | 1.3 | 56 | 0.18 | 0.91 |
| | day:treatment | 4.39 | 6,56 | 0.001 | 8 | 4.4 | 56 | <0.0001 | 0.7 |
| | | | | | 16 | -1.3 | 56 | 0.19 | 1.1 |
| | | | | | 22 | -2.1 | 56 | 0.037 | 1.17 |
| | | | | | 29 | 0.2 | 56 | 0.81 | 0.98 |
| | | | | | 36 | 0.9 | 56 | 0.38 | 0.94 |
| 693.425 (+) (lmm) | day | 180.64 | 6 | <0.0001 | -3 | -0.3 | 47 | 0.8 | 1.01 |
| | treatment | 35.14 | 1 | <0.0001 | 1 | -4.3 | 47 | <0.0001 | 1.36 |
| | day:treatment | 24.89 | 6 | <0.0001 | 8 | -3.6 | 47 | 0.001 | 1.4 |
| | | | | | 16 | -2.7 | 47 | 0.009 | 1.29 |
| | | | | | 22 | -1.5 | 47 | 0.14 | 1.15 |
| | | | | | 29 | -4.4 | 47 | <0.0001 | 1.54 |
| | | | | | 36 | -5.6 | 47 | <0.0001 | 1.68 |
| 694.4306 (+) (lmm) | day | 213.2 | 6 | <0.0001 | -3 | -0.1 | 48 | 0.93 | 1 |
| | treatment | 32.45 | 1 | <0.0001 | 1 | -4 | 48 | <0.0001 | 1.25 |
| | day:treatment | 21.18 | 6 | 0.002 | 8 | -3.6 | 48 | 0.001 | 1.28 |
| | | | | | 16 | -3.2 | 48 | 0.002 | 1.25 |
| | | | | | 22 | -1.5 | 48 | 0.15 | 1.11 |
| | | | | | 29 | -4.1 | 48 | <0.0001 | 1.36 |
| | | | | | 36 | -5 | 48 | <0.0001 | 1.42 |
| 704.6229 (+) (lmm) | day | 121.64 | 6 | <0.0001 | -3 | 0.4 | 35 | 0.68 | 0.94 |
| | treatment | 8.17 | 1 | 0.005 | 1 | -0.7 | 35 | 0.49 | 1.09 |
| | day:treatment | 68.4 | 6 | <0.0001 | 8 | 0.9 | 35 | 0.39 | 0.91 |
| | | | | | 16 | -3.4 | 35 | 0.002 | 1.33 |
| | | | | | 22 | -7.2 | 35 | <0.0001 | 1.91 |
| | | | | | 29 | -2.2 | 35 | 0.036 | 1.22 |
| | | | | | 36 | -0.6 | 35 | 0.54 | 1.07 |
| 706.6441 (+) (lmm) | day | 181.3 | 6 | <0.0001 | -3 | 0.6 | 37 | 0.56 | 0.94 |
| | treatment | 0.77 | 1 | 0.39 | 1 | -3.1 | 37 | 0.004 | 1.3 |
| | day:treatment | 36.19 | 6 | <0.0001 | 8 | 0.4 | 37 | 0.73 | 0.98 |
| | | | | | 16 | -0.5 | 37 | 0.61 | 1.03 |
| | | | | | 22 | -3.8 | 37 | 0.001 | 1.28 |
| | | | | | 29 | 1.2 | 37 | 0.25 | 0.93 |
| | | | | | 36 | 1.4 | 37 | 0.18 | 0.91 |
| 730.6486 (+) (lmm) | day | 209.56 | 6 | <0.0001 | -3 | 0.6 | 39 | 0.56 | 0.94 |
| | treatment | 4.47 | 1 | 0.038 | 1 | -2.3 | 39 | 0.028 | 1.22 |
| | day:treatment | 30.1 | 6 | <0.0001 | 8 | -0.7 | 39 | 0.49 | 1.05 |
| | | | | | 16 | -2.2 | 39 | 0.035 | 1.14 |
| | | | | | 22 | -4.8 | 39 | <0.0001 | 1.35 |

Continued on next page

Table 7 – continued from previous page

| Pre-resistance pattern | | | | | | | | | |
|---|---|---|---|---|---|---|---|---|---|
| | **lmm/lm** | | | | **Pairwise contrasts** | | | | |
| **m/z peak** | *fixed effect* | $\chi^2/F$ | *df* | *adj. p* | *contrast* | *t-ratio* | *df* | *p* | *fold change* |
| | | | | | 29 | 0 | 39 | 0.96 | 1 |
| | | | | | 36 | 0.2 | 39 | 0.84 | 0.99 |
| 732.6665 (+) (lmm) | day | 161.87 | 6 | <0.0001 | -3 | 0.9 | 47 | 0.39 | 0.96 |
| | treatment | 0.2 | 1 | 0.66 | 1 | -5.2 | 47 | <0.0001 | 1.22 |
| | day:treatment | 51.5 | 6 | <0.0001 | 8 | -2.6 | 47 | 0.014 | 1.1 |
| | | | | | 16 | 1.6 | 47 | 0.12 | 0.94 |
| | | | | | 22 | 0.8 | 47 | 0.4 | 0.97 |
| | | | | | 29 | 0.8 | 47 | 0.42 | 0.97 |
| | | | | | 36 | 2 | 47 | 0.056 | 0.94 |
| 733.6681 (+) (lmm) | day | 129.03 | 6 | <0.0001 | -3 | 0.9 | 47 | 0.36 | 0.96 |
| | treatment | 0.08 | 1 | 0.79 | 1 | -4.8 | 47 | <0.0001 | 1.22 |
| | day:treatment | 51.68 | 6 | <0.0001 | 8 | -2.3 | 47 | 0.028 | 1.1 |
| | | | | | 16 | 2.3 | 47 | 0.029 | 0.92 |
| | | | | | 22 | 1.6 | 47 | 0.12 | 0.94 |
| | | | | | 29 | 1.2 | 47 | 0.24 | 0.96 |
| | | | | | 36 | 2.2 | 47 | 0.036 | 0.92 |
| 734.6806 (+) (lmm) | day | 206.96 | 6 | <0.0001 | -3 | 0.8 | 42 | 0.45 | 0.97 |
| | treatment | 3.64 | 1 | 0.062 | 1 | -5.4 | 42 | <0.0001 | 1.2 |
| | day:treatment | 142.41 | 6 | <0.0001 | 8 | -3.1 | 42 | 0.003 | 1.11 |
| | | | | | 16 | 4 | 42 | <0.0001 | 0.88 |
| | | | | | 22 | 7.2 | 42 | <0.0001 | 0.78 |
| | | | | | 29 | 2.7 | 42 | 0.009 | 0.92 |
| | | | | | 36 | 1.8 | 42 | 0.085 | 0.95 |
| 736.6934 (+) (lmm) | day | 147.75 | 6 | <0.0001 | -3 | 0.5 | 38 | 0.63 | 0.98 |
| | treatment | 8.48 | 1 | 0.004 | 1 | -3.5 | 38 | 0.001 | 1.15 |
| | day:treatment | 146.21 | 6 | <0.0001 | 8 | -2.5 | 38 | 0.018 | 1.11 |
| | | | | | 16 | 4.7 | 38 | <0.0001 | 0.82 |
| | | | | | 22 | 8.5 | 38 | <0.0001 | 0.69 |
| | | | | | 29 | 3.5 | 38 | 0.001 | 0.87 |
| | | | | | 36 | 1.5 | 38 | 0.14 | 0.95 |
| 737.6999 (+) (lmm) | day | 128.45 | 6 | <0.0001 | -3 | 0.3 | 39 | 0.8 | 0.99 |
| | treatment | 11.82 | 1 | 0.001 | 1 | -3 | 39 | 0.005 | 1.14 |
| | day:treatment | 145.85 | 6 | <0.0001 | 8 | -2.1 | 39 | 0.04 | 1.1 |
| | | | | | 16 | 5 | 39 | <0.0001 | 0.79 |
| | | | | | 22 | 9 | 39 | <0.0001 | 0.64 |
| | | | | | 29 | 3.8 | 39 | <0.0001 | 0.85 |
| | | | | | 36 | 1.8 | 39 | 0.086 | 0.93 |
| 738.7081 (+) (lmm) | day | 111.04 | 6 | <0.0001 | -3 | 0 | 43 | 0.96 | 1 |
| | treatment | 6.75 | 1 | 0.011 | 1 | -3.8 | 43 | 0.001 | 1.2 |
| | day:treatment | 120.41 | 6 | <0.0001 | 8 | -2.2 | 43 | 0.031 | 1.12 |
| | | | | | 16 | 4.7 | 43 | <0.0001 | 0.77 |
| | | | | | 22 | 7.7 | 43 | <0.0001 | 0.64 |
| | | | | | 29 | 2.8 | 43 | 0.008 | 0.87 |
| | | | | | 36 | 1.4 | 43 | 0.15 | 0.94 |
| 739.7131 (+) (lmm) | day | 100.04 | 6 | <0.0001 | -3 | -0.2 | 47 | 0.87 | 1.01 |
| | treatment | 9.39 | 1 | 0.003 | 1 | -3.4 | 47 | 0.001 | 1.2 |
| | day:treatment | 106.69 | 6 | <0.0001 | 8 | -1.7 | 47 | 0.093 | 1.11 |
| | | | | | 16 | 4.9 | 47 | <0.0001 | 0.73 |
| | | | | | 22 | 7.5 | 47 | <0.0001 | 0.6 |
| | | | | | 29 | 2.8 | 47 | 0.008 | 0.85 |
| | | | | | 36 | 1.9 | 47 | 0.069 | 0.91 |
| 758.6823 (+) (lmm) | day | 192.65 | 6 | <0.0001 | -3 | 0.7 | 41 | 0.48 | 0.96 |
| | treatment | 12.17 | 1 | 0.001 | 1 | -4.1 | 41 | <0.0001 | 1.2 |
| | day:treatment | 147.23 | 6 | <0.0001 | 8 | -1.9 | 41 | 0.064 | 1.08 |
| | | | | | 16 | 5.9 | 41 | <0.0001 | 0.77 |
| | | | | | 22 | 8.2 | 41 | <0.0001 | 0.69 |
| | | | | | 29 | 4 | 41 | <0.0001 | 0.85 |
| | | | | | 36 | 1.7 | 41 | 0.09 | 0.94 |
| 760.6995 (+) (lmm) | day | 171.65 | 6 | <0.0001 | -3 | 0.3 | 38 | 0.79 | 0.99 |
| | treatment | 6.41 | 1 | 0.013 | 1 | -3.9 | 38 | <0.0001 | 1.2 |
| | day:treatment | 140.02 | 6 | <0.0001 | 8 | -2.5 | 38 | 0.018 | 1.12 |
| | | | | | 16 | 4.9 | 38 | <0.0001 | 0.79 |
| | | | | | 22 | 7.7 | 38 | <0.0001 | 0.68 |
| | | | | | 29 | 3.4 | 38 | 0.002 | 0.86 |
| | | | | | 36 | 1.1 | 38 | 0.3 | 0.96 |
| 761.7011 (+) (lmm) | day | 147.87 | 6 | <0.0001 | -3 | 0.3 | 38 | 0.73 | 0.98 |
| | treatment | 8.16 | 1 | 0.005 | 1 | -3.5 | 38 | 0.001 | 1.2 |
| | day:treatment | 141.2 | 6 | <0.0001 | 8 | -2.2 | 38 | 0.034 | 1.12 |
| | | | | | 16 | 5.3 | 38 | <0.0001 | 0.76 |
| | | | | | 22 | 8 | 38 | <0.0001 | 0.64 |
| | | | | | 29 | 3.6 | 38 | 0.001 | 0.84 |
| | | | | | 36 | 1 | 38 | 0.32 | 0.96 |
| 762.7091 (+) (lmm) | day | 133.83 | 6 | <0.0001 | -3 | 0 | 37 | 1 | 1 |
| | treatment | 8.16 | 1 | 0.005 | 1 | -2.8 | 37 | 0.009 | 1.16 |
| | day:treatment | 124.88 | 6 | <0.0001 | 8 | -2.1 | 37 | 0.047 | 1.11 |
| | | | | | 16 | 4.9 | 37 | <0.0001 | 0.77 |
| | | | | | 22 | 7.8 | 37 | <0.0001 | 0.65 |

Continued on next page

Table 7 – continued from previous page

| Pre-resistance pattern | | | | | | | | | |
|---|---|---|---|---|---|---|---|---|---|
| | **lmm/lm** | | | | **Pairwise contrasts** | | | | |
| **m/z peak** | *fixed effect* | $\chi^2/F$ | *df* | *adj. p* | *contrast* | *t-ratio* | *df* | *p* | *fold change* |
| | | | | | 29 | 3.7 | 37 | 0.001 | 0.83 |
| | | | | | 36 | 0.9 | 37 | 0.36 | 0.96 |
| 763.7131 (+) (lmm) | day | 109.94 | 6 | <0.0001 | -3 | 0 | 38 | 0.99 | 1 |
| | treatment | 12.9 | 1 | <0.0001 | 1 | -1.9 | 38 | 0.062 | 1.11 |
| | day:treatment | 119.8 | 6 | <0.0001 | 8 | -1.4 | 38 | 0.17 | 1.08 |
| | | | | | 16 | 5.3 | 38 | <0.0001 | 0.73 |
| | | | | | 22 | 8.3 | 38 | <0.0001 | 0.6 |
| | | | | | 29 | 4.1 | 38 | <0.0001 | 0.8 |
| | | | | | 36 | 1.1 | 38 | 0.28 | 0.95 |
| 766.6665 (+) (lmm) | day | 27.43 | 6 | <0.0001 | -3 | 1.9 | 51 | 0.063 | 0.88 |
| | treatment | 2.59 | 1 | 0.12 | 1 | 0.6 | 51 | 0.54 | 0.96 |
| | day:treatment | 11.26 | 6 | 0.088 | 8 | 2.4 | 51 | 0.02 | 0.82 |
| | | | | | 16 | -0.6 | 51 | 0.54 | 1.04 |
| | | | | | 22 | -1.1 | 51 | 0.29 | 1.07 |
| | | | | | 29 | 1.5 | 51 | 0.14 | 0.89 |
| | | | | | 36 | 1 | 51 | 0.34 | 0.94 |
| 767.5632 (+) (lmm) | day | 387.47 | 6 | <0.0001 | -3 | -1.4 | 41 | 0.17 | 1.28 |
| | treatment | 1.79 | 1 | 0.19 | 1 | 0.4 | 41 | 0.69 | 0.95 |
| | day:treatment | 10.91 | 6 | 0.099 | 8 | -0.3 | 41 | 0.73 | 1.03 |
| | | | | | 16 | -2.6 | 41 | 0.012 | 1.22 |
| | | | | | 22 | -1.7 | 41 | 0.09 | 1.13 |
| | | | | | 29 | -0.5 | 41 | 0.65 | 1.03 |
| | | | | | 36 | 0.6 | 41 | 0.58 | 0.95 |
| 768.5688 (+) (lmm) | day | 378.08 | 6 | <0.0001 | -3 | -1.1 | 37 | 0.28 | 1.18 |
| | treatment | 1.2 | 1 | 0.28 | 1 | 0.5 | 37 | 0.63 | 0.94 |
| | day:treatment | 13.72 | 6 | 0.037 | 8 | 0.1 | 37 | 0.93 | 0.99 |
| | | | | | 16 | -2.7 | 37 | 0.011 | 1.21 |
| | | | | | 22 | -2 | 37 | 0.054 | 1.15 |
| | | | | | 29 | -0.3 | 37 | 0.8 | 1.02 |
| | | | | | 36 | 0.6 | 37 | 0.53 | 0.94 |
| 769.5764 (+) (lmm) | day | 487.1 | 6 | <0.0001 | -3 | -1.8 | 51 | 0.081 | 1.23 |
| | treatment | 0.05 | 1 | 0.83 | 1 | 1.2 | 51 | 0.23 | 0.89 |
| | day:treatment | 12.54 | 6 | 0.056 | 8 | -0.2 | 51 | 0.85 | 1.01 |
| | | | | | 16 | -2.2 | 51 | 0.036 | 1.13 |
| | | | | | 22 | 0.6 | 51 | 0.57 | 0.97 |
| | | | | | 29 | 1.1 | 51 | 0.27 | 0.95 |
| | | | | | 36 | 0.4 | 51 | 0.66 | 0.97 |
| 776.6966 (+) (lmm) | day | 49.03 | 6 | <0.0001 | -3 | 0.8 | 22 | 0.41 | 0.96 |
| | treatment | 0.75 | 1 | 0.4 | 1 | -2.3 | 22 | 0.03 | 1.12 |
| | day:treatment | 151.99 | 6 | <0.0001 | 8 | -3.4 | 22 | 0.002 | 1.18 |
| | | | | | 16 | 4.8 | 22 | <0.0001 | 0.79 |
| | | | | | 22 | 5.5 | 22 | <0.0001 | 0.75 |
| | | | | | 29 | 1.3 | 22 | 0.21 | 0.94 |
| | | | | | 36 | -2 | 22 | 0.054 | 1.1 |
| 777.6983 (+) (lmm) | day | 44.25 | 6 | <0.0001 | -3 | 0.9 | 21 | 0.39 | 0.95 |
| | treatment | 1.33 | 1 | 0.26 | 1 | -2.1 | 21 | 0.043 | 1.11 |
| | day:treatment | 167.74 | 6 | <0.0001 | 8 | -3.3 | 21 | 0.004 | 1.18 |
| | | | | | 16 | 5.4 | 21 | <0.0001 | 0.75 |
| | | | | | 22 | 5.9 | 21 | <0.0001 | 0.72 |
| | | | | | 29 | 1.5 | 21 | 0.14 | 0.92 |
| | | | | | 36 | -2.1 | 21 | 0.051 | 1.11 |
| 784.5426 (+) (lmm) | day | 595.3 | 6 | <0.0001 | -3 | -1.7 | 41 | 0.092 | 1.26 |
| | treatment | 14.5 | 1 | <0.0001 | 1 | 0.5 | 41 | 0.6 | 0.94 |
| | day:treatment | 35.14 | 6 | <0.0001 | 8 | -4.1 | 41 | <0.0001 | 1.3 |
| | | | | | 16 | -5.1 | 41 | <0.0001 | 1.35 |
| | | | | | 22 | -3.7 | 41 | 0.001 | 1.24 |
| | | | | | 29 | -1.7 | 41 | 0.09 | 1.1 |
| | | | | | 36 | -0.1 | 41 | 0.95 | 1 |
| 788.6551 (+) (lmm) | day | 34.5 | 6 | <0.0001 | -3 | 1.1 | 50 | 0.28 | 0.92 |
| | treatment | 2.22 | 1 | 0.14 | 1 | 1.1 | 50 | 0.29 | 0.92 |
| | day:treatment | 24.93 | 6 | <0.0001 | 8 | 3.6 | 50 | 0.001 | 0.74 |
| | | | | | 16 | -0.8 | 50 | 0.46 | 1.05 |
| | | | | | 22 | -2.4 | 50 | 0.021 | 1.17 |
| | | | | | 29 | 1.2 | 50 | 0.24 | 0.92 |
| | | | | | 36 | 1.6 | 50 | 0.11 | 0.89 |
| 793.5764 (-) (lmm) | day | 113.32 | 6 | <0.0001 | -3 | 1.9 | 56 | 0.062 | 0.92 |
| | treatment | 5.96 | 1 | 0.017 | 1 | -3.6 | 56 | 0.001 | 1.15 |
| | day:treatment | 54.75 | 6 | <0.0001 | 8 | -2.6 | 56 | 0.013 | 1.1 |
| | | | | | 16 | 1.1 | 56 | 0.27 | 0.96 |
| | | | | | 22 | 4.2 | 56 | <0.0001 | 0.86 |
| | | | | | 29 | 3.9 | 56 | <0.0001 | 0.87 |
| | | | | | 36 | 1.8 | 56 | 0.071 | 0.94 |
| 794.5791 (-) (lmm) | day | 98.57 | 6 | <0.0001 | -3 | 1.9 | 56 | 0.06 | 0.92 |
| | treatment | 10.85 | 1 | 0.001 | 1 | -3.5 | 56 | 0.001 | 1.14 |
| | day:treatment | 62.83 | 6 | <0.0001 | 8 | -2.5 | 56 | 0.014 | 1.1 |
| | | | | | 16 | 1.5 | 56 | 0.13 | 0.94 |
| | | | | | 22 | 5 | 56 | <0.0001 | 0.83 |

Continued on next page

Table 7 – continued from previous page

| Pre-resistance pattern | | | | | | | | | |
|---|---|---|---|---|---|---|---|---|---|
| | lmm/lm | | | | Pairwise contrasts | | | | |
| **m/z peak** | *fixed effect* | $\chi^2/F$ | *df* | *adj. p* | *contrast* | *t-ratio* | *df* | *p* | *fold change* |
| | | | | | 29 | 4.4 | 56 | <0.0001 | 0.85 |
| | | | | | 36 | 2.3 | 56 | 0.027 | 0.92 |
| 798.6798 (+) (lmm) | day | 46.75 | 6 | <0.0001 | -3 | 1.9 | 30 | 0.068 | 0.9 |
| | treatment | 3.45 | 1 | 0.069 | 1 | -1.9 | 30 | 0.062 | 1.1 |
| | day:treatment | 57.95 | 6 | <0.0001 | 8 | -1.9 | 30 | 0.064 | 1.1 |
| | | | | | 16 | 4 | 30 | <0.0001 | 0.83 |
| | | | | | 22 | 3.7 | 30 | 0.001 | 0.83 |
| | | | | | 29 | 2.5 | 30 | 0.02 | 0.89 |
| | | | | | 36 | 0.7 | 30 | 0.48 | 0.97 |
| 800.6925 (+) (lmm) | day | 40.2 | 6 | <0.0001 | -3 | 1.5 | 27 | 0.14 | 0.91 |
| | treatment | 10.36 | 1 | 0.002 | 1 | -2.5 | 27 | 0.02 | 1.14 |
| | day:treatment | 149.75 | 6 | <0.0001 | 8 | -0.3 | 27 | 0.78 | 1.02 |
| | | | | | 16 | 7.8 | 27 | <0.0001 | 0.63 |
| | | | | | 22 | 6.8 | 27 | <0.0001 | 0.67 |
| | | | | | 29 | 2.6 | 27 | 0.014 | 0.86 |
| | | | | | 36 | 0 | 27 | 0.97 | 1 |
| 807.5381 (-) (lmm) | day | 161.53 | 6 | <0.0001 | -3 | -1.1 | 44 | 0.26 | 1.15 |
| | treatment | 5.78 | 1 | 0.018 | 1 | 1.8 | 44 | 0.074 | 0.81 |
| | day:treatment | 87.66 | 6 | <0.0001 | 8 | 1.2 | 44 | 0.22 | 0.88 |
| | | | | | 16 | -5.5 | 44 | <0.0001 | 1.55 |
| | | | | | 22 | -6.7 | 44 | <0.0001 | 1.62 |
| | | | | | 29 | -0.2 | 44 | 0.87 | 1.01 |
| | | | | | 36 | 0.8 | 44 | 0.46 | 0.92 |
| 821.5621 (-) (lmm) | day | 354.4 | 6 | <0.0001 | -3 | -1.1 | 46 | 0.29 | 1.06 |
| | treatment | 21.24 | 1 | <0.0001 | 1 | 0.5 | 46 | 0.63 | 0.98 |
| | day:treatment | 146.24 | 6 | <0.0001 | 8 | 1 | 46 | 0.32 | 0.96 |
| | | | | | 16 | -7.2 | 46 | <0.0001 | 1.32 |
| | | | | | 22 | -10.4 | 46 | <0.0001 | 1.45 |
| | | | | | 29 | -1.4 | 46 | 0.16 | 1.06 |
| | | | | | 36 | 0.7 | 46 | 0.46 | 0.97 |
| 824.5551 (-) (lmm) | day | 145.32 | 6 | <0.0001 | -3 | -1.5 | 50 | 0.14 | 1.08 |
| | treatment | 1.44 | 1 | 0.24 | 1 | 4.6 | 50 | <0.0001 | 0.78 |
| | day:treatment | 107.72 | 6 | <0.0001 | 8 | 4.7 | 50 | <0.0001 | 0.79 |
| | | | | | 16 | -3.7 | 50 | 0.001 | 1.17 |
| | | | | | 22 | -4.6 | 50 | <0.0001 | 1.21 |
| | | | | | 29 | 1.3 | 50 | 0.2 | 0.94 |
| | | | | | 36 | 3.5 | 50 | 0.001 | 0.83 |
| 831.5377 (+) (lmm) | day | 88.37 | 6 | <0.0001 | -3 | -1.1 | 49 | 0.3 | 1.23 |
| | treatment | 11.08 | 1 | 0.001 | 1 | 1.5 | 49 | 0.14 | 0.76 |
| | day:treatment | 94.67 | 6 | <0.0001 | 8 | 1.9 | 49 | 0.066 | 0.73 |
| | | | | | 16 | -6.6 | 49 | <0.0001 | 2.21 |
| | | | | | 22 | -7.1 | 49 | <0.0001 | 2.16 |
| | | | | | 29 | -1.1 | 49 | 0.3 | 1.16 |
| | | | | | 36 | 0.1 | 49 | 0.93 | 0.98 |
| 847.5316 (+) (lmm) | day | 75.11 | 6 | <0.0001 | -3 | -0.8 | 51 | 0.44 | 1.23 |
| | treatment | 11.5 | 1 | 0.001 | 1 | 1.2 | 51 | 0.26 | 0.76 |
| | day:treatment | 79.43 | 6 | <0.0001 | 8 | 1.7 | 51 | 0.091 | 0.67 |
| | | | | | 16 | -6.3 | 51 | <0.0001 | 2.56 |
| | | | | | 22 | -6.6 | 51 | <0.0001 | 2.41 |
| | | | | | 29 | -1 | 51 | 0.33 | 1.2 |
| | | | | | 36 | 0 | 51 | 0.98 | 1 |
| 848.5355 (+) (lmm) | day | 62.2 | 6 | <0.0001 | -3 | -0.5 | 52 | 0.63 | 1.12 |
| | treatment | 9.88 | 1 | 0.002 | 1 | 1 | 52 | 0.3 | 0.79 |
| | day:treatment | 68.88 | 6 | <0.0001 | 8 | 1.4 | 52 | 0.16 | 0.73 |
| | | | | | 16 | -5.8 | 52 | <0.0001 | 2.37 |
| | | | | | 22 | -6.3 | 52 | <0.0001 | 2.31 |
| | | | | | 29 | -0.8 | 52 | 0.46 | 1.15 |
| | | | | | 36 | 0.1 | 52 | 0.93 | 0.98 |
| 849.5403 (+) (lmm) | day | 48.66 | 6 | <0.0001 | -3 | -0.5 | 54 | 0.59 | 1.13 |
| | treatment | 6.77 | 1 | 0.011 | 1 | 1.2 | 54 | 0.23 | 0.78 |
| | day:treatment | 56.83 | 6 | <0.0001 | 8 | 1.7 | 54 | 0.088 | 0.7 |
| | | | | | 16 | -5.6 | 54 | <0.0001 | 2.21 |
| | | | | | 22 | -5.1 | 54 | <0.0001 | 1.95 |
| | | | | | 29 | -0.2 | 54 | 0.84 | 1.04 |
| | | | | | 36 | 0.3 | 54 | 0.76 | 0.94 |
| 857.5345 (-) (lmm) | day | 453.03 | 6 | <0.0001 | -3 | -1.3 | 40 | 0.21 | 1.12 |
| | treatment | 0.81 | 1 | 0.38 | 1 | 1.8 | 40 | 0.081 | 0.87 |
| | day:treatment | 31.71 | 6 | <0.0001 | 8 | -1 | 40 | 0.31 | 1.06 |
| | | | | | 16 | -3 | 40 | 0.005 | 1.17 |
| | | | | | 22 | -2.8 | 40 | 0.008 | 1.14 |
| | | | | | 29 | 1.2 | 40 | 0.24 | 0.94 |
| | | | | | 36 | 1.2 | 40 | 0.22 | 0.92 |
| 905.5121 (-) (lmm) | day | 62.63 | 6 | <0.0001 | -3 | -0.8 | 42 | 0.42 | 1.09 |
| | treatment | 17.21 | 1 | <0.0001 | 1 | 0.4 | 42 | 0.72 | 0.97 |
| | day:treatment | 80.05 | 6 | <0.0001 | 8 | -0.2 | 42 | 0.83 | 1.02 |
| | | | | | 16 | -6.8 | 42 | <0.0001 | 1.73 |
| | | | | | 22 | -7.2 | 42 | <0.0001 | 1.75 |

Continued on next page

Table 7 – continued from previous page

| Pre-resistance pattern | | | | | | | | | |
|---|---|---|---|---|---|---|---|---|---|
| | lmm/lm | | | | Pairwise contrasts | | | | |
| **m/z peak** | *fixed effect* | $\chi^2/F$ | *df* | *adj. p* | *contrast* | *t-ratio* | *df* | *p* | *fold change* |
| | | | | | 29 | -1.5 | 42 | 0.13 | 1.15 |
| | | | | | 36 | -0.7 | 42 | 0.48 | 1.08 |
| 969.6719 (+) (lm) | day | 12.58 | 6,56 | <0.0001 | -3 | 1.8 | 56 | 0.083 | 0.88 |
| | treatment | 13.83 | 1,56 | 0.001 | 1 | 2.7 | 56 | 0.008 | 0.81 |
| | day:treatment | 1.95 | 6,56 | 0.095 | 8 | 3.1 | 56 | 0.003 | 0.72 |
| | | | | | 16 | -0.5 | 56 | 0.64 | 1.05 |
| | | | | | 22 | -0.4 | 56 | 0.72 | 1.03 |
| | | | | | 29 | 1.2 | 56 | 0.23 | 0.88 |
| | | | | | 36 | 1.8 | 56 | 0.074 | 0.85 |
| 985.6313 (-) (lmm) | day | 37.25 | 6 | <0.0001 | -3 | -0.8 | 45 | 0.44 | 1.03 |
| | treatment | 12.65 | 1 | <0.0001 | 1 | 2.4 | 45 | 0.019 | 0.9 |
| | day:treatment | 46.5 | 6 | <0.0001 | 8 | 6.4 | 45 | <0.0001 | 0.74 |
| | | | | | 16 | 1 | 45 | 0.31 | 0.96 |
| | | | | | 22 | -0.7 | 45 | 0.49 | 1.03 |
| | | | | | 29 | 2.4 | 45 | 0.021 | 0.89 |
| | | | | | 36 | 3.1 | 45 | 0.003 | 0.87 |
| 987.6336 (-) (lmm) | day | 29.62 | 6 | <0.0001 | -3 | 0.2 | 45 | 0.88 | 0.99 |
| | treatment | 16.47 | 1 | <0.0001 | 1 | 2.4 | 45 | 0.023 | 0.89 |
| | day:treatment | 25.87 | 6 | <0.0001 | 8 | 5.2 | 45 | <0.0001 | 0.76 |
| | | | | | 16 | 1.6 | 45 | 0.11 | 0.92 |
| | | | | | 22 | 0.2 | 45 | 0.88 | 0.99 |
| | | | | | 29 | 2.8 | 45 | 0.008 | 0.86 |
| | | | | | 36 | 3.7 | 45 | 0.001 | 0.83 |

Table 8: Analysis of Deviance Table (Type II Wald $\chi^2$-tests) for linear mixed effects model for all m/z peaks of interest with post-resistance pattern from exploratory analysis with percentage ion count (arc sine transformed) of the m/z peak as response. Where the lmm yielded a singular fit, an lm has been fit instead. P-values have been adjusted using the Benjamini-Hochberg procedure. Pairwise contrasts by day, as estimated by the `emmeans` package for R.

| Post-resistance pattern | | | | | | | | | |
|---|---|---|---|---|---|---|---|---|---|
| | lmm/lm | | | | Pairwise contrasts | | | | |
| **m/z peak** | *fixed effect* | $\chi^2/F$ | *df* | *adj. p* | *contrast* | *t-ratio* | *df* | *p* | *fold change* |
| 146.978 (-) (lmm) | day | 59.83 | 6 | <0.0001 | -3 | -0.3 | 53 | 0.74 | 1.06 |
| | treatment | 8.4 | 1 | 0.005 | 1 | 0.2 | 53 | 0.83 | 0.97 |
| | day:treatment | 11.05 | 6 | 0.094 | 8 | 0.8 | 53 | 0.41 | 0.89 |
| | | | | | 16 | 2.4 | 53 | 0.02 | 0.69 |
| | | | | | 22 | 3.4 | 53 | 0.001 | 0.62 |
| | | | | | 29 | 1.8 | 53 | 0.083 | 0.82 |
| | | | | | 36 | 1.2 | 53 | 0.23 | 0.88 |
| 158.9903 (-) (lmm) | day | 83.14 | 6 | <0.0001 | -3 | -1.6 | 54 | 0.13 | 1.06 |
| | treatment | 63.55 | 1 | <0.0001 | 1 | 3.7 | 54 | 0.001 | 0.87 |
| | day:treatment | 78.23 | 6 | <0.0001 | 8 | 7.3 | 54 | <0.0001 | 0.73 |
| | | | | | 16 | 8.6 | 54 | <0.0001 | 0.67 |
| | | | | | 22 | 4 | 54 | <0.0001 | 0.83 |
| | | | | | 29 | 1.7 | 54 | 0.097 | 0.93 |
| | | | | | 36 | 2.1 | 54 | 0.042 | 0.92 |
| 215.0478 (-) (lmm) | day | 76.5 | 6 | <0.0001 | -3 | 0.7 | 45 | 0.51 | 0.95 |
| | treatment | 1.03 | 1 | 0.32 | 1 | -3.5 | 45 | 0.001 | 1.25 |
| | day:treatment | 96.44 | 6 | <0.0001 | 8 | -3.6 | 45 | 0.001 | 1.23 |
| | | | | | 16 | 5.2 | 45 | <0.0001 | 0.69 |
| | | | | | 22 | 5.2 | 45 | <0.0001 | 0.68 |
| | | | | | 29 | 0.4 | 45 | 0.68 | 0.97 |
| | | | | | 36 | -0.4 | 45 | 0.71 | 1.02 |
| 223.1496 (-) (lm) | day | 12.04 | 6,56 | <0.0001 | -3 | -0.8 | 56 | 0.45 | 1.07 |
| | treatment | 22.89 | 1,56 | <0.0001 | 1 | 0.9 | 56 | 0.39 | 0.93 |
| | day:treatment | 6.83 | 6,56 | <0.0001 | 8 | 1.2 | 56 | 0.24 | 0.85 |
| | | | | | 16 | -5.5 | 56 | <0.0001 | 1.73 |
| | | | | | 22 | -4.9 | 56 | <0.0001 | 1.72 |
| | | | | | 29 | -1.9 | 56 | 0.067 | 1.27 |
| | | | | | 36 | -1.6 | 56 | 0.11 | 1.19 |
| 228.0716 (-) (lmm) | day | 175.59 | 6 | <0.0001 | -3 | 0.9 | 46 | 0.38 | 0.96 |
| | treatment | 0.04 | 1 | 0.84 | 1 | -2.7 | 46 | 0.01 | 1.11 |
| | day:treatment | 34.3 | 6 | <0.0001 | 8 | -1.5 | 46 | 0.15 | 1.05 |
| | | | | | 16 | 1.4 | 46 | 0.17 | 0.95 |
| | | | | | 22 | 3.6 | 46 | 0.001 | 0.88 |
| | | | | | 29 | 0.7 | 46 | 0.48 | 0.98 |
| | | | | | 36 | -1.7 | 46 | 0.1 | 1.06 |
| 229.0732 (-) (lmm) | day | 142.24 | 6 | <0.0001 | -3 | 0.7 | 47 | 0.48 | 0.96 |
| | treatment | 2.32 | 1 | 0.14 | 1 | -1.9 | 47 | 0.065 | 1.09 |
| | day:treatment | 32.27 | 6 | <0.0001 | 8 | -0.6 | 47 | 0.52 | 1.03 |
| | | | | | 16 | 2.1 | 47 | 0.039 | 0.91 |

Continued on next page

Table 8 – continued from previous page

| Post-resistance pattern | | | | | | | | | |
|---|---|---|---|---|---|---|---|---|---|
| | lmm/lm | | | | Pairwise contrasts | | | | |
| **m/z peak** | *fixed effect* | $\chi^2/F$ | *df* | *adj. p* | *contrast* | *t-ratio* | *df* | *p* | *fold change* |
| | | | | | 22 | 4.4 | 47 | <0.0001 | 0.83 |
| | | | | | 29 | 1.7 | 47 | 0.092 | 0.93 |
| | | | | | 36 | -0.5 | 47 | 0.6 | 1.02 |
| 230.0696 (-) (lmm) | day | 139.46 | 6 | <0.0001 | -3 | 0.8 | 47 | 0.42 | 0.96 |
| | treatment | 2.77 | 1 | 0.1 | 1 | -2 | 47 | 0.049 | 1.1 |
| | day:treatment | 34.44 | 6 | <0.0001 | 8 | -0.4 | 47 | 0.7 | 1.02 |
| | | | | | 16 | 2.3 | 47 | 0.024 | 0.91 |
| | | | | | 22 | 4.5 | 47 | <0.0001 | 0.82 |
| | | | | | 29 | 1.7 | 47 | 0.097 | 0.93 |
| | | | | | 36 | -0.6 | 47 | 0.53 | 1.02 |
| 242.073 (-) (lmm) | day | 93.73 | 6 | <0.0001 | -3 | 0.7 | 42 | 0.52 | 0.94 |
| | treatment | 0.06 | 1 | 0.81 | 1 | -4.2 | 42 | <0.0001 | 1.4 |
| | day:treatment | 85.73 | 6 | <0.0001 | 8 | -4 | 42 | <0.0001 | 1.33 |
| | | | | | 16 | 3.1 | 42 | 0.003 | 0.77 |
| | | | | | 22 | 4.4 | 42 | <0.0001 | 0.68 |
| | | | | | 29 | 0.5 | 42 | 0.64 | 0.97 |
| | | | | | 36 | -1.6 | 42 | 0.12 | 1.11 |
| 261.09 (-) (lmm) | day | 124.86 | 6 | <0.0001 | -3 | 0.7 | 52 | 0.46 | 0.96 |
| | treatment | 27.73 | 1 | <0.0001 | 1 | 7.4 | 52 | <0.0001 | 0.66 |
| | day:treatment | 39.28 | 6 | <0.0001 | 8 | 1.6 | 52 | 0.11 | 0.9 |
| | | | | | 16 | 3.4 | 52 | 0.001 | 0.78 |
| | | | | | 22 | 3.2 | 52 | 0.002 | 0.8 |
| | | | | | 29 | 1.2 | 52 | 0.24 | 0.92 |
| | | | | | 36 | 0.6 | 52 | 0.58 | 0.97 |
| 269.2311 (-) (lmm) | day | 365.62 | 6 | <0.0001 | -3 | 0 | 56 | 0.98 | 1 |
| | treatment | 11.54 | 1 | 0.001 | 1 | 3.2 | 56 | 0.002 | 0.83 |
| | day:treatment | 15.8 | 6 | 0.017 | 8 | 3.4 | 56 | 0.001 | 0.79 |
| | | | | | 16 | 1.5 | 56 | 0.15 | 0.9 |
| | | | | | 22 | 1.8 | 56 | 0.072 | 0.86 |
| | | | | | 29 | -0.3 | 56 | 0.78 | 1.02 |
| | | | | | 36 | -0.5 | 56 | 0.64 | 1.03 |
| 286.1728 (+) (lmm) | day | 156.51 | 6 | <0.0001 | -3 | 0.3 | 51 | 0.74 | 0.98 |
| | treatment | 229.86 | 1 | <0.0001 | 1 | 2.3 | 51 | 0.025 | 0.86 |
| | day:treatment | 292.45 | 6 | <0.0001 | 8 | 11 | 51 | <0.0001 | 0.53 |
| | | | | | 16 | 15.1 | 51 | <0.0001 | 0.42 |
| | | | | | 22 | 15.6 | 51 | <0.0001 | 0.46 |
| | | | | | 29 | 7.6 | 51 | <0.0001 | 0.7 |
| | | | | | 36 | 1 | 51 | 0.31 | 0.94 |
| 297.263 (-) (lmm) | day | 366.94 | 6 | <0.0001 | -3 | -0.7 | 54 | 0.46 | 1.03 |
| | treatment | 7.44 | 1 | 0.007 | 1 | 3.2 | 54 | 0.003 | 0.86 |
| | day:treatment | 19.01 | 6 | 0.005 | 8 | 3.7 | 54 | <0.0001 | 0.81 |
| | | | | | 16 | 1 | 54 | 0.33 | 0.95 |
| | | | | | 22 | 1.7 | 54 | 0.091 | 0.9 |
| | | | | | 29 | 0.2 | 54 | 0.85 | 0.99 |
| | | | | | 36 | -0.3 | 54 | 0.8 | 1.01 |
| 301.1833 (-) (lmm) | day | 149 | 6 | <0.0001 | -3 | 0 | 50 | 0.99 | 1 |
| | treatment | 234.39 | 1 | <0.0001 | 1 | 3.6 | 50 | 0.001 | 0.71 |
| | day:treatment | 283.97 | 6 | <0.0001 | 8 | 11.9 | 50 | <0.0001 | 0.38 |
| | | | | | 16 | 15.1 | 50 | <0.0001 | 0.29 |
| | | | | | 22 | 15.3 | 50 | <0.0001 | 0.35 |
| | | | | | 29 | 7.6 | 50 | <0.0001 | 0.63 |
| | | | | | 36 | 1.5 | 50 | 0.13 | 0.88 |
| 302.1732 (-) (lmm) | day | 72.54 | 6 | <0.0001 | -3 | 1.4 | 50 | 0.17 | 0.92 |
| | treatment | 131.79 | 1 | <0.0001 | 1 | 1.9 | 50 | 0.066 | 0.86 |
| | day:treatment | 177.2 | 6 | <0.0001 | 8 | 5.9 | 50 | <0.0001 | 0.69 |
| | | | | | 16 | 12.6 | 50 | <0.0001 | 0.4 |
| | | | | | 22 | 12.8 | 50 | <0.0001 | 0.43 |
| | | | | | 29 | 5.8 | 50 | <0.0001 | 0.7 |
| | | | | | 36 | 1.1 | 50 | 0.29 | 0.92 |
| 303.2167 (+) (lmm) | day | 131.61 | 6 | <0.0001 | -3 | 0 | 54 | 0.97 | 1 |
| | treatment | 170.75 | 1 | <0.0001 | 1 | 1.1 | 54 | 0.3 | 0.92 |
| | day:treatment | 224.97 | 6 | <0.0001 | 8 | 9.5 | 54 | <0.0001 | 0.52 |
| | | | | | 16 | 12.8 | 54 | <0.0001 | 0.42 |
| | | | | | 22 | 13.2 | 54 | <0.0001 | 0.46 |
| | | | | | 29 | 5.6 | 54 | <0.0001 | 0.75 |
| | | | | | 36 | 0 | 54 | 0.98 | 1 |
| 304.1946 (+) (lmm) | day | 55.3 | 6 | <0.0001 | -3 | 1.4 | 45 | 0.17 | 0.91 |
| | treatment | 71.31 | 1 | <0.0001 | 1 | 0.7 | 45 | 0.49 | 0.94 |
| | day:treatment | 176.08 | 6 | <0.0001 | 8 | 3.5 | 45 | 0.001 | 0.79 |
| | | | | | 16 | 11.5 | 45 | <0.0001 | 0.39 |
| | | | | | 22 | 11.5 | 45 | <0.0001 | 0.44 |
| | | | | | 29 | 4.3 | 45 | <0.0001 | 0.76 |
| | | | | | 36 | 0.1 | 45 | 0.89 | 0.99 |
| 311.1896 (-) (lmm) | day | 411.77 | 6 | <0.0001 | -3 | 0.7 | 47 | 0.49 | 0.97 |
| | treatment | 0.03 | 1 | 0.87 | 1 | -7.8 | 47 | <0.0001 | 1.28 |
| | day:treatment | 107.55 | 6 | <0.0001 | 8 | -0.8 | 47 | 0.41 | 1.02 |
| | | | | | 16 | 3.1 | 47 | 0.004 | 0.92 |

Continued on next page

Table 8 – continued from previous page

| Post-resistance pattern | | | | | | | | | |
|---|---|---|---|---|---|---|---|---|---|
| | lmm/lm | | | | Pairwise contrasts | | | | |
| **m/z peak** | *fixed effect* | $\chi^2/F$ | *df* | *adj. p* | *contrast* | *t-ratio* | *df* | *p* | *fold change* |
| | | | | | 22 | 3.9 | 47 | <0.0001 | 0.89 |
| | | | | | 29 | 1 | 47 | 0.34 | 0.97 |
| | | | | | 36 | 0.7 | 47 | 0.51 | 0.98 |
| 311.2436 (-) (lm) | day | 30.54 | 6,56 | <0.0001 | -3 | 0.8 | 56 | 0.41 | 0.95 |
| | treatment | 4.84 | 1,56 | 0.036 | 1 | 0.7 | 56 | 0.46 | 0.95 |
| | day:treatment | 3.55 | 6,56 | 0.006 | 8 | 1.7 | 56 | 0.097 | 0.86 |
| | | | | | 16 | -3.2 | 56 | 0.002 | 1.3 |
| | | | | | 22 | -2.4 | 56 | 0.022 | 1.24 |
| | | | | | 29 | -1.7 | 56 | 0.1 | 1.19 |
| | | | | | 36 | -1.8 | 56 | 0.071 | 1.18 |
| 346.0795 (-) (lmm) | day | 93.52 | 6 | <0.0001 | -3 | -1.8 | 21 | 0.085 | 1.22 |
| | treatment | 3.16 | 1 | 0.082 | 1 | -0.8 | 21 | 0.4 | 1.08 |
| | day:treatment | 113.62 | 6 | <0.0001 | 8 | -2.4 | 21 | 0.025 | 1.19 |
| | | | | | 16 | 3.9 | 21 | 0.001 | 0.72 |
| | | | | | 22 | 4.6 | 21 | <0.0001 | 0.67 |
| | | | | | 29 | 4.1 | 21 | <0.0001 | 0.7 |
| | | | | | 36 | 1.9 | 21 | 0.069 | 0.85 |
| 431.3313 (-) (lm) | day | 11.22 | 6,56 | <0.0001 | -3 | -1.7 | 56 | 0.1 | 1.1 |
| | treatment | 17.14 | 1,56 | <0.0001 | 1 | -0.4 | 56 | 0.69 | 1.02 |
| | day:treatment | 15.27 | 6,56 | <0.0001 | 8 | -0.5 | 56 | 0.61 | 1.03 |
| | | | | | 16 | 8.3 | 56 | <0.0001 | 0.52 |
| | | | | | 22 | 5.9 | 56 | <0.0001 | 0.64 |
| | | | | | 29 | -1.3 | 56 | 0.21 | 1.09 |
| | | | | | 36 | 0.6 | 56 | 0.54 | 0.96 |
| 459.3297 (+) (lm) | day | 17.15 | 6,56 | <0.0001 | -3 | -1.9 | 56 | 0.06 | 1.07 |
| | treatment | 318.89 | 1,56 | <0.0001 | 1 | 2.2 | 56 | 0.033 | 0.91 |
| | day:treatment | 41.99 | 6,56 | <0.0001 | 8 | 9.1 | 56 | <0.0001 | 0.69 |
| | | | | | 16 | 14.2 | 56 | <0.0001 | 0.58 |
| | | | | | 22 | 15 | 56 | <0.0001 | 0.59 |
| | | | | | 29 | 7.4 | 56 | <0.0001 | 0.77 |
| | | | | | 36 | 1.5 | 56 | 0.15 | 0.95 |
| 466.3661 (+) (lmm) | day | 147.7 | 6 | <0.0001 | -3 | -0.3 | 25 | 0.78 | 1.06 |
| | treatment | 23.01 | 1 | <0.0001 | 1 | -0.4 | 25 | 0.68 | 1.09 |
| | day:treatment | 127.66 | 6 | <0.0001 | 8 | -2.1 | 25 | 0.042 | 1.48 |
| | | | | | 16 | -8.5 | 25 | <0.0001 | 2.78 |
| | | | | | 22 | -7.9 | 25 | <0.0001 | 2.62 |
| | | | | | 29 | -3.7 | 25 | 0.001 | 1.82 |
| | | | | | 36 | -1.5 | 25 | 0.14 | 1.37 |
| 468.3813 (+) (lmm) | day | 147.74 | 6 | <0.0001 | -3 | -0.3 | 24 | 0.76 | 1.06 |
| | treatment | 21.77 | 1 | <0.0001 | 1 | -0.8 | 24 | 0.41 | 1.15 |
| | day:treatment | 102.18 | 6 | <0.0001 | 8 | -2.8 | 24 | 0.011 | 1.53 |
| | | | | | 16 | -7.9 | 24 | <0.0001 | 2.33 |
| | | | | | 22 | -7.2 | 24 | <0.0001 | 2.21 |
| | | | | | 29 | -3.4 | 24 | 0.002 | 1.64 |
| | | | | | 36 | -1.8 | 24 | 0.086 | 1.38 |
| 470.3982 (+) (lmm) | day | 137.27 | 6 | <0.0001 | -3 | -0.3 | 23 | 0.76 | 1.05 |
| | treatment | 19.89 | 1 | <0.0001 | 1 | -1.2 | 23 | 0.23 | 1.22 |
| | day:treatment | 65.64 | 6 | <0.0001 | 8 | -4.2 | 23 | <0.0001 | 1.73 |
| | | | | | 16 | -6.9 | 23 | <0.0001 | 2.02 |
| | | | | | 22 | -5.5 | 23 | <0.0001 | 1.8 |
| | | | | | 29 | -3.3 | 23 | 0.003 | 1.55 |
| | | | | | 36 | -2.1 | 23 | 0.051 | 1.41 |
| 474.4323 (+) (lm) | day | 6.56 | 6,56 | <0.0001 | -3 | -0.9 | 56 | 0.4 | 1.04 |
| | treatment | 0.48 | 1,56 | 0.5 | 1 | -2.4 | 56 | 0.022 | 1.12 |
| | day:treatment | 12.39 | 6,56 | <0.0001 | 8 | -3.5 | 56 | 0.001 | 1.19 |
| | | | | | 16 | 5.6 | 56 | <0.0001 | 0.72 |
| | | | | | 22 | 3.5 | 56 | 0.001 | 0.82 |
| | | | | | 29 | -3.4 | 56 | 0.001 | 1.19 |
| | | | | | 36 | -0.9 | 56 | 0.38 | 1.05 |
| 475.4348 (+) (lm) | day | 8.2 | 6,56 | <0.0001 | -3 | -1.1 | 56 | 0.26 | 1.07 |
| | treatment | 0.02 | 1,56 | 0.9 | 1 | -1.9 | 56 | 0.057 | 1.12 |
| | day:treatment | 11.82 | 6,56 | <0.0001 | 8 | -2.9 | 56 | 0.006 | 1.18 |
| | | | | | 16 | 5.9 | 56 | <0.0001 | 0.66 |
| | | | | | 22 | 3.8 | 56 | <0.0001 | 0.77 |
| | | | | | 29 | -2.9 | 56 | 0.006 | 1.19 |
| | | | | | 36 | -0.5 | 56 | 0.6 | 1.04 |
| 476.436 (+) (lm) | day | 9.88 | 6,56 | <0.0001 | -3 | -1.4 | 56 | 0.17 | 1.09 |
| | treatment | 1.37 | 1,56 | 0.26 | 1 | -1.1 | 56 | 0.27 | 1.07 |
| | day:treatment | 11.52 | 6,56 | <0.0001 | 8 | -2.3 | 56 | 0.024 | 1.16 |
| | | | | | 16 | 6.3 | 56 | <0.0001 | 0.61 |
| | | | | | 22 | 4 | 56 | <0.0001 | 0.74 |
| | | | | | 29 | -2.4 | 56 | 0.019 | 1.17 |
| | | | | | 36 | 0 | 56 | 1 | 1 |
| 498.4334 (+) (lmm) | day | 12.19 | 6 | 0.063 | -3 | -1.7 | 48 | 0.099 | 1.12 |
| | treatment | 32.85 | 1 | <0.0001 | 1 | -4 | 48 | <0.0001 | 1.3 |
| | day:treatment | 49.96 | 6 | <0.0001 | 8 | -4.9 | 48 | <0.0001 | 1.36 |
| | | | | | 16 | 0.2 | 48 | 0.85 | 0.99 |

Continued on next page

Table 8 – continued from previous page

| Post-resistance pattern | | | | | | | | | |
|---|---|---|---|---|---|---|---|---|---|
| | lmm/lm | | | | Pairwise contrasts | | | | |
| **m/z peak** | *fixed effect* | $\chi^2/F$ | *df* | *adj. p* | *contrast* | *t-ratio* | *df* | *p* | *fold change* |
| | | | | | 22 | 0.3 | 48 | 0.74 | 0.98 |
| | | | | | 29 | -5.3 | 48 | <0.0001 | 1.39 |
| | | | | | 36 | -6 | 48 | <0.0001 | 1.49 |
| 499.4369 (+) (lmm) | day | 9.36 | 6 | 0.16 | -3 | -1.7 | 48 | 0.099 | 1.14 |
| | treatment | 25.66 | 1 | <0.0001 | 1 | -3.6 | 48 | 0.001 | 1.31 |
| | day:treatment | 47.52 | 6 | <0.0001 | 8 | -4.4 | 48 | <0.0001 | 1.37 |
| | | | | | 16 | 0.6 | 48 | 0.54 | 0.95 |
| | | | | | 22 | 0.7 | 48 | 0.5 | 0.95 |
| | | | | | 29 | -4.9 | 48 | <0.0001 | 1.41 |
| | | | | | 36 | -5.5 | 48 | <0.0001 | 1.52 |
| 500.4505 (+) (lmm) | day | 16.49 | 6 | 0.013 | -3 | -1.6 | 42 | 0.11 | 1.11 |
| | treatment | 13.01 | 1 | <0.0001 | 1 | -3.2 | 42 | 0.003 | 1.23 |
| | day:treatment | 58.05 | 6 | <0.0001 | 8 | -3.4 | 42 | 0.002 | 1.25 |
| | | | | | 16 | 2.7 | 42 | 0.01 | 0.82 |
| | | | | | 22 | 0 | 42 | 0.99 | 1 |
| | | | | | 29 | -4.7 | 42 | <0.0001 | 1.36 |
| | | | | | 36 | -4.7 | 42 | <0.0001 | 1.36 |
| 501.451 (+) (lmm) | day | 16.92 | 6 | 0.011 | -3 | -1.8 | 43 | 0.084 | 1.13 |
| | treatment | 9.36 | 1 | 0.003 | 1 | -2.7 | 43 | 0.009 | 1.21 |
| | day:treatment | 53.21 | 6 | <0.0001 | 8 | -2.7 | 43 | 0.01 | 1.21 |
| | | | | | 16 | 3 | 43 | 0.005 | 0.78 |
| | | | | | 22 | 0.3 | 43 | 0.77 | 0.98 |
| | | | | | 29 | -4.2 | 43 | <0.0001 | 1.35 |
| | | | | | 36 | -4.3 | 43 | <0.0001 | 1.36 |
| 518.4089 (-) (lm) | day | 9.88 | 6,56 | <0.0001 | -3 | -1.6 | 56 | 0.12 | 1.1 |
| | treatment | 9.81 | 1,56 | 0.003 | 1 | -0.9 | 56 | 0.4 | 1.05 |
| | day:treatment | 14.52 | 6,56 | <0.0001 | 8 | -1.2 | 56 | 0.24 | 1.08 |
| | | | | | 16 | 7.7 | 56 | <0.0001 | 0.54 |
| | | | | | 22 | 5.4 | 56 | <0.0001 | 0.66 |
| | | | | | 29 | -1.6 | 56 | 0.12 | 1.11 |
| | | | | | 36 | 0.3 | 56 | 0.76 | 0.98 |
| 519.4099 (-) (lm) | day | 12.78 | 6,56 | <0.0001 | -3 | -1.7 | 56 | 0.1 | 1.1 |
| | treatment | 19.86 | 1,56 | <0.0001 | 1 | -0.3 | 56 | 0.8 | 1.02 |
| | day:treatment | 15.12 | 6,56 | <0.0001 | 8 | -0.4 | 56 | 0.72 | 1.02 |
| | | | | | 16 | 8.4 | 56 | <0.0001 | 0.51 |
| | | | | | 22 | 6 | 56 | <0.0001 | 0.64 |
| | | | | | 29 | -1.1 | 56 | 0.27 | 1.08 |
| | | | | | 36 | 0.8 | 56 | 0.41 | 0.94 |
| 538.432 (+) (lmm) | day | 4.01 | 6 | 0.68 | -3 | -0.6 | 41 | 0.58 | 1.05 |
| | treatment | 17.65 | 1 | <0.0001 | 1 | -3.3 | 41 | 0.002 | 1.33 |
| | day:treatment | 44.68 | 6 | <0.0001 | 8 | -4.4 | 41 | <0.0001 | 1.44 |
| | | | | | 16 | 0 | 41 | 0.98 | 1 |
| | | | | | 22 | 0.3 | 41 | 0.75 | 0.97 |
| | | | | | 29 | -4.4 | 41 | <0.0001 | 1.45 |
| | | | | | 36 | -5.2 | 41 | <0.0001 | 1.58 |
| 540.3954 (-) (lmm) | day | 20.12 | 6 | 0.003 | -3 | -1.6 | 31 | 0.13 | 1.17 |
| | treatment | 15.37 | 1 | <0.0001 | 1 | -2.6 | 31 | 0.015 | 1.27 |
| | day:treatment | 27.59 | 6 | <0.0001 | 8 | -3.8 | 31 | 0.001 | 1.39 |
| | | | | | 16 | -0.9 | 31 | 0.36 | 1.08 |
| | | | | | 22 | -0.6 | 31 | 0.56 | 1.05 |
| | | | | | 29 | -4.4 | 31 | <0.0001 | 1.43 |
| | | | | | 36 | -4.8 | 31 | <0.0001 | 1.59 |
| 553.2922 (-) (lmm) | day | 42.44 | 6 | <0.0001 | -3 | -0.3 | 36 | 0.76 | 1.03 |
| | treatment | 15.32 | 1 | <0.0001 | 1 | -0.2 | 36 | 0.86 | 1.02 |
| | day:treatment | 61.82 | 6 | <0.0001 | 8 | 0 | 36 | 0.98 | 1 |
| | | | | | 16 | -6.8 | 36 | <0.0001 | 1.75 |
| | | | | | 22 | -5.4 | 36 | <0.0001 | 1.57 |
| | | | | | 29 | -2.8 | 36 | 0.009 | 1.32 |
| | | | | | 36 | -2 | 36 | 0.058 | 1.22 |
| 567.374 (+) (lm) | day | 20.2 | 6,56 | <0.0001 | -3 | -1.4 | 56 | 0.17 | 1.07 |
| | treatment | 1.86 | 1,56 | 0.19 | 1 | 3.7 | 56 | <0.0001 | 0.84 |
| | day:treatment | 7.92 | 6,56 | <0.0001 | 8 | 1.6 | 56 | 0.11 | 0.9 |
| | | | | | 16 | -3.9 | 56 | <0.0001 | 1.24 |
| | | | | | 22 | -3.9 | 56 | <0.0001 | 1.27 |
| | | | | | 29 | -0.5 | 56 | 0.64 | 1.03 |
| | | | | | 36 | 0.7 | 56 | 0.46 | 0.96 |
| 579.4701 (+) (lmm) | day | 34.39 | 6 | <0.0001 | -3 | -1.6 | 56 | 0.12 | 1.1 |
| | treatment | 1.03 | 1 | 0.32 | 1 | 3.6 | 56 | 0.001 | 0.8 |
| | day:treatment | 85.33 | 6 | <0.0001 | 8 | 4.3 | 56 | <0.0001 | 0.74 |
| | | | | | 16 | -4.2 | 56 | <0.0001 | 1.28 |
| | | | | | 22 | -5.8 | 56 | <0.0001 | 1.39 |
| | | | | | 29 | 0 | 56 | 0.97 | 1 |
| | | | | | 36 | 0.8 | 56 | 0.42 | 0.95 |
| 584.4998 (+) (lmm) | day | 83.88 | 6 | <0.0001 | -3 | -1 | 39 | 0.33 | 1.04 |
| | treatment | 4.1 | 1 | 0.047 | 1 | 2.6 | 39 | 0.012 | 0.89 |
| | day:treatment | 25.05 | 6 | <0.0001 | 8 | 4.2 | 39 | <0.0001 | 0.85 |
| | | | | | 16 | 1.1 | 39 | 0.28 | 0.96 |

Continued on next page

Table 8 – continued from previous page

| Post-resistance pattern | | | | | | | | | |
|---|---|---|---|---|---|---|---|---|---|
| | lmm/lm | | | | Pairwise contrasts | | | | |
| **m/z peak** | *fixed effect* | $\chi^2/F$ | *df* | *adj. p* | *contrast* | *t-ratio* | *df* | *p* | *fold change* |
| | | | | | 22 | -0.2 | 39 | 0.85 | 1.01 |
| | | | | | 29 | 1.2 | 39 | 0.25 | 0.96 |
| | | | | | 36 | 0.6 | 39 | 0.52 | 0.97 |
| 597.4818 (+) (lm) | day | 6.91 | 6,56 | <0.0001 | -3 | -1.6 | 56 | 0.11 | 1.08 |
| | treatment | 0.75 | 1,56 | 0.4 | 1 | 4 | 56 | <0.0001 | 0.83 |
| | day:treatment | 12.95 | 6,56 | <0.0001 | 8 | 5.4 | 56 | <0.0001 | 0.75 |
| | | | | | 16 | -2.5 | 56 | 0.017 | 1.13 |
| | | | | | 22 | -4.7 | 56 | <0.0001 | 1.25 |
| | | | | | 29 | 0.6 | 56 | 0.56 | 0.97 |
| | | | | | 36 | 1.1 | 56 | 0.28 | 0.95 |
| 600.4962 (+) (lmm) | day | 74.72 | 6 | <0.0001 | -3 | -0.2 | 52 | 0.86 | 1.01 |
| | treatment | 40.28 | 1 | <0.0001 | 1 | 1.6 | 52 | 0.12 | 0.95 |
| | day:treatment | 99.49 | 6 | <0.0001 | 8 | 7.7 | 52 | <0.0001 | 0.78 |
| | | | | | 16 | 8.5 | 52 | <0.0001 | 0.75 |
| | | | | | 22 | 4.1 | 52 | <0.0001 | 0.88 |
| | | | | | 29 | 1.4 | 52 | 0.16 | 0.96 |
| | | | | | 36 | -1.5 | 52 | 0.13 | 1.04 |
| 601.4983 (+) (lmm) | day | 95.55 | 6 | <0.0001 | -3 | 0 | 52 | 0.98 | 1 |
| | treatment | 78.03 | 1 | <0.0001 | 1 | 1.7 | 52 | 0.1 | 0.93 |
| | day:treatment | 98.38 | 6 | <0.0001 | 8 | 7.5 | 52 | <0.0001 | 0.72 |
| | | | | | 16 | 10.5 | 52 | <0.0001 | 0.61 |
| | | | | | 22 | 5.8 | 52 | <0.0001 | 0.77 |
| | | | | | 29 | 3.9 | 52 | <0.0001 | 0.86 |
| | | | | | 36 | 1 | 52 | 0.3 | 0.96 |
| 602.5017 (+) (lmm) | day | 106.54 | 6 | <0.0001 | -3 | 0 | 52 | 0.97 | 1 |
| | treatment | 104.12 | 1 | <0.0001 | 1 | 2.7 | 52 | 0.008 | 0.9 |
| | day:treatment | 100.85 | 6 | <0.0001 | 8 | 8.4 | 52 | <0.0001 | 0.71 |
| | | | | | 16 | 10.9 | 52 | <0.0001 | 0.62 |
| | | | | | 22 | 6.4 | 52 | <0.0001 | 0.76 |
| | | | | | 29 | 4.7 | 52 | <0.0001 | 0.83 |
| | | | | | 36 | 1.8 | 52 | 0.082 | 0.94 |
| 691.4552 (-) (lm) | day | 4.81 | 6,56 | 0.001 | -3 | 0 | 56 | 1 | 1 |
| | treatment | 1.62 | 1,56 | 0.22 | 1 | 1.3 | 56 | 0.18 | 0.91 |
| | day:treatment | 4.39 | 6,56 | 0.001 | 8 | 4.4 | 56 | <0.0001 | 0.7 |
| | | | | | 16 | -1.3 | 56 | 0.19 | 1.1 |
| | | | | | 22 | -2.1 | 56 | 0.037 | 1.17 |
| | | | | | 29 | 0.2 | 56 | 0.81 | 0.98 |
| | | | | | 36 | 0.9 | 56 | 0.38 | 0.94 |
| 693.425 (+) (lmm) | day | 180.64 | 6 | <0.0001 | -3 | -0.3 | 47 | 0.8 | 1.01 |
| | treatment | 35.14 | 1 | <0.0001 | 1 | -4.3 | 47 | <0.0001 | 1.36 |
| | day:treatment | 24.89 | 6 | <0.0001 | 8 | -3.6 | 47 | 0.001 | 1.4 |
| | | | | | 16 | -2.7 | 47 | 0.009 | 1.29 |
| | | | | | 22 | -1.5 | 47 | 0.14 | 1.15 |
| | | | | | 29 | -4.4 | 47 | <0.0001 | 1.54 |
| | | | | | 36 | -5.6 | 47 | <0.0001 | 1.68 |
| 694.4306 (+) (lmm) | day | 213.2 | 6 | <0.0001 | -3 | -0.1 | 48 | 0.93 | 1 |
| | treatment | 32.45 | 1 | <0.0001 | 1 | -4 | 48 | <0.0001 | 1.25 |
| | day:treatment | 21.18 | 6 | 0.002 | 8 | -3.6 | 48 | 0.001 | 1.28 |
| | | | | | 16 | -3.2 | 48 | 0.002 | 1.25 |
| | | | | | 22 | -1.5 | 48 | 0.15 | 1.11 |
| | | | | | 29 | -4.1 | 48 | <0.0001 | 1.36 |
| | | | | | 36 | -5 | 48 | <0.0001 | 1.42 |
| 704.6229 (+) (lmm) | day | 121.64 | 6 | <0.0001 | -3 | 0.4 | 35 | 0.68 | 0.94 |
| | treatment | 8.17 | 1 | 0.005 | 1 | -0.7 | 35 | 0.49 | 1.09 |
| | day:treatment | 68.4 | 6 | <0.0001 | 8 | 0.9 | 35 | 0.39 | 0.91 |
| | | | | | 16 | -3.4 | 35 | 0.002 | 1.33 |
| | | | | | 22 | -7.2 | 35 | <0.0001 | 1.91 |
| | | | | | 29 | -2.2 | 35 | 0.036 | 1.22 |
| | | | | | 36 | -0.6 | 35 | 0.54 | 1.07 |
| 706.6441 (+) (lmm) | day | 181.3 | 6 | <0.0001 | -3 | 0.6 | 37 | 0.56 | 0.94 |
| | treatment | 0.77 | 1 | 0.39 | 1 | -3.1 | 37 | 0.004 | 1.3 |
| | day:treatment | 36.19 | 6 | <0.0001 | 8 | 0.4 | 37 | 0.73 | 0.98 |
| | | | | | 16 | -0.5 | 37 | 0.61 | 1.03 |
| | | | | | 22 | -3.8 | 37 | 0.001 | 1.28 |
| | | | | | 29 | 1.2 | 37 | 0.25 | 0.93 |
| | | | | | 36 | 1.4 | 37 | 0.18 | 0.91 |
| 730.6486 (+) (lmm) | day | 209.56 | 6 | <0.0001 | -3 | 0.6 | 39 | 0.56 | 0.94 |
| | treatment | 4.47 | 1 | 0.038 | 1 | -2.3 | 39 | 0.028 | 1.22 |
| | day:treatment | 30.1 | 6 | <0.0001 | 8 | -0.7 | 39 | 0.49 | 1.05 |
| | | | | | 16 | -2.2 | 39 | 0.035 | 1.14 |
| | | | | | 22 | -4.8 | 39 | <0.0001 | 1.35 |
| | | | | | 29 | 0 | 39 | 0.96 | 1 |
| | | | | | 36 | 0.2 | 39 | 0.84 | 0.99 |
| 732.6665 (+) (lmm) | day | 161.87 | 6 | <0.0001 | -3 | 0.9 | 47 | 0.39 | 0.96 |
| | treatment | 0.2 | 1 | 0.66 | 1 | -5.2 | 47 | <0.0001 | 1.22 |
| | day:treatment | 51.5 | 6 | <0.0001 | 8 | -2.6 | 47 | 0.014 | 1.1 |
| | | | | | 16 | 1.6 | 47 | 0.12 | 0.94 |

Continued on next page

Table 8 – continued from previous page

| Post-resistance pattern | | | | | | | | | |
|---|---|---|---|---|---|---|---|---|---|
| | lmm/lm | | | | Pairwise contrasts | | | | |
| **m/z peak** | *fixed effect* | $\chi^2/F$ | *df* | *adj. p* | *contrast* | *t-ratio* | *df* | *p* | *fold change* |
| | | | | | 22 | 0.8 | 47 | 0.4 | 0.97 |
| | | | | | 29 | 0.8 | 47 | 0.42 | 0.97 |
| | | | | | 36 | 2 | 47 | 0.056 | 0.94 |
| 733.6681 (+) (lmm) | day | 129.03 | 6 | <0.0001 | -3 | 0.9 | 47 | 0.36 | 0.96 |
| | treatment | 0.08 | 1 | 0.79 | 1 | -4.8 | 47 | <0.0001 | 1.22 |
| | day:treatment | 51.68 | 6 | <0.0001 | 8 | -2.3 | 47 | 0.028 | 1.1 |
| | | | | | 16 | 2.3 | 47 | 0.029 | 0.92 |
| | | | | | 22 | 1.6 | 47 | 0.12 | 0.94 |
| | | | | | 29 | 1.2 | 47 | 0.24 | 0.96 |
| | | | | | 36 | 2.2 | 47 | 0.036 | 0.92 |
| 734.6806 (+) (lmm) | day | 206.96 | 6 | <0.0001 | -3 | 0.8 | 42 | 0.45 | 0.97 |
| | treatment | 3.64 | 1 | 0.062 | 1 | -5.4 | 42 | <0.0001 | 1.2 |
| | day:treatment | 142.41 | 6 | <0.0001 | 8 | -3.1 | 42 | 0.003 | 1.11 |
| | | | | | 16 | 4 | 42 | <0.0001 | 0.88 |
| | | | | | 22 | 7.2 | 42 | <0.0001 | 0.78 |
| | | | | | 29 | 2.7 | 42 | 0.009 | 0.92 |
| | | | | | 36 | 1.8 | 42 | 0.085 | 0.95 |
| 736.6934 (+) (lmm) | day | 147.75 | 6 | <0.0001 | -3 | 0.5 | 38 | 0.63 | 0.98 |
| | treatment | 8.48 | 1 | 0.004 | 1 | -3.5 | 38 | 0.001 | 1.15 |
| | day:treatment | 146.21 | 6 | <0.0001 | 8 | -2.5 | 38 | 0.018 | 1.11 |
| | | | | | 16 | 4.7 | 38 | <0.0001 | 0.82 |
| | | | | | 22 | 8.5 | 38 | <0.0001 | 0.69 |
| | | | | | 29 | 3.5 | 38 | 0.001 | 0.87 |
| | | | | | 36 | 1.5 | 38 | 0.14 | 0.95 |
| 737.6999 (+) (lmm) | day | 128.45 | 6 | <0.0001 | -3 | 0.3 | 39 | 0.8 | 0.99 |
| | treatment | 11.82 | 1 | 0.001 | 1 | -3 | 39 | 0.005 | 1.14 |
| | day:treatment | 145.85 | 6 | <0.0001 | 8 | -2.1 | 39 | 0.04 | 1.1 |
| | | | | | 16 | 5 | 39 | <0.0001 | 0.79 |
| | | | | | 22 | 9 | 39 | <0.0001 | 0.64 |
| | | | | | 29 | 3.8 | 39 | <0.0001 | 0.85 |
| | | | | | 36 | 1.8 | 39 | 0.086 | 0.93 |
| 738.7081 (+) (lmm) | day | 111.04 | 6 | <0.0001 | -3 | 0 | 43 | 0.96 | 1 |
| | treatment | 6.75 | 1 | 0.011 | 1 | -3.8 | 43 | 0.001 | 1.2 |
| | day:treatment | 120.41 | 6 | <0.0001 | 8 | -2.2 | 43 | 0.031 | 1.12 |
| | | | | | 16 | 4.7 | 43 | <0.0001 | 0.77 |
| | | | | | 22 | 7.7 | 43 | <0.0001 | 0.64 |
| | | | | | 29 | 2.8 | 43 | 0.008 | 0.87 |
| | | | | | 36 | 1.4 | 43 | 0.15 | 0.94 |
| 739.7131 (+) (lmm) | day | 100.04 | 6 | <0.0001 | -3 | -0.2 | 47 | 0.87 | 1.01 |
| | treatment | 9.39 | 1 | 0.003 | 1 | -3.4 | 47 | 0.001 | 1.2 |
| | day:treatment | 106.69 | 6 | <0.0001 | 8 | -1.7 | 47 | 0.093 | 1.11 |
| | | | | | 16 | 4.9 | 47 | <0.0001 | 0.73 |
| | | | | | 22 | 7.5 | 47 | <0.0001 | 0.6 |
| | | | | | 29 | 2.8 | 47 | 0.008 | 0.85 |
| | | | | | 36 | 1.9 | 47 | 0.069 | 0.91 |
| 758.6823 (+) (lmm) | day | 192.65 | 6 | <0.0001 | -3 | 0.7 | 41 | 0.48 | 0.96 |
| | treatment | 12.17 | 1 | 0.001 | 1 | -4.1 | 41 | <0.0001 | 1.2 |
| | day:treatment | 147.23 | 6 | <0.0001 | 8 | -1.9 | 41 | 0.064 | 1.08 |
| | | | | | 16 | 5.9 | 41 | <0.0001 | 0.77 |
| | | | | | 22 | 8.2 | 41 | <0.0001 | 0.69 |
| | | | | | 29 | 4 | 41 | <0.0001 | 0.85 |
| | | | | | 36 | 1.7 | 41 | 0.09 | 0.94 |
| 760.6995 (+) (lmm) | day | 171.65 | 6 | <0.0001 | -3 | 0.3 | 38 | 0.79 | 0.99 |
| | treatment | 6.41 | 1 | 0.013 | 1 | -3.9 | 38 | <0.0001 | 1.2 |
| | day:treatment | 140.02 | 6 | <0.0001 | 8 | -2.5 | 38 | 0.018 | 1.12 |
| | | | | | 16 | 4.9 | 38 | <0.0001 | 0.79 |
| | | | | | 22 | 7.7 | 38 | <0.0001 | 0.68 |
| | | | | | 29 | 3.4 | 38 | 0.002 | 0.86 |
| | | | | | 36 | 1.1 | 38 | 0.3 | 0.96 |
| 761.7011 (+) (lmm) | day | 147.87 | 6 | <0.0001 | -3 | 0.3 | 38 | 0.73 | 0.98 |
| | treatment | 8.16 | 1 | 0.005 | 1 | -3.5 | 38 | 0.001 | 1.2 |
| | day:treatment | 141.2 | 6 | <0.0001 | 8 | -2.2 | 38 | 0.034 | 1.12 |
| | | | | | 16 | 5.3 | 38 | <0.0001 | 0.76 |
| | | | | | 22 | 8 | 38 | <0.0001 | 0.64 |
| | | | | | 29 | 3.6 | 38 | 0.001 | 0.84 |
| | | | | | 36 | 1 | 38 | 0.32 | 0.96 |
| 762.7091 (+) (lmm) | day | 133.83 | 6 | <0.0001 | -3 | 0 | 37 | 1 | 1 |
| | treatment | 8.16 | 1 | 0.005 | 1 | -2.8 | 37 | 0.009 | 1.16 |
| | day:treatment | 124.88 | 6 | <0.0001 | 8 | -2.1 | 37 | 0.047 | 1.11 |
| | | | | | 16 | 4.9 | 37 | <0.0001 | 0.77 |
| | | | | | 22 | 7.8 | 37 | <0.0001 | 0.65 |
| | | | | | 29 | 3.7 | 37 | 0.001 | 0.83 |
| | | | | | 36 | 0.9 | 37 | 0.36 | 0.96 |
| 763.7131 (+) (lmm) | day | 109.94 | 6 | <0.0001 | -3 | 0 | 38 | 0.99 | 1 |
| | treatment | 12.9 | 1 | <0.0001 | 1 | -1.9 | 38 | 0.062 | 1.11 |
| | day:treatment | 119.8 | 6 | <0.0001 | 8 | -1.4 | 38 | 0.17 | 1.08 |
| | | | | | 16 | 5.3 | 38 | <0.0001 | 0.73 |

Continued on next page

Table 8 – continued from previous page

| Post-resistance pattern | | | | | | | | | |
|---|---|---|---|---|---|---|---|---|---|
| | lmm/lm | | | | Pairwise contrasts | | | | |
| **m/z peak** | *fixed effect* | $\chi^2/F$ | *df* | *adj. p* | *contrast* | *t-ratio* | *df* | *p* | *fold change* |
| | | | | | 22 | 8.3 | 38 | <0.0001 | 0.6 |
| | | | | | 29 | 4.1 | 38 | <0.0001 | 0.8 |
| | | | | | 36 | 1.1 | 38 | 0.28 | 0.95 |
| 766.6665 (+) (lmm) | day | 27.43 | 6 | <0.0001 | -3 | 1.9 | 51 | 0.063 | 0.88 |
| | treatment | 2.59 | 1 | 0.12 | 1 | 0.6 | 51 | 0.54 | 0.96 |
| | day:treatment | 11.26 | 6 | 0.088 | 8 | 2.4 | 51 | 0.02 | 0.82 |
| | | | | | 16 | -0.6 | 51 | 0.54 | 1.04 |
| | | | | | 22 | -1.1 | 51 | 0.29 | 1.07 |
| | | | | | 29 | 1.5 | 51 | 0.14 | 0.89 |
| | | | | | 36 | 1 | 51 | 0.34 | 0.94 |
| 767.5632 (+) (lmm) | day | 387.47 | 6 | <0.0001 | -3 | -1.4 | 41 | 0.17 | 1.28 |
| | treatment | 1.79 | 1 | 0.19 | 1 | 0.4 | 41 | 0.69 | 0.95 |
| | day:treatment | 10.91 | 6 | 0.099 | 8 | -0.3 | 41 | 0.73 | 1.03 |
| | | | | | 16 | -2.6 | 41 | 0.012 | 1.22 |
| | | | | | 22 | -1.7 | 41 | 0.09 | 1.13 |
| | | | | | 29 | -0.5 | 41 | 0.65 | 1.03 |
| | | | | | 36 | 0.6 | 41 | 0.58 | 0.95 |
| 768.5688 (+) (lmm) | day | 378.08 | 6 | <0.0001 | -3 | -1.1 | 37 | 0.28 | 1.18 |
| | treatment | 1.2 | 1 | 0.28 | 1 | 0.5 | 37 | 0.63 | 0.94 |
| | day:treatment | 13.72 | 6 | 0.037 | 8 | 0.1 | 37 | 0.93 | 0.99 |
| | | | | | 16 | -2.7 | 37 | 0.011 | 1.21 |
| | | | | | 22 | -2 | 37 | 0.054 | 1.15 |
| | | | | | 29 | -0.3 | 37 | 0.8 | 1.02 |
| | | | | | 36 | 0.6 | 37 | 0.53 | 0.94 |
| 769.5764 (+) (lmm) | day | 487.1 | 6 | <0.0001 | -3 | -1.8 | 51 | 0.081 | 1.23 |
| | treatment | 0.05 | 1 | 0.83 | 1 | 1.2 | 51 | 0.23 | 0.89 |
| | day:treatment | 12.54 | 6 | 0.056 | 8 | -0.2 | 51 | 0.85 | 1.01 |
| | | | | | 16 | -2.2 | 51 | 0.036 | 1.13 |
| | | | | | 22 | 0.6 | 51 | 0.57 | 0.97 |
| | | | | | 29 | 1.1 | 51 | 0.27 | 0.95 |
| | | | | | 36 | 0.4 | 51 | 0.66 | 0.97 |
| 776.6966 (+) (lmm) | day | 49.03 | 6 | <0.0001 | -3 | 0.8 | 22 | 0.41 | 0.96 |
| | treatment | 0.75 | 1 | 0.4 | 1 | -2.3 | 22 | 0.03 | 1.12 |
| | day:treatment | 151.99 | 6 | <0.0001 | 8 | -3.4 | 22 | 0.002 | 1.18 |
| | | | | | 16 | 4.8 | 22 | <0.0001 | 0.79 |
| | | | | | 22 | 5.5 | 22 | <0.0001 | 0.75 |
| | | | | | 29 | 1.3 | 22 | 0.21 | 0.94 |
| | | | | | 36 | -2 | 22 | 0.054 | 1.1 |
| 777.6983 (+) (lmm) | day | 44.25 | 6 | <0.0001 | -3 | 0.9 | 21 | 0.39 | 0.95 |
| | treatment | 1.33 | 1 | 0.26 | 1 | -2.1 | 21 | 0.043 | 1.11 |
| | day:treatment | 167.74 | 6 | <0.0001 | 8 | -3.3 | 21 | 0.004 | 1.18 |
| | | | | | 16 | 5.4 | 21 | <0.0001 | 0.75 |
| | | | | | 22 | 5.9 | 21 | <0.0001 | 0.72 |
| | | | | | 29 | 1.5 | 21 | 0.14 | 0.92 |
| | | | | | 36 | -2.1 | 21 | 0.051 | 1.11 |
| 784.5426 (+) (lmm) | day | 595.3 | 6 | <0.0001 | -3 | -1.7 | 41 | 0.092 | 1.26 |
| | treatment | 14.5 | 1 | <0.0001 | 1 | 0.5 | 41 | 0.6 | 0.94 |
| | day:treatment | 35.14 | 6 | <0.0001 | 8 | -4.1 | 41 | <0.0001 | 1.3 |
| | | | | | 16 | -5.1 | 41 | <0.0001 | 1.35 |
| | | | | | 22 | -3.7 | 41 | 0.001 | 1.24 |
| | | | | | 29 | -1.7 | 41 | 0.09 | 1.1 |
| | | | | | 36 | -0.1 | 41 | 0.95 | 1 |
| 788.6551 (+) (lmm) | day | 34.5 | 6 | <0.0001 | -3 | 1.1 | 50 | 0.28 | 0.92 |
| | treatment | 2.22 | 1 | 0.14 | 1 | 1.1 | 50 | 0.29 | 0.92 |
| | day:treatment | 24.93 | 6 | <0.0001 | 8 | 3.6 | 50 | 0.001 | 0.74 |
| | | | | | 16 | -0.8 | 50 | 0.46 | 1.05 |
| | | | | | 22 | -2.4 | 50 | 0.021 | 1.17 |
| | | | | | 29 | 1.2 | 50 | 0.24 | 0.92 |
| | | | | | 36 | 1.6 | 50 | 0.11 | 0.89 |
| 793.5764 (-) (lmm) | day | 113.32 | 6 | <0.0001 | -3 | 1.9 | 56 | 0.062 | 0.92 |
| | treatment | 5.96 | 1 | 0.017 | 1 | -3.6 | 56 | 0.001 | 1.15 |
| | day:treatment | 54.75 | 6 | <0.0001 | 8 | -2.6 | 56 | 0.013 | 1.1 |
| | | | | | 16 | 1.1 | 56 | 0.27 | 0.96 |
| | | | | | 22 | 4.2 | 56 | <0.0001 | 0.86 |
| | | | | | 29 | 3.9 | 56 | <0.0001 | 0.87 |
| | | | | | 36 | 1.8 | 56 | 0.071 | 0.94 |
| 794.5791 (-) (lmm) | day | 98.57 | 6 | <0.0001 | -3 | 1.9 | 56 | 0.06 | 0.92 |
| | treatment | 10.85 | 1 | 0.001 | 1 | -3.5 | 56 | 0.001 | 1.14 |
| | day:treatment | 62.83 | 6 | <0.0001 | 8 | -2.5 | 56 | 0.014 | 1.1 |
| | | | | | 16 | 1.5 | 56 | 0.13 | 0.94 |
| | | | | | 22 | 5 | 56 | <0.0001 | 0.83 |
| | | | | | 29 | 4.4 | 56 | <0.0001 | 0.85 |
| | | | | | 36 | 2.3 | 56 | 0.027 | 0.92 |
| 798.6798 (+) (lmm) | day | 46.75 | 6 | <0.0001 | -3 | 1.9 | 30 | 0.068 | 0.9 |
| | treatment | 3.45 | 1 | 0.069 | 1 | -1.9 | 30 | 0.062 | 1.1 |
| | day:treatment | 57.95 | 6 | <0.0001 | 8 | -1.9 | 30 | 0.064 | 1.1 |
| | | | | | 16 | 4 | 30 | <0.0001 | 0.83 |

Continued on next page

Table 8 – continued from previous page

| Post-resistance pattern | | | | | | | | | |
|---|---|---|---|---|---|---|---|---|---|
| | lmm/lm | | | | Pairwise contrasts | | | | |
| **m/z peak** | *fixed effect* | $\chi^2/F$ | *df* | *adj. p* | *contrast* | *t-ratio* | *df* | *p* | *fold change* |
| | | | | | 22 | 3.7 | 30 | 0.001 | 0.83 |
| | | | | | 29 | 2.5 | 30 | 0.02 | 0.89 |
| | | | | | 36 | 0.7 | 30 | 0.48 | 0.97 |
| 800.6925 (+) (lmm) | day | 40.2 | 6 | <0.0001 | -3 | 1.5 | 27 | 0.14 | 0.91 |
| | treatment | 10.36 | 1 | 0.002 | 1 | -2.5 | 27 | 0.02 | 1.14 |
| | day:treatment | 149.75 | 6 | <0.0001 | 8 | -0.3 | 27 | 0.78 | 1.02 |
| | | | | | 16 | 7.8 | 27 | <0.0001 | 0.63 |
| | | | | | 22 | 6.8 | 27 | <0.0001 | 0.67 |
| | | | | | 29 | 2.6 | 27 | 0.014 | 0.86 |
| | | | | | 36 | 0 | 27 | 0.97 | 1 |
| 807.5381 (-) (lmm) | day | 161.53 | 6 | <0.0001 | -3 | -1.1 | 44 | 0.26 | 1.15 |
| | treatment | 5.78 | 1 | 0.018 | 1 | 1.8 | 44 | 0.074 | 0.81 |
| | day:treatment | 87.66 | 6 | <0.0001 | 8 | 1.2 | 44 | 0.22 | 0.88 |
| | | | | | 16 | -5.5 | 44 | <0.0001 | 1.55 |
| | | | | | 22 | -6.7 | 44 | <0.0001 | 1.62 |
| | | | | | 29 | -0.2 | 44 | 0.87 | 1.01 |
| | | | | | 36 | 0.8 | 44 | 0.46 | 0.92 |
| 821.5621 (-) (lmm) | day | 354.4 | 6 | <0.0001 | -3 | -1.1 | 46 | 0.29 | 1.06 |
| | treatment | 21.24 | 1 | <0.0001 | 1 | 0.5 | 46 | 0.63 | 0.98 |
| | day:treatment | 146.24 | 6 | <0.0001 | 8 | 1 | 46 | 0.32 | 0.96 |
| | | | | | 16 | -7.2 | 46 | <0.0001 | 1.32 |
| | | | | | 22 | -10.4 | 46 | <0.0001 | 1.45 |
| | | | | | 29 | -1.4 | 46 | 0.16 | 1.06 |
| | | | | | 36 | 0.7 | 46 | 0.46 | 0.97 |
| 824.5551 (-) (lmm) | day | 145.32 | 6 | <0.0001 | -3 | -1.5 | 50 | 0.14 | 1.08 |
| | treatment | 1.44 | 1 | 0.24 | 1 | 4.6 | 50 | <0.0001 | 0.78 |
| | day:treatment | 107.72 | 6 | <0.0001 | 8 | 4.7 | 50 | <0.0001 | 0.79 |
| | | | | | 16 | -3.7 | 50 | 0.001 | 1.17 |
| | | | | | 22 | -4.6 | 50 | <0.0001 | 1.21 |
| | | | | | 29 | 1.3 | 50 | 0.2 | 0.94 |
| | | | | | 36 | 3.5 | 50 | 0.001 | 0.83 |
| 831.5377 (+) (lmm) | day | 88.37 | 6 | <0.0001 | -3 | -1.1 | 49 | 0.3 | 1.23 |
| | treatment | 11.08 | 1 | 0.001 | 1 | 1.5 | 49 | 0.14 | 0.76 |
| | day:treatment | 94.67 | 6 | <0.0001 | 8 | 1.9 | 49 | 0.066 | 0.73 |
| | | | | | 16 | -6.6 | 49 | <0.0001 | 2.21 |
| | | | | | 22 | -7.1 | 49 | <0.0001 | 2.16 |
| | | | | | 29 | -1.1 | 49 | 0.3 | 1.16 |
| | | | | | 36 | 0.1 | 49 | 0.93 | 0.98 |
| 847.5316 (+) (lmm) | day | 75.11 | 6 | <0.0001 | -3 | -0.8 | 51 | 0.44 | 1.23 |
| | treatment | 11.5 | 1 | 0.001 | 1 | 1.2 | 51 | 0.26 | 0.76 |
| | day:treatment | 79.43 | 6 | <0.0001 | 8 | 1.7 | 51 | 0.091 | 0.67 |
| | | | | | 16 | -6.3 | 51 | <0.0001 | 2.56 |
| | | | | | 22 | -6.6 | 51 | <0.0001 | 2.41 |
| | | | | | 29 | -1 | 51 | 0.33 | 1.2 |
| | | | | | 36 | 0 | 51 | 0.98 | 1 |
| 848.5355 (+) (lmm) | day | 62.2 | 6 | <0.0001 | -3 | -0.5 | 52 | 0.63 | 1.12 |
| | treatment | 9.88 | 1 | 0.002 | 1 | 1 | 52 | 0.3 | 0.79 |
| | day:treatment | 68.88 | 6 | <0.0001 | 8 | 1.4 | 52 | 0.16 | 0.73 |
| | | | | | 16 | -5.8 | 52 | <0.0001 | 2.37 |
| | | | | | 22 | -6.3 | 52 | <0.0001 | 2.31 |
| | | | | | 29 | -0.8 | 52 | 0.46 | 1.15 |
| | | | | | 36 | 0.1 | 52 | 0.93 | 0.98 |
| 849.5403 (+) (lmm) | day | 48.66 | 6 | <0.0001 | -3 | -0.5 | 54 | 0.59 | 1.13 |
| | treatment | 6.77 | 1 | 0.011 | 1 | 1.2 | 54 | 0.23 | 0.78 |
| | day:treatment | 56.83 | 6 | <0.0001 | 8 | 1.7 | 54 | 0.088 | 0.7 |
| | | | | | 16 | -5.6 | 54 | <0.0001 | 2.21 |
| | | | | | 22 | -5.1 | 54 | <0.0001 | 1.95 |
| | | | | | 29 | -0.2 | 54 | 0.84 | 1.04 |
| | | | | | 36 | 0.3 | 54 | 0.76 | 0.94 |
| 857.5345 (-) (lmm) | day | 453.03 | 6 | <0.0001 | -3 | -1.3 | 40 | 0.21 | 1.12 |
| | treatment | 0.81 | 1 | 0.38 | 1 | 1.8 | 40 | 0.081 | 0.87 |
| | day:treatment | 31.71 | 6 | <0.0001 | 8 | -1 | 40 | 0.31 | 1.06 |
| | | | | | 16 | -3 | 40 | 0.005 | 1.17 |
| | | | | | 22 | -2.8 | 40 | 0.008 | 1.14 |
| | | | | | 29 | 1.2 | 40 | 0.24 | 0.94 |
| | | | | | 36 | 1.2 | 40 | 0.22 | 0.92 |
| 905.5121 (-) (lmm) | day | 62.63 | 6 | <0.0001 | -3 | -0.8 | 42 | 0.42 | 1.09 |
| | treatment | 17.21 | 1 | <0.0001 | 1 | 0.4 | 42 | 0.72 | 0.97 |
| | day:treatment | 80.05 | 6 | <0.0001 | 8 | -0.2 | 42 | 0.83 | 1.02 |
| | | | | | 16 | -6.8 | 42 | <0.0001 | 1.73 |
| | | | | | 22 | -7.2 | 42 | <0.0001 | 1.75 |
| | | | | | 29 | -1.5 | 42 | 0.13 | 1.15 |
| | | | | | 36 | -0.7 | 42 | 0.48 | 1.08 |
| 969.6719 (+) (lm) | day | 12.58 | 6,56 | <0.0001 | -3 | 1.8 | 56 | 0.083 | 0.88 |
| | treatment | 13.83 | 1,56 | 0.001 | 1 | 2.7 | 56 | 0.008 | 0.81 |
| | day:treatment | 1.95 | 6,56 | 0.095 | 8 | 3.1 | 56 | 0.003 | 0.72 |
| | | | | | 16 | -0.5 | 56 | 0.64 | 1.05 |

Continued on next page

Table 8 – continued from previous page

| Post-resistance pattern | | | | | | | | | |
|---|---|---|---|---|---|---|---|---|---|
| | lmm/lm | | | | Pairwise contrasts | | | | |
| **m/z peak** | *fixed effect* | $\chi^2/F$ | *df* | *adj. p* | *contrast* | *t-ratio* | *df* | *p* | *fold change* |
| | | | | | 22 | -0.4 | 56 | 0.72 | 1.03 |
| | | | | | 29 | 1.2 | 56 | 0.23 | 0.88 |
| | | | | | 36 | 1.8 | 56 | 0.074 | 0.85 |
| 985.6313 (-) (lmm) | day | 37.25 | 6 | <0.0001 | -3 | -0.8 | 45 | 0.44 | 1.03 |
| | treatment | 12.65 | 1 | <0.0001 | 1 | 2.4 | 45 | 0.019 | 0.9 |
| | day:treatment | 46.5 | 6 | <0.0001 | 8 | 6.4 | 45 | <0.0001 | 0.74 |
| | | | | | 16 | 1 | 45 | 0.31 | 0.96 |
| | | | | | 22 | -0.7 | 45 | 0.49 | 1.03 |
| | | | | | 29 | 2.4 | 45 | 0.021 | 0.89 |
| | | | | | 36 | 3.1 | 45 | 0.003 | 0.87 |
| 987.6336 (-) (lmm) | day | 29.62 | 6 | <0.0001 | -3 | 0.2 | 45 | 0.88 | 0.99 |
| | treatment | 16.47 | 1 | <0.0001 | 1 | 2.4 | 45 | 0.023 | 0.89 |
| | day:treatment | 25.87 | 6 | <0.0001 | 8 | 5.2 | 45 | <0.0001 | 0.76 |
| | | | | | 16 | 1.6 | 45 | 0.11 | 0.92 |
| | | | | | 22 | 0.2 | 45 | 0.88 | 0.99 |
| | | | | | 29 | 2.8 | 45 | 0.008 | 0.86 |
| | | | | | 36 | 3.7 | 45 | 0.001 | 0.83 |

Table 9: Analysis of Deviance Table (Type II Wald $\chi^2$-tests) for linear mixed effects model for all m/z peaks of interest with persistent pattern from exploratory analysis with percentage ion count (arc sine transformed) of the m/z peak as response. Where the lmm yielded a singular fit, an lm has been fit instead. P-values have been adjusted using the Benjamini-Hochberg procedure. Pairwise contrasts by day, as estimated by the `emmeans` package for R.

| Persistent pattern | | | | | | | | | |
|---|---|---|---|---|---|---|---|---|---|
| | lmm/lm | | | | Pairwise contrasts | | | | |
| **m/z peak** | *fixed effect* | $\chi^2/F$ | *df* | *adj. p* | *contrast* | *t-ratio* | *df* | *p* | *fold change* |
| 157.0178 (-) (lm) | day | 9.28 | 6,56 | <0.0001 | -3 | -0.9 | 56 | 0.4 | 1.04 |
| | treatment | 233.27 | 1,56 | <0.0001 | 1 | 4.1 | 56 | <0.0001 | 0.82 |
| | day:treatment | 11.42 | 6,56 | <0.0001 | 8 | 7.6 | 56 | <0.0001 | 0.66 |
| | | | | | 16 | 9.3 | 56 | <0.0001 | 0.58 |
| | | | | | 22 | 7.6 | 56 | <0.0001 | 0.65 |
| | | | | | 29 | 5.3 | 56 | <0.0001 | 0.77 |
| | | | | | 36 | 7.4 | 56 | <0.0001 | 0.69 |
| 174.9695 (-) (lmm) | day | 89.25 | 6 | <0.0001 | -3 | -0.6 | 45 | 0.58 | 1.03 |
| | treatment | 54.64 | 1 | <0.0001 | 1 | 4.6 | 45 | <0.0001 | 0.77 |
| | day:treatment | 51.36 | 6 | <0.0001 | 8 | 5.3 | 45 | <0.0001 | 0.72 |
| | | | | | 16 | 7.3 | 45 | <0.0001 | 0.61 |
| | | | | | 22 | 6.2 | 45 | <0.0001 | 0.67 |
| | | | | | 29 | 3.9 | 45 | <0.0001 | 0.79 |
| | | | | | 36 | 2.4 | 45 | 0.021 | 0.87 |
| 266.0025 (-) (lmm) | day | 277.99 | 6 | <0.0001 | -3 | 1.6 | 38 | 0.12 | 0.94 |
| | treatment | 77.71 | 1 | <0.0001 | 1 | 11.1 | 38 | <0.0001 | 0.61 |
| | day:treatment | 91.39 | 6 | <0.0001 | 8 | 3.6 | 38 | 0.001 | 0.85 |
| | | | | | 16 | 7.8 | 38 | <0.0001 | 0.66 |
| | | | | | 22 | 7.1 | 38 | <0.0001 | 0.68 |
| | | | | | 29 | 3.9 | 38 | <0.0001 | 0.81 |
| | | | | | 36 | 3.1 | 38 | 0.003 | 0.85 |
| 472.4163 (+) (lmm) | day | 44.6 | 6 | <0.0001 | -3 | -0.5 | 26 | 0.63 | 1.05 |
| | treatment | 16.56 | 1 | <0.0001 | 1 | -2.4 | 26 | 0.024 | 1.29 |
| | day:treatment | 18.28 | 6 | 0.007 | 8 | -2.6 | 26 | 0.015 | 1.31 |
| | | | | | 16 | -4.3 | 26 | <0.0001 | 1.45 |
| | | | | | 22 | -4.2 | 26 | <0.0001 | 1.44 |
| | | | | | 29 | -3.5 | 26 | 0.002 | 1.42 |
| | | | | | 36 | -3.2 | 26 | 0.004 | 1.39 |
| 494.4037 (+) (lmm) | day | 46.67 | 6 | <0.0001 | -3 | -1.1 | 25 | 0.28 | 1.13 |
| | treatment | 31.82 | 1 | <0.0001 | 1 | -2.5 | 25 | 0.021 | 1.3 |
| | day:treatment | 31.38 | 6 | <0.0001 | 8 | -4.1 | 25 | <0.0001 | 1.56 |
| | | | | | 16 | -5.2 | 25 | <0.0001 | 1.64 |
| | | | | | 22 | -5.3 | 25 | <0.0001 | 1.6 |
| | | | | | 29 | -5.5 | 25 | <0.0001 | 1.72 |
| | | | | | 36 | -5 | 25 | <0.0001 | 1.79 |
| 496.4186 (+) (lmm) | day | 40.34 | 6 | <0.0001 | -3 | -1.5 | 34 | 0.14 | 1.12 |
| | treatment | 37.6 | 1 | <0.0001 | 1 | -3.7 | 34 | 0.001 | 1.29 |
| | day:treatment | 25.78 | 6 | <0.0001 | 8 | -5.5 | 34 | <0.0001 | 1.43 |
| | | | | | 16 | -3.2 | 34 | 0.003 | 1.22 |
| | | | | | 22 | -2.5 | 34 | 0.019 | 1.16 |
| | | | | | 29 | -5.5 | 34 | <0.0001 | 1.4 |
| | | | | | 36 | -6 | 34 | <0.0001 | 1.52 |
| 497.4213 (+) (lmm) | day | 34.07 | 6 | <0.0001 | -3 | -1.5 | 33 | 0.13 | 1.14 |
| | treatment | 30.87 | 1 | <0.0001 | 1 | -3.3 | 33 | 0.002 | 1.31 |
| | day:treatment | 21.89 | 6 | 0.002 | 8 | -5 | 33 | <0.0001 | 1.46 |

Continued on next page

Table 9 – continued from previous page

| Persistent pattern | | | | | | | | | |
|---|---|---|---|---|---|---|---|---|---|
| | lmm/lm | | | | Pairwise contrasts | | | | |
| **m/z peak** | *fixed effect* | $\chi^2/F$ | *df* | *adj. p* | *contrast* | *t-ratio* | *df* | *p* | *fold change* |
| | | | | | 16 | -2.9 | 33 | 0.006 | 1.24 |
| | | | | | 22 | -2.3 | 33 | 0.029 | 1.18 |
| | | | | | 29 | -5.1 | 33 | <0.0001 | 1.45 |
| | | | | | 36 | -5.5 | 33 | <0.0001 | 1.58 |
| 542.4081 (-) (lmm) | day | 9.79 | 6 | 0.14 | -3 | -1.9 | 48 | 0.069 | 1.15 |
| | treatment | 10.54 | 1 | 0.001 | 1 | -3 | 48 | 0.005 | 1.24 |
| | day:treatment | 67.42 | 6 | <0.0001 | 8 | -3.1 | 48 | 0.003 | 1.25 |
| | | | | | 16 | 2.6 | 48 | 0.011 | 0.81 |
| | | | | | 22 | 2.4 | 48 | 0.02 | 0.83 |
| | | | | | 29 | -4.1 | 48 | <0.0001 | 1.34 |
| | | | | | 36 | -5.1 | 48 | <0.0001 | 1.48 |
| 555.3261 (-) (lmm) | day | 28.64 | 6 | <0.0001 | -3 | 0.3 | 25 | 0.79 | 0.97 |
| | treatment | 24.85 | 1 | <0.0001 | 1 | -3.8 | 25 | 0.001 | 1.54 |
| | day:treatment | 40.83 | 6 | <0.0001 | 8 | -2.7 | 25 | 0.013 | 1.5 |
| | | | | | 16 | -4.5 | 25 | <0.0001 | 1.81 |
| | | | | | 22 | -4.5 | 25 | <0.0001 | 1.84 |
| | | | | | 29 | -4.9 | 25 | <0.0001 | 2.02 |
| | | | | | 36 | -5.4 | 25 | <0.0001 | 2.21 |
| 556.3296 (-) (lmm) | day | 27.89 | 6 | <0.0001 | -3 | 0.3 | 24 | 0.8 | 0.97 |
| | treatment | 19.66 | 1 | <0.0001 | 1 | -3.5 | 24 | 0.002 | 1.54 |
| | day:treatment | 36.69 | 6 | <0.0001 | 8 | -2.2 | 24 | 0.036 | 1.45 |
| | | | | | 16 | -4.1 | 24 | <0.0001 | 1.79 |
| | | | | | 22 | -4.2 | 24 | <0.0001 | 1.84 |
| | | | | | 29 | -4.4 | 24 | <0.0001 | 1.99 |
| | | | | | 36 | -4.9 | 24 | <0.0001 | 2.18 |
| 571.3289 (-) (lmm) | day | 6.99 | 6 | 0.33 | -3 | 0.1 | 21 | 0.94 | 0.99 |
| | treatment | 13.55 | 1 | <0.0001 | 1 | -2.3 | 21 | 0.035 | 1.45 |
| | day:treatment | 31.53 | 6 | <0.0001 | 8 | -2.8 | 21 | 0.011 | 1.66 |
| | | | | | 16 | -3.3 | 21 | 0.004 | 1.68 |
| | | | | | 22 | -2.8 | 21 | 0.012 | 1.53 |
| | | | | | 29 | -4.1 | 21 | 0.001 | 1.93 |
| | | | | | 36 | -4.8 | 21 | <0.0001 | 2.23 |
| 585.5159 (+) (lmm) | day | 146.86 | 6 | <0.0001 | -3 | -1 | 34 | 0.35 | 1.04 |
| | treatment | 17.92 | 1 | <0.0001 | 1 | 3.5 | 34 | 0.001 | 0.88 |
| | day:treatment | 29.73 | 6 | <0.0001 | 8 | 4.2 | 34 | <0.0001 | 0.87 |
| | | | | | 16 | 2.3 | 34 | 0.03 | 0.93 |
| | | | | | 22 | 3.9 | 34 | <0.0001 | 0.88 |
| | | | | | 29 | 4 | 34 | <0.0001 | 0.89 |
| | | | | | 36 | 2.2 | 34 | 0.033 | 0.93 |
| 587.5342 (+) (lmm) | day | 81.35 | 6 | <0.0001 | -3 | 0.5 | 32 | 0.6 | 0.98 |
| | treatment | 62.04 | 1 | <0.0001 | 1 | 4.7 | 32 | <0.0001 | 0.84 |
| | day:treatment | 66.48 | 6 | <0.0001 | 8 | 4.3 | 32 | <0.0001 | 0.87 |
| | | | | | 16 | 6.8 | 32 | <0.0001 | 0.79 |
| | | | | | 22 | 9 | 32 | <0.0001 | 0.73 |
| | | | | | 29 | 6.9 | 32 | <0.0001 | 0.8 |
| | | | | | 36 | 4.5 | 32 | <0.0001 | 0.86 |
| 588.5368 (+) (lmm) | day | 60.6 | 6 | <0.0001 | -3 | 0.5 | 32 | 0.6 | 0.98 |
| | treatment | 77.98 | 1 | <0.0001 | 1 | 5.5 | 32 | <0.0001 | 0.82 |
| | day:treatment | 79.48 | 6 | <0.0001 | 8 | 5.2 | 32 | <0.0001 | 0.84 |
| | | | | | 16 | 7.7 | 32 | <0.0001 | 0.76 |
| | | | | | 22 | 9.7 | 32 | <0.0001 | 0.71 |
| | | | | | 29 | 7.7 | 32 | <0.0001 | 0.78 |
| | | | | | 36 | 4.8 | 32 | <0.0001 | 0.85 |
| 589.5486 (+) (lmm) | day | 71.57 | 6 | <0.0001 | -3 | 1 | 21 | 0.34 | 0.96 |
| | treatment | 35.12 | 1 | <0.0001 | 1 | 3.6 | 21 | 0.002 | 0.86 |
| | day:treatment | 58.53 | 6 | <0.0001 | 8 | 3.6 | 21 | 0.002 | 0.87 |
| | | | | | 16 | 6 | 21 | <0.0001 | 0.78 |
| | | | | | 22 | 7.5 | 21 | <0.0001 | 0.73 |
| | | | | | 29 | 6 | 21 | <0.0001 | 0.79 |
| | | | | | 36 | 4.2 | 21 | <0.0001 | 0.85 |
| 615.4999 (+) (lmm) | day | 172.78 | 6 | <0.0001 | -3 | -0.3 | 45 | 0.8 | 1 |
| | treatment | 110.01 | 1 | <0.0001 | 1 | 4.4 | 45 | <0.0001 | 0.92 |
| | day:treatment | 177.94 | 6 | <0.0001 | 8 | 13.5 | 45 | <0.0001 | 0.75 |
| | | | | | 16 | 11.5 | 45 | <0.0001 | 0.78 |
| | | | | | 22 | 3.6 | 45 | 0.001 | 0.93 |
| | | | | | 29 | 5 | 45 | <0.0001 | 0.91 |
| | | | | | 36 | 3.4 | 45 | 0.002 | 0.93 |
| 702.2965 (+) (lmm) | day | 23.65 | 6 | 0.001 | -3 | -1.2 | 56 | 0.23 | 1.1 |
| | treatment | 33.31 | 1 | <0.0001 | 1 | 3.4 | 56 | 0.001 | 0.75 |
| | day:treatment | 16.84 | 6 | 0.011 | 8 | 3.7 | 56 | 0.001 | 0.72 |
| | | | | | 16 | 3.1 | 56 | 0.003 | 0.74 |
| | | | | | 22 | 2.7 | 56 | 0.01 | 0.79 |
| | | | | | 29 | 2.3 | 56 | 0.023 | 0.83 |
| | | | | | 36 | 2.7 | 56 | 0.01 | 0.8 |
| 735.6856 (+) (lmm) | day | 182.35 | 6 | <0.0001 | -3 | 0.6 | 45 | 0.57 | 0.98 |
| | treatment | 8.14 | 1 | 0.005 | 1 | -5.1 | 45 | <0.0001 | 1.2 |
| | day:treatment | 146.41 | 6 | <0.0001 | 8 | -2.7 | 45 | 0.011 | 1.1 |

Continued on next page

Table 9 – continued from previous page

| Persistent pattern | | | | | | | | | |
|---|---|---|---|---|---|---|---|---|---|
| | **lmm/lm** | | | | **Pairwise contrasts** | | | | |
| **m/z peak** | *fixed effect* | $\chi^2/F$ | *df* | *adj. p* | *contrast* | *t-ratio* | *df* | *p* | *fold change* |
| | | | | | 16 | 4.6 | 45 | <0.0001 | 0.84 |
| | | | | | 22 | 7.7 | 45 | <0.0001 | 0.75 |
| | | | | | 29 | 3.5 | 45 | 0.001 | 0.89 |
| | | | | | 36 | 2.6 | 45 | 0.013 | 0.92 |
| 776.3239 (+) (lmm) | day | 26.66 | 6 | <0.0001 | -3 | -1.5 | 54 | 0.13 | 1.12 |
| | treatment | 29 | 1 | <0.0001 | 1 | 3.8 | 54 | <0.0001 | 0.73 |
| | day:treatment | 22.79 | 6 | 0.001 | 8 | 3.9 | 54 | <0.0001 | 0.72 |
| | | | | | 16 | 3.1 | 54 | 0.003 | 0.75 |
| | | | | | 22 | 2.4 | 54 | 0.019 | 0.81 |
| | | | | | 29 | 2.5 | 54 | 0.014 | 0.82 |
| | | | | | 36 | 3.2 | 54 | 0.002 | 0.77 |
| 812.648 (+) (lm) | day | 3.72 | 6,56 | 0.004 | -3 | 0.3 | 56 | 0.74 | 0.98 |
| | treatment | 112.71 | 1,56 | <0.0001 | 1 | 3.9 | 56 | <0.0001 | 0.81 |
| | day:treatment | 5.27 | 6,56 | <0.0001 | 8 | 7.8 | 56 | <0.0001 | 0.61 |
| | | | | | 16 | 3 | 56 | 0.004 | 0.84 |
| | | | | | 22 | 3.2 | 56 | 0.002 | 0.84 |
| | | | | | 29 | 4.8 | 56 | <0.0001 | 0.75 |
| | | | | | 36 | 5.1 | 56 | <0.0001 | 0.77 |
| 825.559 (-) (lm) | day | 9.28 | 6,56 | <0.0001 | -3 | -0.8 | 56 | 0.4 | 1.05 |
| | treatment | 9.72 | 1,56 | 0.003 | 1 | 4.2 | 56 | <0.0001 | 0.78 |
| | day:treatment | 10.7 | 6,56 | <0.0001 | 8 | 5.1 | 56 | <0.0001 | 0.74 |
| | | | | | 16 | -2.5 | 56 | 0.016 | 1.13 |
| | | | | | 22 | -3 | 56 | 0.004 | 1.16 |
| | | | | | 29 | 2 | 56 | 0.045 | 0.89 |
| | | | | | 36 | 3.3 | 56 | 0.002 | 0.82 |
| 826.5628 (-) (lm) | day | 8.3 | 6,56 | <0.0001 | -3 | -1 | 56 | 0.32 | 1.06 |
| | treatment | 16.93 | 1,56 | <0.0001 | 1 | 4.6 | 56 | <0.0001 | 0.78 |
| | day:treatment | 11.36 | 6,56 | <0.0001 | 8 | 5.6 | 56 | <0.0001 | 0.74 |
| | | | | | 16 | -2.2 | 56 | 0.032 | 1.11 |
| | | | | | 22 | -2.4 | 56 | 0.019 | 1.12 |
| | | | | | 29 | 2.5 | 56 | 0.014 | 0.88 |
| | | | | | 36 | 3.7 | 56 | 0.001 | 0.82 |
| 926.6889 (+) (lmm) | day | 19.39 | 6 | 0.004 | -3 | -0.6 | 35 | 0.56 | 1.03 |
| | treatment | 65.58 | 1 | <0.0001 | 1 | 5.2 | 35 | <0.0001 | 0.75 |
| | day:treatment | 65.73 | 6 | <0.0001 | 8 | 8.2 | 35 | <0.0001 | 0.63 |
| | | | | | 16 | 6.2 | 35 | <0.0001 | 0.69 |
| | | | | | 22 | 6.2 | 35 | <0.0001 | 0.69 |
| | | | | | 29 | 6 | 35 | <0.0001 | 0.7 |
| | | | | | 36 | 5.2 | 35 | <0.0001 | 0.74 |
| 928.7071 (+) (lmm) | day | 25.7 | 6 | <0.0001 | -3 | 0.7 | 37 | 0.52 | 0.97 |
| | treatment | 90.34 | 1 | <0.0001 | 1 | 5.1 | 37 | <0.0001 | 0.76 |
| | day:treatment | 56.95 | 6 | <0.0001 | 8 | 6.3 | 37 | <0.0001 | 0.72 |
| | | | | | 16 | 8 | 37 | <0.0001 | 0.63 |
| | | | | | 22 | 8.4 | 37 | <0.0001 | 0.61 |
| | | | | | 29 | 7 | 37 | <0.0001 | 0.67 |
| | | | | | 36 | 6.5 | 37 | <0.0001 | 0.7 |
| 930.7205 (+) (lmm) | day | 23.57 | 6 | 0.001 | -3 | 0.4 | 35 | 0.66 | 0.98 |
| | treatment | 60.15 | 1 | <0.0001 | 1 | 4.2 | 35 | <0.0001 | 0.79 |
| | day:treatment | 41.22 | 6 | <0.0001 | 8 | 5.4 | 35 | <0.0001 | 0.75 |
| | | | | | 16 | 6.2 | 35 | <0.0001 | 0.7 |
| | | | | | 22 | 6.5 | 35 | <0.0001 | 0.69 |
| | | | | | 29 | 6.3 | 35 | <0.0001 | 0.7 |
| | | | | | 36 | 6.1 | 35 | <0.0001 | 0.73 |
| 932.7325 (+) (lmm) | day | 146.61 | 6 | <0.0001 | -3 | 0.8 | 35 | 0.42 | 0.96 |
| | treatment | 67.45 | 1 | <0.0001 | 1 | 5 | 35 | <0.0001 | 0.78 |
| | day:treatment | 47.97 | 6 | <0.0001 | 8 | 5.3 | 35 | <0.0001 | 0.76 |
| | | | | | 16 | 6.8 | 35 | <0.0001 | 0.67 |
| | | | | | 22 | 8.5 | 35 | <0.0001 | 0.62 |
| | | | | | 29 | 5 | 35 | <0.0001 | 0.77 |
| | | | | | 36 | 5.6 | 35 | <0.0001 | 0.77 |
| 934.7432 (+) (lmm) | day | 141.87 | 6 | <0.0001 | -3 | 0.7 | 36 | 0.47 | 0.96 |
| | treatment | 78.39 | 1 | <0.0001 | 1 | 4.9 | 36 | <0.0001 | 0.77 |
| | day:treatment | 76.05 | 6 | <0.0001 | 8 | 5.1 | 36 | <0.0001 | 0.75 |
| | | | | | 16 | 8.3 | 36 | <0.0001 | 0.58 |
| | | | | | 22 | 10.2 | 36 | <0.0001 | 0.53 |
| | | | | | 29 | 5.3 | 36 | <0.0001 | 0.74 |
| | | | | | 36 | 4.9 | 36 | <0.0001 | 0.79 |
| 962.6851 (-) (lmm) | day | 112.35 | 6 | <0.0001 | -3 | 0.4 | 45 | 0.67 | 0.98 |
| | treatment | 62.1 | 1 | <0.0001 | 1 | 2.1 | 45 | 0.041 | 0.91 |
| | day:treatment | 71.26 | 6 | <0.0001 | 8 | 3.5 | 45 | 0.001 | 0.86 |
| | | | | | 16 | 8.4 | 45 | <0.0001 | 0.66 |
| | | | | | 22 | 8.7 | 45 | <0.0001 | 0.66 |
| | | | | | 29 | 4 | 45 | <0.0001 | 0.84 |
| | | | | | 36 | 3.9 | 45 | <0.0001 | 0.86 |

Table 10: Analysis of Deviance Table (Type II Wald $\chi^2$-tests) for linear mixed effects model for all m/z peaks of interest with unclassified fluctuating pattern from exploratory analysis with percentage ion count (arc sine transformed) of the m/z peak as response. Where the lmm yielded a singular fit, an lm has been fit instead. P-values have been adjusted using the Benjamini-Hochberg procedure. Pairwise contrasts by day, as estimated by the `emmeans` package for R.

| **Unclassified fluctuating pattern** | | | | | | | | | |
|---|---|---|---|---|---|---|---|---|---|
| | **lmm/lm** | | | | **Pairwise contrasts** | | | | |
| **m/z peak** | *fixed effect* | $\chi^2/F$ | *df* | *adj. p* | *contrast* | *t-ratio* | *df* | *p* | *fold change* |
| 245.0606 (-) (lmm) | day | 5.38 | 6 | 0.5 | -3 | 1 | 40 | 0.34 | 0.93 |
| | treatment | 2.54 | 1 | 0.12 | 1 | -3.7 | 40 | 0.001 | 1.3 |
| | day:treatment | 28.68 | 6 | <0.0001 | 8 | -1.9 | 40 | 0.059 | 1.15 |
| | | | | | 16 | -0.7 | 40 | 0.48 | 1.05 |
| | | | | | 22 | 1.6 | 40 | 0.12 | 0.89 |
| | | | | | 29 | -0.5 | 40 | 0.62 | 1.04 |
| | | | | | 36 | -2.4 | 40 | 0.019 | 1.19 |
| 317.2156 (+) (lm) | day | 28.56 | 6,56 | <0.0001 | -3 | 1.6 | 56 | 0.12 | 0.96 |
| | treatment | 0.06 | 1,56 | 0.81 | 1 | -0.6 | 56 | 0.55 | 1.02 |
| | day:treatment | 3.55 | 6,56 | 0.006 | 8 | 1.8 | 56 | 0.079 | 0.94 |
| | | | | | 16 | -2.8 | 56 | 0.007 | 1.1 |
| | | | | | 22 | 1.2 | 56 | 0.22 | 0.96 |
| | | | | | 29 | 0.5 | 56 | 0.6 | 0.98 |
| | | | | | 36 | -2.4 | 56 | 0.022 | 1.08 |
| 333.0886 (-) (lmm) | day | 38.04 | 6 | <0.0001 | -3 | 2 | 53 | 0.054 | 0.81 |
| | treatment | 9.08 | 1 | 0.003 | 1 | -4.6 | 53 | <0.0001 | 1.66 |
| | day:treatment | 38.29 | 6 | <0.0001 | 8 | -1.8 | 53 | 0.078 | 1.31 |
| | | | | | 16 | -0.7 | 53 | 0.49 | 1.1 |
| | | | | | 22 | 0.4 | 53 | 0.68 | 0.94 |
| | | | | | 29 | -0.8 | 53 | 0.41 | 1.12 |
| | | | | | 36 | -4.4 | 53 | <0.0001 | 1.73 |
| 991.664 (-) (lm) | day | 10.92 | 6,56 | <0.0001 | -3 | 1.4 | 56 | 0.17 | 0.92 |
| | treatment | 12.35 | 1,56 | 0.001 | 1 | 1.9 | 56 | 0.058 | 0.89 |
| | day:treatment | 3.41 | 6,56 | 0.007 | 8 | 4.1 | 56 | <0.0001 | 0.73 |
| | | | | | 16 | -0.5 | 56 | 0.59 | 1.04 |
| | | | | | 22 | -1.4 | 56 | 0.17 | 1.1 |
| | | | | | 29 | 1.3 | 56 | 0.22 | 0.91 |
| | | | | | 36 | 2.6 | 56 | 0.013 | 0.84 |
